# Manifold Agentic Reasoning: Extending Agentic POMDPs and Post-Training Reasoning to Riemannian State and Reasoning Spaces

**DOI:** 10.64898/2026.07.26.740848

**Authors:** Tao Xu, Zixin Hu, Xiaodian Sun, Li Jin, Momiao Xiong

## Abstract

Agentic reasoning systems increasingly interact with environments whose states are only partially observed, dynamically evolving, and constrained by physical, biological, or logical structure. Existing agentic reasoning frameworks often model internal reasoning, tool use, and post-training adaptation using flat latent representations and struggle in curved manifold space environments. However, many scientific and embodied domains naturally lie on curved state spaces, including tissue geometry, developmental trajectories, protein conformations, robotic configuration spaces, and constrained physical systems. We introduce Manifold Agentic Reasoning, a geometric framework that extends agentic reasoning from Euclidean latent spaces to Riemannian manifolds. In the proposed framework, observations are encoded as manifold-valued states, memory is retrieved by geodesic similarity, candidate hypotheses are generated in tangent spaces, predicted transitions are projected by exponential maps, and decisions are admitted through verification-gated commitment or repaired by manifold self-correction. We further extend the framework to graph-agentic manifold reasoning, where node states live on manifolds and neighbor information is transported by logarithmic maps before attention-based aggregation. Manifold agent reasoning moves AI past brittle, prompt-chained templates to solve four critical production flaws: silent hallucinations and reasoning drift, brittle tool and context misuse, the black-box evaluation problem and stiff behavior profiles.

To evaluate the framework, we introduce a Curved Tissue Manipulation and Recovery benchmark in which an agent must repair damaged tissue on a curved manifold. Simulated results show that the full manifold-agent substantially outperforms both a baseline reasoning agent and a full flat-agent reasoning system, achieving higher recovery success, lower geodesic shape error, lower pattern error, and fewer invalid transitions. Curvature and ablation studies indicate that the performance gain is driven by geometry-aware reasoning, verification, memory, and self-repair. These results suggest that manifold-aware agentic reasoning provides a principled foundation for reliable scientific and embodied AI systems operating in curved, constrained, or mechanistic domains.

## 1. Introduction

Agentic reasoning has achieved remarkable gains and marks a fundamental shift in artificial intelligence, transitioning models from passive, “one-shot” text generators into autonomous, goal-directed problem solvers. Rather than predicting the next word in a single breath, systems utilizing agentic reasoning employ an ongoing **plan-act-observe-reflect loop** to accomplish highly complex tasks over extended periods. The field has advanced past simple trial-and-error scripts into a sophisticated discipline centered on efficiency, standardized protocols, and architectural scaling (Zhao et al. 2025).

Most agentic reasoning is flat reasoning that assumes data follows Euclidean geometry. The Euclidean geometry has three remarkable properties: (1) linear Relationships: it assumes the shortest path between two points is a straight line; (2) grid Structure: it views variables as independent, standard grid coordinates (like *X* and *Y*); and (3) limitations: it fails when data features interact in complex, non-linear ways.

While they often visualize this space as a flat vector grid (Euclidean space), AI agents need more complex reasoning shapes to handle complex data structures. Many biological, physical, cognitive, and robotic systems are not naturally Euclidean. Examples include: cell states (Wang et al. 2026), RNA-velocity trajectories (Dunican et al. 2026), tissue morphologies (Autorino et al. 2026), robot poses (Liu et al. 2025), protein conformations (Cho et al. 2026), hypothesis spaces (Tiblias et al. 2025). Concretely, a manifold is defined as a space that looks flat up close but is curved globally (like Earth). Real-world data (language, physics, biology, human behavior) sits on these curved shapes. The best path between two points on a manifold is a curve, not a straight line. The agent changes its decision-making rules depending on where it is in the data space. When an AI agent uses “flat” logic on “curved” data, it makes bad predictions because its mental map does not match the actual shape of the problem.

The current approach to manifold agentic reasoning represents a paradigm shift in how artificial intelligence structures, monitors, and optimizes multi-step problem solving. Rather than analyzing an LLM statically based on training data or treated as a simple token predictor, this framework maps the high-dimensional internal trajectories of a model’s hidden states during execution into a continuous mathematical landscape known as the Reasoning Manifold Developing manifold agentic reasoning requires shifting AI architecture away from rigid, linear processing and toward methods that map, navigate, and steer across the curved semantic topologies of high-dimensional data (ΔΦ Nexus 2025; Chun et al. 2025).

There are several published papers that directly investigate “manifold reasoning” and how it interfaces with “agentic AI” and large reasoning models. Rather than treating agentic reasoning purely as text-based chains-of-thought, these papers explore it from a geometric and topological perspective—viewing an artificial agent’s internal states, thinking paths, and tool-interaction boundaries as low-dimensional manifolds within high-dimensional activation spaces (Cao et al. 2026). Key published papers and frameworks in this domain include:

Paper 1: “REMA: A Unified Reasoning Manifold Framework for Interpreting Large Language Model (Li et al.2025)”. This paper formally defines a Reasoning Manifold as a latent, low-dimensional geometric structure formed by the internal representations of all correctly reasoned model. It maps out the “effective thinking paths” learned by models to solve tasks. By tracking when an AI agent drifts off this manifold, researchers can quantitatively diagnose exactly where an agentic workflow or reasoning sequence failed.

Paper 2: “Mitigating Overthinking in Large Reasoning Models via Manifold Steering (Huang et al. 2025)”. Large reasoning models often suffer from “overthinking” (generating unnecessary internal text tokens). This paper proposes projecting steering directions onto a low-dimensional manifold. By constraining the reasoning path to this manifold, it eliminates high-dimensional interference noise. It successfully reduces output tokens by up to 71% without sacrificing the agent’s accuracy on complex tasks.

Paper 3: “Characterizing AlphaEarth Embedding Geometry for Agentic Environmental Reasoning (Rahman et al. 2026), This study investigates the manifold geometry of geospatial foundational embeddings. It explicitly builds an agentic system that navigates non-Euclidean manifold structures to execute advanced environmental and spatial reasoning.

Paper 4: “Paper: Emergent Manifold Separability during Reasoning in Large Language Models (Chun et al. 2026) This paper proposed applies manifold capacity theory (MCT) to evaluate the temporal dynamics of an AI model’s internal geometry while solving complex, compositional logic puzzles.

Existing agentic reasoning work models agents as planning-and-acting systems, often using POMDP-like abstractions and latent reasoning variables. Existing geometric RL and manifold learning work models curved state spaces. However, a unified framework that treats both the environment state and the internal reasoning process as manifold-valued objects optimized through post-training reinforcement learning remains underdeveloped. Therefore, we propose MAR / MAP-R to fill this gap.

The key four contributions are as follows:

1. Proposes Manifold Agentic Reasoning, a general framework for agentic reasoning on Riemannian state and reasoning spaces and fully develop a Manifold-driven Agentic Architecture..
2. Extends the POMDP and internal reasoning variable formulation to manifold-valued world states, reasoning states, beliefs, and policies.
3. Creates a shared geometric manifold where multiple autonomous agents can track, intersect, and merge their respective reasoning paths during long-horizon tasks and introduces a full algorithmic pipeline with tangent-space hypothesis generation, exponential-map projection, verification-gated commitment, and manifold self-repair.
4. Demonstrates the framework on a curved tissue manipulation and recovery benchmark, showing improved recovery success, lower invalid transition rate, and robustness to curvature.

## 2. MAR: Manifold Agentic Reasoning

We first present a mathematical framework for extending agent reasoning from Euclidean space to manifold space. We will call the framework Manifold Agentic Reasoning, or MAR.

### 2.1. Euclidean agent reasoning

Euclidean agent reasoning refers to a framework where an artificial intelligence or autonomous agent models its environment, decisions, and goals using geometry—specifically, Euclidean space (Liang 2026). Instead of using purely logical rules, the agent treats concepts, states, and relationships as points and distances in a multi-dimensional coordinate system. Here is a breakdown of how Euclidean agent reasoning works, its core components, and its applications. A standard Euclidean agent can be written as

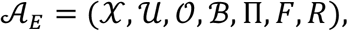

where:

*X* ⊆ ℝ^*d*^ is the state space, *U* ⊆ ℝ^*m*^ is the action space, *o*_*t*_ ∈ *O* is the observation, *b*_*t*_ ∈ ℬ is the belief state, *π*_*θ*_(*u*_*t*_ ∣ *b*_*t*_) is the policy, *x*_*t*+1_ = *F*_*θ*_(*x*_*t*_, *u*_*t*_, *e*_*t*_) is the transition model, an *R*(*x*_*t*_, *u*_*t*_, *x*_*t*+1_) is the reward or utility.

The agent translates complex real-world data (words, images, or sensory inputs) into numerical vectors. These vectors exist as coordinates in a high-dimensional Euclidean space where similar concepts are clustered close together. The agent calculates the straight-line distance (Euclidean distance) between points to determine similarity or relevance. Every possible situation or environment snapshot is represented as a point *P* = (*x*_1_, *x*_2_, ⋯, *x*_*d*_). The agent’s objective is also a point in the same space. The agent continuously uses the standard Euclidean distance formula to measure how far it is from its goal:

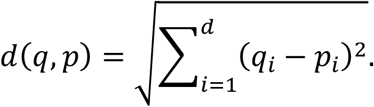

Actions are vectors that shift the agent’s current position closer to the goal coordinates. The reasoning loop is:

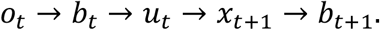

In a flat space, update rules often use vector addition:

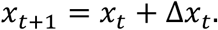

But on a curved manifold, this is generally invalid because *x*_*t*_ and Δ*x*_*t*_ do not live in the same space. Instead, updates must occur in the tangent space and then be mapped back to the manifold.

### 2.2. Basic idea of extension from Euclidean space to manifold space

Classical agent reasoning often assumes that data, states, actions, memories, and hypotheses live in a flat Euclidean space:

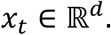

The agent observes a state *x*_*t*_, reasons, chooses an action *u*_*t*_, updates its belief, and predicts the next state:

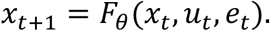

This is appropriate when the data geometry is approximately flat. However, many biological, physical, cognitive, and robotic systems are not naturally Euclidean. Their states may lie on curved spaces:

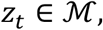

where ℳ is a differentiable manifold.

Examples include:

cell states,RNA-velocity trajectories,tissue morphologies,robot poses,protein conformations,hypothesis spaces.

The core idea is to replace flat reasoning ℝ^*d*^ by geometric reasoning on a manifold

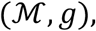

where *g* is a Riemannian metric that defines local distance, angle, curvature, geodesics, and energy.

The Euclidean case is recovered when

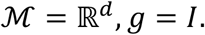

So manifold reasoning is a strict generalization of Euclidean reasoning.

### 2.3. Manifold agent reasoning

The following formulation shifts agent reasoning from flat, straight-line geometry (Euclidean space) to curved, restricted surfaces (manifolds). It models an agent that must navigate and make decisions while strictly adhering to the complex, underlying shape of its environment.

#### 2.3.1. The Environment Geometry (*M, T*_*z*_*M, M, g*)

Instead of moving freely in any direction, the agent is bound to a specific “shape.” Let the true state space be a smooth manifold: ℳ, which is a smooth, continuous surface (like the surface of a sphere, a donut, or a complex physical constraint). The agent can only exist *on* this manifold.

At each state *z* ∈ ℳ, the tangent space is

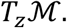

Imagine placing a flat sheet of paper perfectly balanced on one point (*z*) of a globe. That flat paper is the tangent space. It represents all the possible directions and velocities the agent can instantly move from that exact spot.

A Riemannian metric *g*_*z*_ is a smoothly varying inner product:

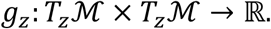

The Riemannian metric *g*_*z*_ is a tool that acts like a local, flexible ruler (Gruffaz and Sassen 2025). Because the surface is curved, a standard straight ruler doesn’t work. *g*_*z*_ takes two velocity vectors in the tangent space and calculates their inner product, allowing the agent to measure actual lengths, angles, and distances along the curved surface.

It defines the local norm:

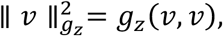

implying the squared speed or magnitude of a movement vector *v* at state *z*, adjusted for the curvature of that exact location.

#### 2.3.2. The Manifold Agent Tuple (*A*_*M*_)

The tuple defines the complete operating system of the agent, adapted for a curved universe:

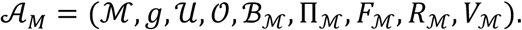

Here:

*z*_*t*_ ∈ ℳ is the latent manifold state, i.e., the agent’s actual, hidden position on the curved surface at time *t*,

*u*_*t*_ ∈ *U* is the action, including the decisions, forces, or steering inputs available to the agent, *b*_*t*_ ∈ ℬ_ℳ_ is the belief over manifold states, representing the agent’s internal probability distribution or “best guess” of where it is on the manifold, since it cannot observe \(z_{t}\) perfectly,

*π*_*θ*_(*u*_*t*_ ∣ *b*_*t*_, *z*_*t*_) is a policy (strategy) function. It determines the next action based on what the agent believes (*b*_*t*_) and where it actually is (*z*_*t*_).

#### 2.3.3. Dynamics and Constraints (*F*_ℳ_, *V*_ℳ_)

This is where the manifold structure forces the agent to behave differently than a standard AI. *F*_ℳ_: ℳ × *U* × ℰ → ℳ is the manifold transition model, the physics engine. It takes the current curved state ℳ, the action *U*, and environmental noise ℰ, and outputs a new state that is guaranteed to still rest on the manifold ℳ. It prevents the agent from flying off the surface into empty space.

*V*_ℳ_(*z*_*t*_, *u*_*t*_, *z*_*t*+1_) is a verifier that checks whether a transition is geometrically, causally, and mechanistically valid. In other words, it ensures the movement obeys three strict rules: Geometrically: Did the path stay perfectly on the manifold without cutting corners through “empty air”?

Causally: Did the action cause the effect, or is it a temporal paradox?

Mechanistically: Did the transition obey the laws of physics and the agent’s internal mechanics? **In summary**, in Euclidean reasoning, an AI calculates the shortest path between two points as a straight line. If an AI is controlling a robotic arm, a straight line might command the arm to slice directly through a metal table.

By using Manifold Agent Reasoning, the “table” is carved out of the state space. The metric *g*_*z*_ forces the agent to calculate geodesics (the shortest paths around constraints), and the verifier *V*_ℳ_ ensures the robot smoothly glides along allowed physical boundaries.

Figure 1 shows information flow in manifold agent reasoning.

**Figure 1.**
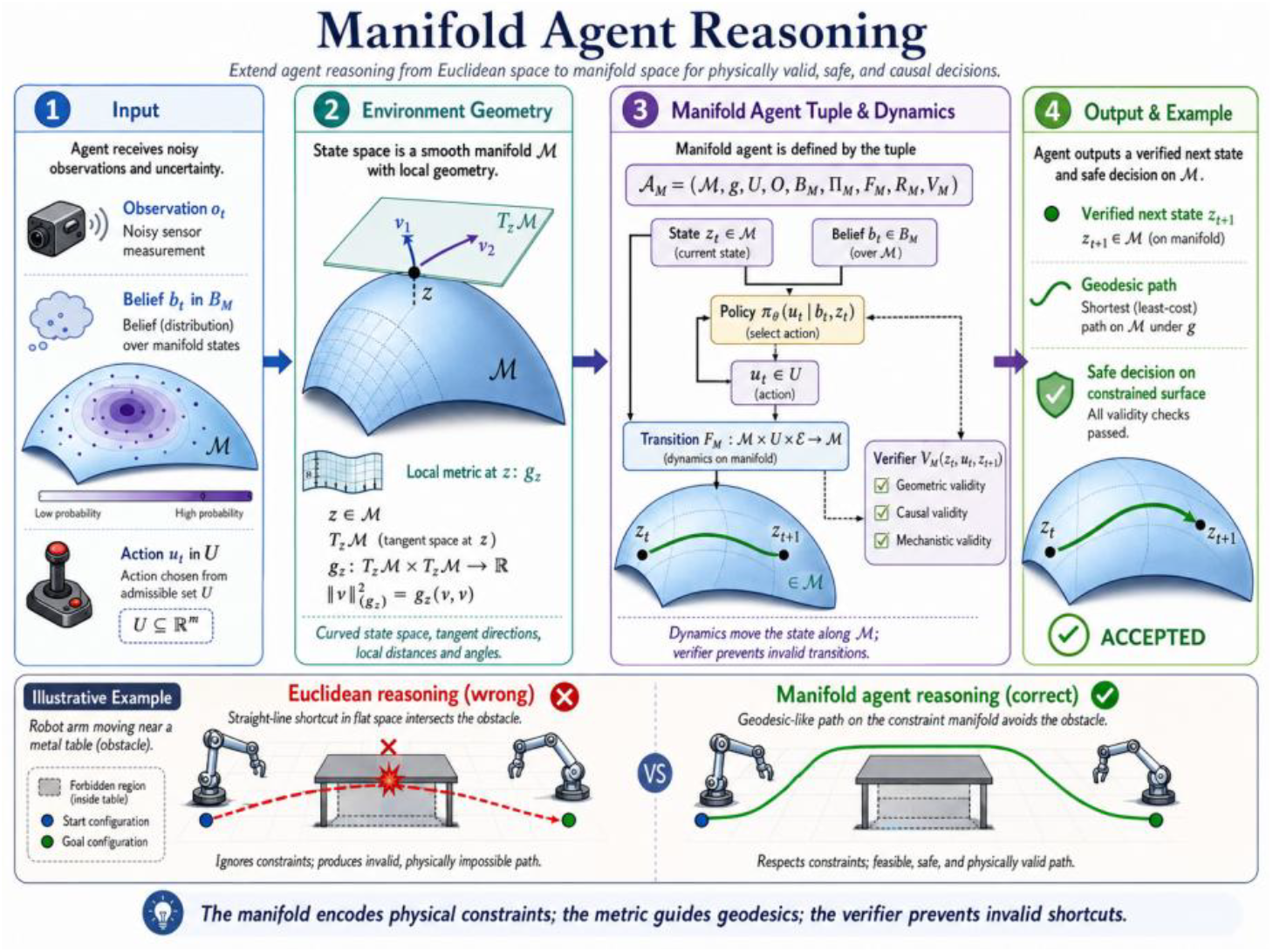
Manifold agent reasoning.

### 2.4. Local charts and coordinates

#### 2.4.1. The Core Concept: Charts and Decoders

Local charts and coordinates are essential in manifold-based AI agent reasoning because a manifold cannot be mapped globally by a single coordinate system without causing severe geometric distortion or mathematical contradictions.

Just as a flat paper map cannot represent the entire round Earth without tearing or stretching the poles, complex data spaces (like robot joint configurations, 3D rotations, or deep learning feature spaces) cannot be modeled using a single flat coordinate system. Local charts allow an AI agent to break a complex, curved space into a collection of overlapping, flat, and easily calculable pieces.

The following mathematical framework describes how a machine learning agent can model a complex, curved environment (a manifold) by sewing together a collection of flat, local coordinate systems. It is the mathematical foundation for Manifold agent reasoning.

Because a manifold may not have one global coordinate system, we use local charts. Instead of mapping everything at once, we look at a small, localized region (an open set) on the manifold, called *U*_*l*_ (patch).

A chart is

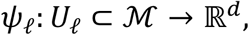

with inverse decoder

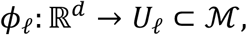

where *ψ*_*ℓ*_ is called the Chart/Encoder and *ϕ*_*ℓ*_ is called the Decoder. *ψ*_*ℓ*_ is a function that takes a true manifold state *z*_*t*_ inside that local patch and flattens it out into a *d*-dimensional vector space (ℝ^*d*^). This 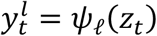 is your local coordinate, a simple list of numbers an AI agent can easily use for calculation.

*ϕ*_*ℓ*_ is the inverse function. It takes the flat numbers 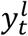 and “inflates” or projects them back up onto the actual curved manifold surface to reconstruct the state

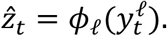

#### 2.4.2. Preserving the True Geometry

When encoding and decoding, the agent must not warp distances or shapes. This loss function forces the neural networks to preserve geometry using two terms:

##### 1. Reconstruction Error *d*_ℳ_(*z*_*t*_, *ϕ*_*ℓ*_(*ψ*_*ℓ*_(*z*_*t*_)))^2^

If you take a manifold state, flatten it into local coordinates, and immediately decode it back, you should end up exactly where you started.

Instead of standard Euclidean distance, it uses geodesic distance *d*_ℳ_, which measures the shortest path along the curved surface of the manifold.

##### 2. Metric Matching Error 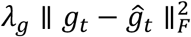

A Riemannian metric (*g*) is a mathematical tool that tells you how to calculate distances, angles, and volumes at any specific point on the manifold. This term forces the agent’s learned metric (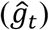) to match the environment’s true metric 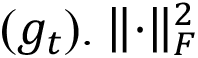 is the Frobenius norm, which is just a way to measure the error between two matrices.

*λ*_*g*_ is a scaling weight.

A geometry-preserving loss is

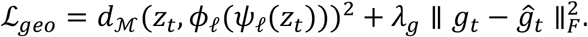

Here *d*_ℳ_ is geodesic distance, and ĝ_*t*_ is the learned metric induced by the chart.

##### 3. The Induced Metric and the Jacobian

How does a flat coordinate system understand a curved surface? It uses the Jacobian 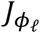.

The induced metric in local coordinates is

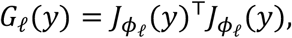

where 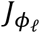 is the Jacobian of the decoder.

###### The Jacobian

This is a matrix of all first-order partial derivatives of the decoder function. It represents the best linear approximation of how the decoder stretches or squishes the space at coordinate *y*. It maps a tiny step in your flat coordinates to a tiny step on the curved manifold.

###### The Metric *G*_*ℓ*_(*y*)

By multiplying the Jacobian by its transpose *J*^*T*^*J*, you get a matrix called the pullback metric. If an agent wants to know the true distance of a small path in its flat coordinate system, it multiplies the path by*G*_*ℓ*_(*y*). It translates “flat steps” into “curved distances.”

##### 4. Overlap and Transition Maps (The Atlas)

As an AI agent moves through a large environment, it will step out of the boundaries of patch *U*_*l*_ and into an adjacent patch *U*_*k*_.

###### The Transition Map

When patch *L* and patch *K* overlap, there must be a seamless way to convert coordinates from Chart *L* directly to Chart *K*. You do this by decoding from *L* up to the manifold *ϕ*_*ℓ*_, and then immediately encoding down into *K*(*ψ*_*k*_). This combined operation *ψ*_*k*_ ∘ *ϕ*_*ℓ*_ is the transition map.

If two charts overlap, they must be consistent:

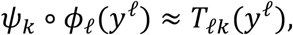

where

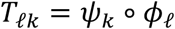

is the transition map between charts.

###### The Overlap Loss ℒ_*overlap*_

This ensures that the chart mappings don’t contradict each other where they meet.

The overlap loss is

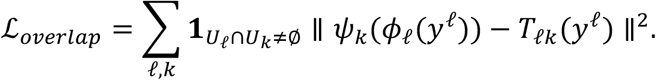

The indicator function 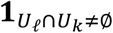 ensures we only calculate this loss for patches that actually touch or overlap. The norm term (‖⋅‖^2^) penalizes the neural networks if converting coordinates via the manifold disagrees with the learned transition mapping.

Without this exact formulation, an agent traveling across a curved space would experience sudden “jumps” in its coordinate data, or its path planning gradients would explode at boundaries. This framework ensures that no matter where the agent travels, it always has a smooth, flat, mathematically accurate notebook (local chart) to calculate its next move.

**Figure 2.**
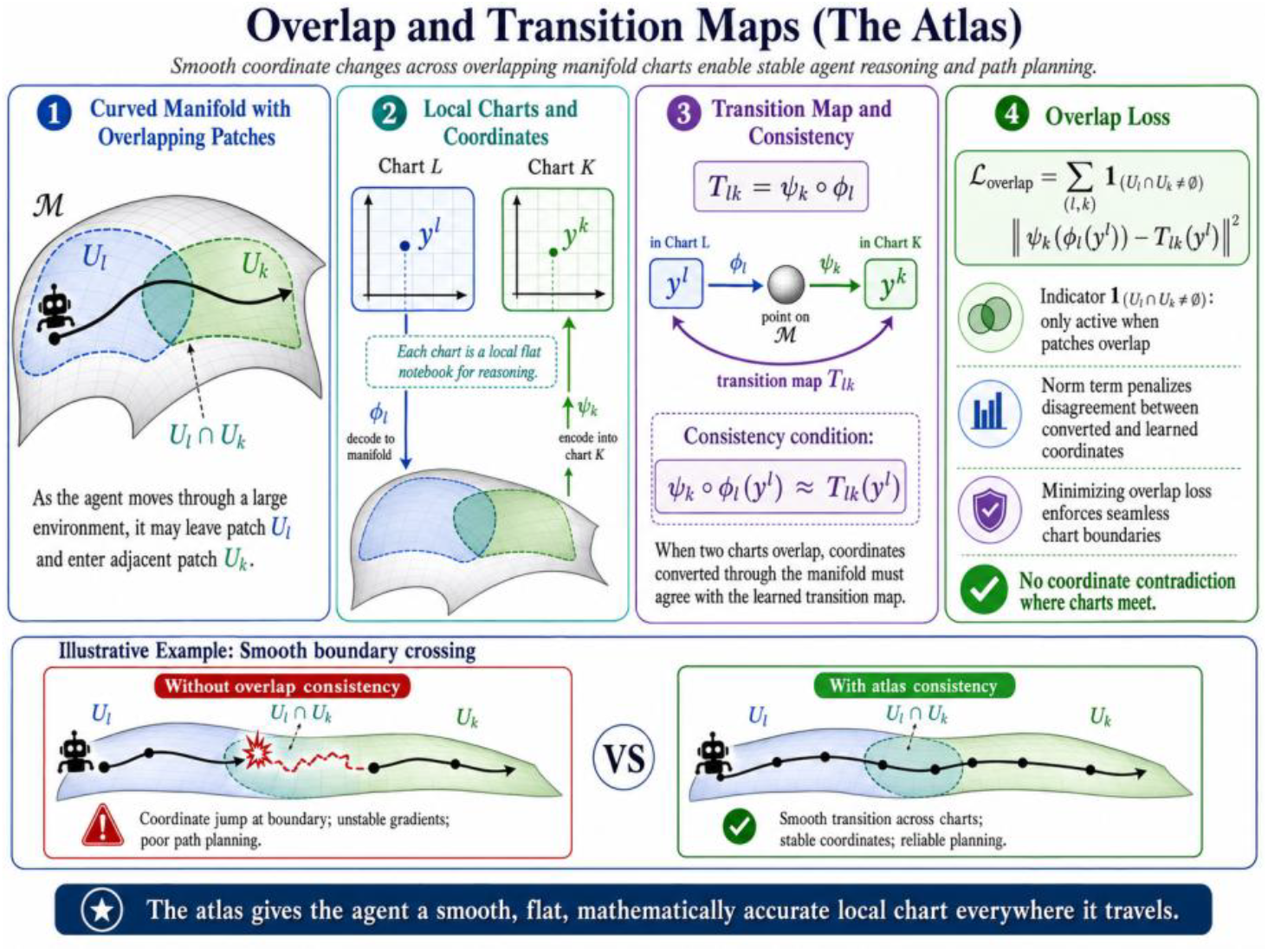
Overlap and transition maps.

### 2.5 Manifold state transition

In manifold agentic reasoning, a manifold transition refers to the mathematical and structural movement of an AI agent’s internal state from one region of its latent “reasoning manifold” to another as it processes a task. Instead of viewing an AI’s thinking as a simple linear chain of text tokens, geometric interpretability frameworks—such as the Unified Theory of Agentic Reasoning and REMA (Reasoning Manifold Framework)—map the agent’s hidden activation states as points moving through a highly structured, low-dimensional curved space (a manifold). When an agent executes an action, reflects, or calls an external tool, its internal mathematical state shifts. This transition occurs across several specialized topological structures within the reasoning manifold.

#### 2.5.1. Euclidean Space vs. Curved Manifold

The formula for manifold state transition defines how an AI agent’s internal state moves along a curved mathematical surface (the reasoning manifold ℳ) instead of a flat, linear space

In Euclidean space, one writes

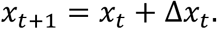

The Problem is that if you do this on a curved surface, adding a straight arrow makes the state fly off the surface into invalid space. To stay perfectly on the curved reasoning surface, the model uses an exponential map:

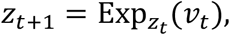

where 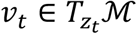 is a tangent vector. It takes a straight velocity vector *v*_*t*_ sitting in the flat tangent space 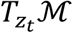 touching the manifold at point *z*_*t*_ and “bends” or walks it along the curved surface to land exactly at the next valid state *z*_*t*+1_.

#### 2.5.2. The Tangent Dynamics Model

The agent learns a tangent dynamics model. The formula defines the velocity vector *v*_*t*_ as a function of the agent’s variables:

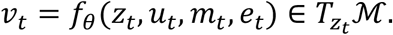

This means a neural network (parameterized by θ) computes the direction and speed of the reasoning shift, based on four core inputs:

*z*_*t*_: the agent’s current internal reasoning state,

*u*_*t*_: the next token, choice, or tool action taken by the agent,

*m*_*t*_: the working memory or hidden context window,

*e*_*t*_: external environmental feedback or retrieved tool results.

Substituting this into the map gives the complete discrete update equation:

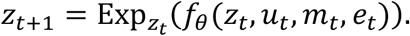

#### 2.5.3. Coordinate Charts and Lifting

Because calculating full exponential maps directly can be computationally heavy, the formula introduces a practical workaround using local coordinates (charts):

##### Flattening

A local chart maps a patch of the curved manifold to a flat grid of local coordinates *y*_*t*_.

##### Euclidean Step

The model takes a standard linear step in this flat grid:

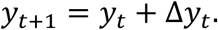

##### Lifting

The function *ϕ*_*ℓ*_(*y*_*t*+1_)takes that flat coordinate and lifts it back onto the actual curved manifold surface to find

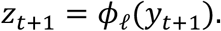

#### 2.5.4. Continuous-Time Geometric Flow

A more geometric version is

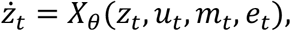

where

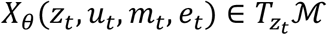

is a learned vector field.

Instead of choppy step-by-step updates (*t* → *t* + 1), this version treats reasoning as a smooth, continuous fluid flow over time (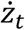 represents the instantaneous derivative or velocity). *X*_*θ*_ is a parameterized vector field that defines a continuous current guiding the agent’s thought trajectory seamlessly along the low-dimensional highway of correct logic.

The continuous-time transition is

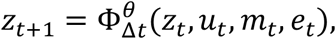

where 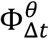 is the flow map generated by *X*_*θ*_.

Figure 3 visualizes manifold state transition.

**Figure 3.**
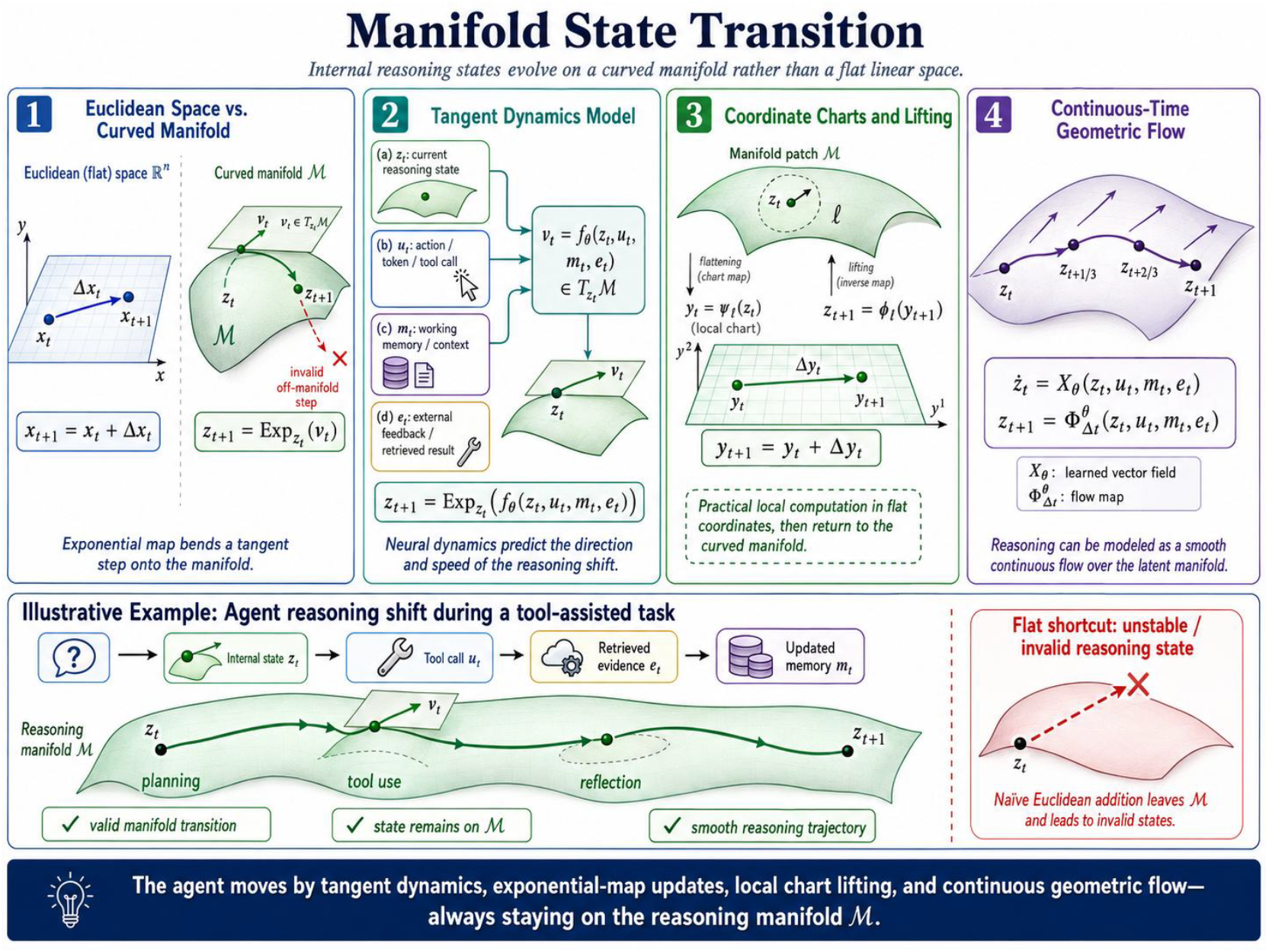
Manifold state transition.

### 2.6 Manifold belief state

A Manifold Belief State is the geometric representation of an AI agent’s internal knowledge, uncertainty, and hypotheses about the world, mathematically constrained to a low-dimensional curved subspace (a manifold). Instead of viewing an agent’s “belief” as a disjointed list of facts, advanced geometric frameworks—such as the Semantic Belief-State World Model (SBWM) and recent studies on Belief Manifolds—treat beliefs as continuous, structured objects with distinct geometric properties like coordinates, mass, and inertia.

The following mathematical formula explains how an AI agent shifts its understanding of reality (its belief state) when tracking a problem along a curved mathematical surface (the manifold ℳ) rather than a flat grid. It generalizes standard Bayesian filtering—like a Kalman filter—into Riemannian geometry.

#### 2.6.1. Shift from Flat Space to Manifold Space

##### Euclidean (Flat Space)

In Euclidean reasoning, belief is often represented as a distribution over ℝ^*d*^:

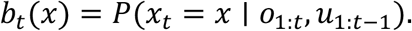

In a typical AI model, a belief state *b*_*t*_(*x*) is simply a probability distribution over flat coordinate ℝ^*d*^. It estimates where the world state *o*_1:*t*_ and actions *u*_1:*t*−1_.

##### Manifold (Curved Space)

In manifold reasoning, belief is a probability measure over ℳ:

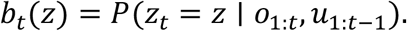

Here, the belief is modeled as a probability measure over a curved space ℳ. The hidden state *z*_*t*_ is mathematically forced to stay strictly on the surface of valid reasoning trajectories.

#### 2.6.2. The Manifold Bayesian Update Rule

The belief update becomes

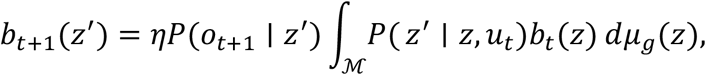

where *dμ*_*g*_(*z*) is the Riemannian volume measure and *η* is a normalization constant. This is a standard hidden Markov model prediction-correction loop, adapted for curved geometry:

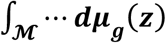: Instead of integrating over flat space, it sums up probabilities over the manifold using the Riemannian volume measure (*dμ*_*g*_(*z*)), which accurately compensates for the stretching or squeezing of the curved surface.

***P***(***z***′|***z, u***_***t***_): The transition probability of moving from old state *z* to new state *z*′ given action *u*_*t*_. ***P***(***o***_***t***+**1**_|***z***′): The likelihood of receiving the newest observation *o*_*t*+1_ given the new proposed state *z*′.

***η*:** A normalization constant ensuring all probabilities sum to exactly 1.

#### 2.6.3. Transition Density with Manifold Gaussian Noise

To calculate the path likelihood, standard flat systems use a clean bell curve (Gaussian noise). On a manifold, straight lines do not exist, so the formula redefines the transition density using geodesics (the shortest paths along curved surfaces):

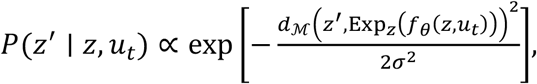

where

**Exp**_***z***_(***f***_***θ***_(***z, u***_***t***_)): The ideal, noise-free spot where the agent’s internal dynamics model *f*_*θ*_ predicts it should land after taking action *u*_*t*_,

***d***_**ℳ**_(⋯): The geodesic distance on the manifold between the actual landing point *z*′ and that predicted ideal spot,

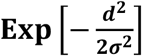: This creates a Riemannian Gaussian distribution (heat kernel approximation). If the agent drifts too far from the predicted reasoning path, the probability drops exponentially, serving as an indicator of geometric deviation or an impending hallucination.

This replaces Euclidean Gaussian transition noise with manifold Gaussian-like noise. Figure 4 plots transition Manifold Gaussian density function.

**Figure 4.**
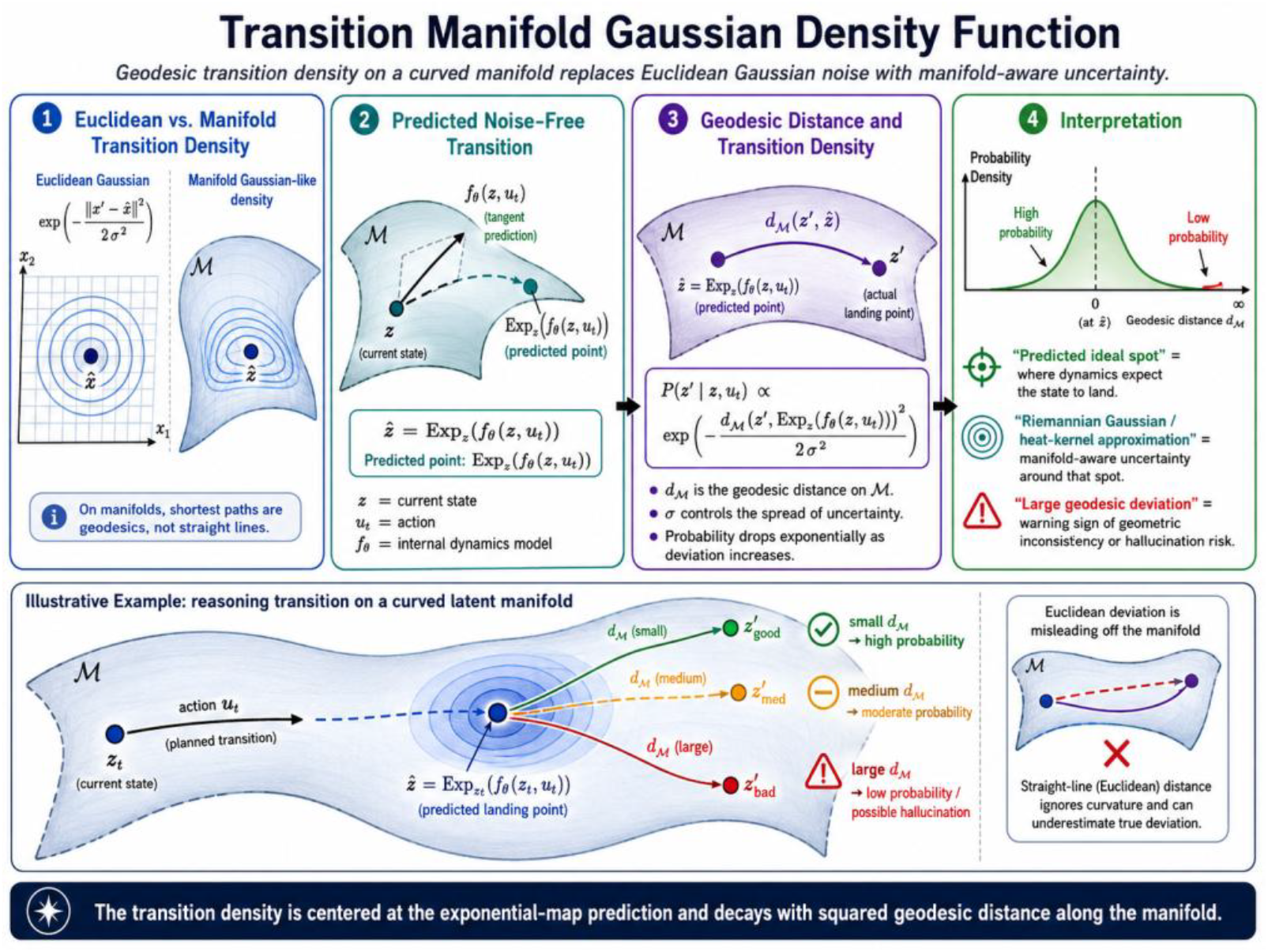
Transition Manifold Gaussian density function.

### 2.7. Manifold policy

The below mathematical formulas define how an AI agent chooses actions and moves when operating directly on a curved geometric surface (a Riemannian manifold), rather than in flat, standard Euclidean space.

#### 2.7.1. The Core Policy Function

A policy maps a belief and state to an action:

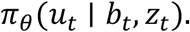

##### What it means

This is the agent’s decision-making rule (policy), parameterized by neural network weights *θ*.

##### The components

It outputs the probability of taking a specific action *u*_*t*_ given the agent’s current hidden belief state *b*_*t*_ (its internal memory or history) and its physical geometric state *z*_*t*_ (its current position on the manifold ℳ.

#### 2.7.2. Tangent Space Action Constraints

If the action itself lives in a tangent space, then

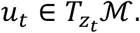

Then the policy can be written as

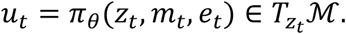

##### What it means

You cannot easily apply a flat, straight-line vector to move on a curved surface like a sphere. Instead, the agent’s action *u*_*t*_ is chosen from the tangent space 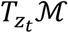.

##### Analogy

Think of the manifold as the Earth. The tangent space is a flat sheet of paper touching the Earth perfectly at your exact current location *z*_*t*_. The action *u*_*t*_ is a straight arrow drawn on that flat paper, representing your velocity vector (speed and direction).

##### The inputs

The policy constructs this vector using the current position *z*_*t*_, internal context/memory *m*_*t*_, and environmental inputs or random noise *e*_*t*_.

#### 2.7.3. Moving on the Manifold (The Exponential Map)

The next state is

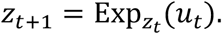

##### What it means

This formula updates the agent’s position from time step *t* to *t* + 1. Since the action vector *u*_*t*_ is flat, you cannot just add it to *z*_*t*_ (*z*_*t*_ + *u*_*t*_ would shoot off into outer space, leaving the surface of the Earth).

##### The mechanism

The Exponential Map 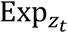 takes that flat velocity vector *u*_*t*_, bends it down tightly against the curved surface, and snaps the agent along a “straight line” on the curved surface (a geodesic).

#### 2.7.4. Random Actions on Curved Surfaces (Stochastic Policies)

For stochastic policies:

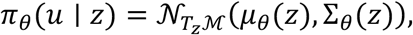

redictable, where the Gaussian is defined in the tangent space *T*_*z*_ℳ.

##### What it means

For an agent to explore effectively, its actions often need to be random (stochastic) rather than perfectly predictable. However, defining a standard bell curve (Gaussian/Normal distribution) directly on a curved surface is mathematically messy.

##### The mechanism

This formula solves that by defining a standard Gaussian distribution *N* inside the flat tangent space *T*_*z*_ℳ.

##### The components

The policy network outputs a mean action vector *μ*_*θ*_(*z*) (the direction the agent *thinks* it should go) and a covariance matrix Σ_*θ*_(*z*) (how much uncertainty or random wiggle room it has). Because the tangent space is completely flat, standard linear algebra and neural networks can calculate this distribution without breaking. The resulting random vector is then projected back down to the manifold using the Exponential map described in 2.7.3.

##### Summary of Agentic Flow

In manifold agentic reasoning, every single decision loop follows this precise geometric sequence:

##### Perceive

The agent evaluates its current position *z*_*t*_ on the curved world map ℳ.

##### Plan

The neural network calculates a normal distribution directly inside a flat localized sheet *T*_*z*_ℳ resting on that point.

##### Sample

The agent samples a random directional velocity vector *u*_*t*_ from that flat distribution. **Act:** The Exponential Map wraps that flat vector onto the curved geometry, smoothly sliding the agent to its true next physical state *z*_*t*+1_.

### 2.8. Manifold reward and value function

In standard agent reasoning, agents assume the world is flat, measuring success with simple straight-line distances. In manifold agentic reasoning, the reward is no longer simply a function of Euclidean distance. The environment is a curved geometric surface ℳ. The reward can depend on geodesic distance, curvature, energy, causal validity, and biological plausibility. The math for reward calculation defines how an agent evaluates its success (the Reward Function), judges long-term prospects (the Value Function), and calculates optimal choices (the Manifold Bellman Equation) under curved geometry.

#### 2.8.1. The Manifold Reward Function

Define

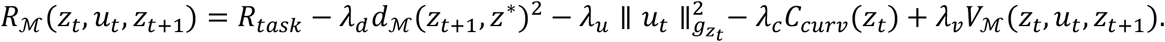

Here:

*z*^∗^ is the target state, 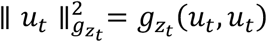 is the geometric action cost *C*_*curv*_ (*z*_*t*_) is a curvature penalty, and *V*_ℳ_ is a verification score.

Instead of just checking if it reached a goal, the agent’s reward balances task completion against structural and geometric rules. The terms in *R*_ℳ_(*z*_*t*_, *u*_*t*_, *z*_*t*+1_) are explained as follows.

***R***_***task***_ **(Task Reward):** The base reward for accomplishing the primary goal (e.g., finding a target, solving a problem).

−***λ***_***d***_***d***_**ℳ**_(***z***_***t***+**1**_, ***z***^∗^)^**2**^ **(Geodesic Distance Penalty):** Penalizes the agent based on how far it is from the target state *z*^∗^. Crucially, *d*_ℳ_ is the geodesic distance (the shortest path along the curved surface), not a straight Euclidean line through empty space.

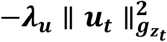 **(Geometric Action Cost):** Penalizes excessive effort or velocity. Because the action *u*_*t*_ lives in the tangent space, its magnitude must be measured using the Riemannian metric tensor 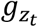. This functions like a coordinate-aware scaling matrix that calculates true physical energy spent based on the agent’s current location.

−***λ***_***c***_***C***_***curv***_(***z***_***t***_) **(Curvature Penalty):** Discourages the agent from entering regions of the manifold that are highly warped, unstable, or geometrically complex (e.g., avoiding chaotic “crumpled” areas of data).

***λ***_***v***_***V***_**ℳ**_(***z***_***t***_, ***u***_***t***_, ***z***_***t***+**1**_) **(Verification Score):** Rewards actions that maintain structural properties like causal validity or biological plausibility. It keeps the agent’s path aligned with real-world logic or constraints.

#### 2.8.2. The Long-Term Value Function

The value function is

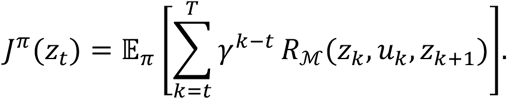

##### What it means

This measures the total expected quality of a position. It is the sum of all future geometric rewards the agent expects to collect from time *t* until the end of the task *T*.: The ***γ***^***k***−***t***^: The discount factor (between 0 and 1) that makes immediate rewards more valuable than distant future rewards.

#### 2.8.3. The General Manifold Bellman Equation

The optimal value satisfies the manifold Bellman equation:

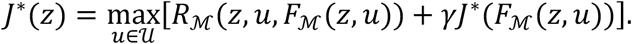

##### What it means

This states that the value of the current state *z* under an optimal policy *J*^∗^ equals the immediate reward of the best possible action plus the discounted value of the next state.

***F***_**ℳ**_(***z, u***): This represents the environment’s transition physics—how state *z* changes to the next state given action *u* on the manifold.

#### 2.8.4. Exponential-Map Dynamics (The Geometric Reality)

With exponential-map dynamics:

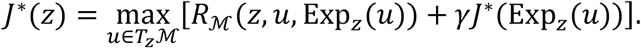

This formula swaps the abstract physics engine *F*_ℳ_(*z, u*) for concrete differential geometry. **The Action Space *u*** ∈ ***T***_***z***_**ℳ:** The agent optimization search (max) happens strictly over flat vectors inside the local tangent space ***T***_***z***_**ℳ**.

##### The State Transition (Exp_*z*_(*u*))

The next state is explicitly defined by the Exponential Map. The agent shoots a velocity vector *u* out into its flat local tangent space, and the geometry automatically bends that path back onto the true surface to find the exact landing coordinates.

### 2.9. Hamiltonian formulation for manifold reasoning

A generalized Hamiltonian framework is designed to model systems where physical laws, geometry, and internal regulations interact. This formulation forces a machine learning model or mathematical simulator to respect energy conservation and geometric constraints while tracking changing internal systems. For mechanistic systems, especially biological dynamics, robotics, and tissue morphogenesis, we may use a Hamiltonian structure.

#### 2.9.1. Breakdown of the State Variables

Let

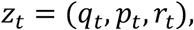

where: *q*_*t*_ ∈ ℳ is a configuration state, 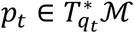 is momentum, and *r*_*t*_ is a mechanism or regulatory state.

The cotangent bundle is

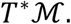

The full state of the system is captured by the vector *z*_*t*_ = (*q*_*t*_, *p*_*t*_, *r*_*t*_). This divides the system into geometric physics and non-physical internal adjustments:

##### Configuration State (q_t_ ∈ ℳ)

The physical position, shape, or structural layout of the system. It lives on a smooth mathematical space called a manifold (ℳ), which enforces structural constraints (e.g., a rigid robot joint or a fixed-volume biological cell).

##### Momentum 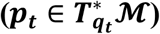

The generalized directional velocity scaled by mass. It lives in the cotangent bundle (*T*^∗^ℳ), which ensures that momentum calculations perfectly match the curvature of the shape space.

##### Mechanism / Regulatory State (*r*_*t*_)

The hidden variables driving changes from within. In robotics, this is the battery depletion or motor heat. In tissue morphogenesis, this is the chemical morphogen concentration or gene expression level.

#### 2.9.2. Anatomy of the Modified Hamiltonian Function

A classical Hamiltonian measures total energy. Here, the energy function *H*_*θ*_ is parameterized by learnable network weights *θ* and accepts external controls *u* and environments *e*. Formally, we define a Hamiltonian:

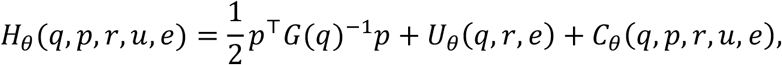

which has three terms:

##### Kinetic Energy 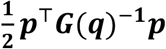

Expresses physical movement. *G*(*q*) is the Riemannian metric tensor, acting as a position-dependent mass matrix that shifts as the system alters its geometry.

##### Potential Energy *U*_*θ*_(*q, r, e*)

Represents stored energy fields. It links physical coordinates *q*, internal regulatory systems *r*, and environmental forces *e*.

##### Interaction/Control Energy *C*_*θ*_(*q, p, r, u, e*)

Accounts for intentional inputs *u* or non-conservative energy exchanges modifying the baseline landscape.

#### 2.9.3. Understanding the System Dynamics

The system evolves over time using a coupled set of differential equations.

The geometric dynamics are

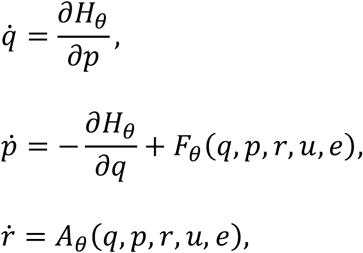

where:

##### Velocity Equation 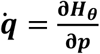

The change in position is strictly tied to generalized momentum. This guarantees smooth, physically realistic trajectories across the manifold.

##### Force Equation 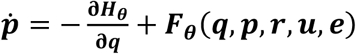

Momentum changes according to conservative potential slopes 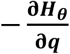 plus a non-conservative force vector ***F***_***θ***_ (***q, p, r, u, e***). ***F***_***θ***_ (***q, p, r, u, e***) models real-world effects like muscle friction, actuator drag, or biological dissipation.

##### Regulatory Equation 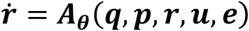

The internal states change according to their own transition network ***A***_***θ***_. Crucially, this rate depends on the physical states (*q, p*), capturing how movement or stress triggers chemical changes or power draw.

Thus the full manifold-mechanistic state evolves as

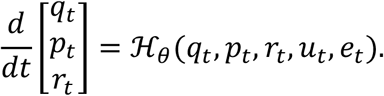

This is useful when reasoning must respect physical, biological, or energetic constraints.

#### 2.9.4. Why This Matters for Manifold Reasoning

Traditional neural networks struggle with physical reasoning because they minimize pixel errors or generic losses without understanding structural rules. This formulation provides distinct advantages:

##### No Unphysical Drift

By operating over the cotangent bundle *T*^∗^ℳ, the system cannot drift into mathematically impossible configurations (e.g., a robotic arm bending backward or a biological tissue tearing unnaturally).

##### Data Efficiency

Because the structural backbone of energy conservation is hard-coded into the derivative equations, the AI needs significantly less training data to predict complex trajectories accurately.

Coupled Feedback loops: It natively tracks how physical stress (*q, p*) alters internal biochemistry or actuator limits (*r*), and vice versa, which is vital for studying living tissues or complex robotics.

Figure 5 visualizes Hamiltonian formulation for manifold reasoning with an illustrative example (tissue recovery after treatment).

**Figure 5.**
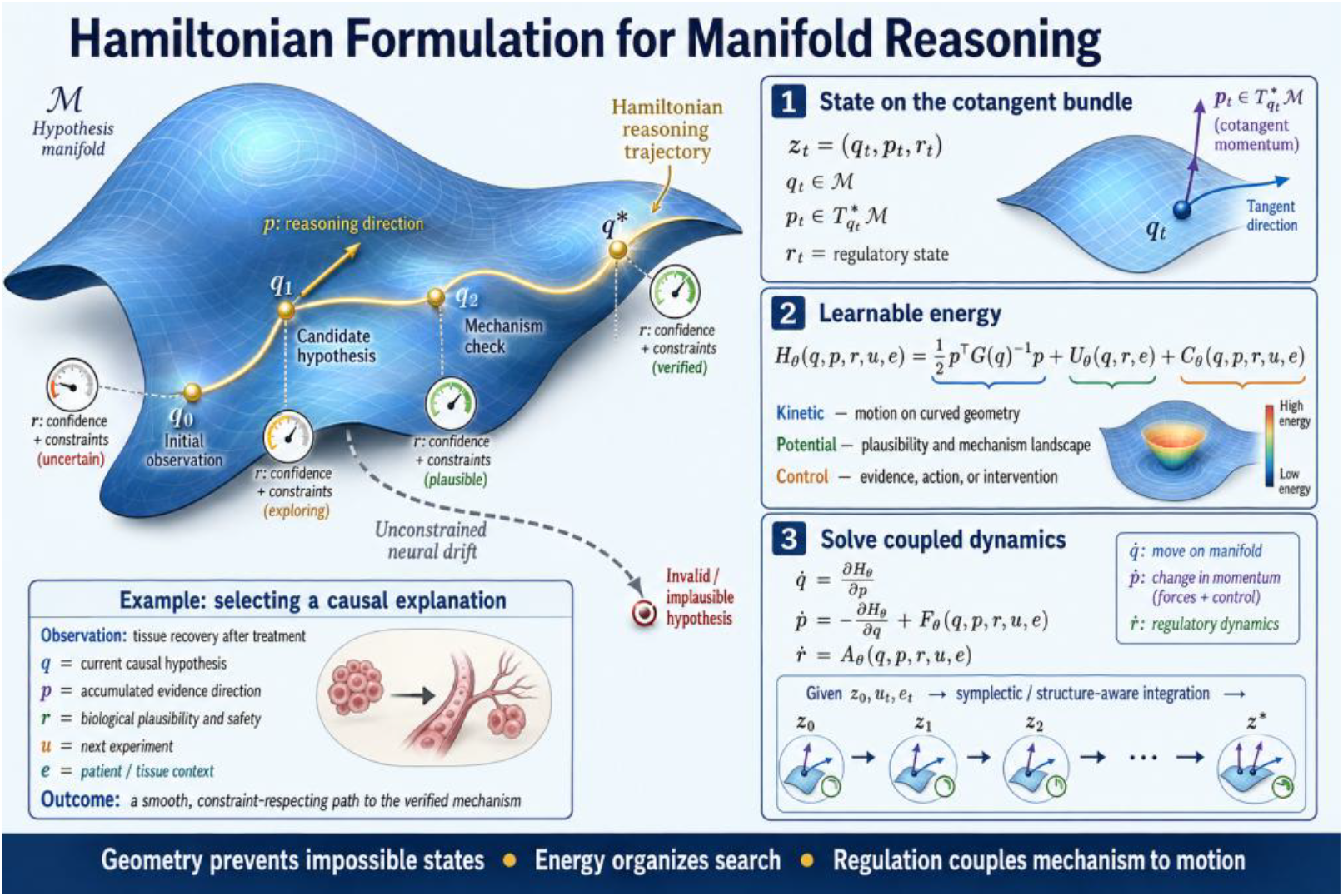
Hamiltonian formulation for manifold reasoning.

### 2.10. Geodesic planning

A geodesic plan is a sequence of states or actions designed to move an agent from a starting point to a destination along the shortest or most energy-efficient path on a curved surface (a manifold), rather than across flat, empty space. In everyday terms, a straight line is the shortest path on a flat map. However, if you fly from Tokyo to New York, the shortest path on the curved globe looks like an arc. That arc is a geodesic. A geodesic plan ensures that an AI or robot plans its movements using the natural geometry of its environment or its own joint constraints, preventing it from attempting physically impossible movements.

#### 2.10.1. Euclidean Planning vs. Manifold Planning

The text contrasts how planning changes when moving from flat spaces to curved manifolds.

##### Euclidean Planning (Flat Space)

In Euclidean space, planning often finds a short path:

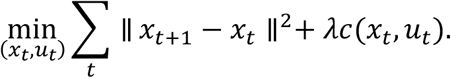

This standard approach minimizes the straight-line distance between consecutive steps ∑_*t*_ ∥ *x*_*t*+1_ − *x*_*t*_ ∥^2^ while balancing a cost function *c*(*x*_*t*_, *u*_*t*_) (like fuel or safety) scaled by a weight *λ*. It assumes the environment is flat and unconstrained.

##### Geodesic Planning (Curved Space)

On a manifold, the path should respect geodesic geometry:

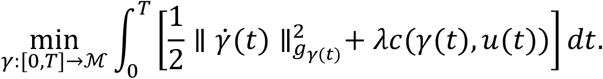

Subject to

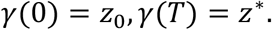

On a manifold ℳ, a path is a continuous curve *γ*(*t*). Velocity is measured using a Riemannian metric *g*_*γ*(*t*)_, written as 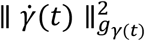. This metric changes depending on where you are on the surface, automatically accounting for terrain, constraints, or obstacles. The optimization guarantees the path natively bends to fit the environment’s true shape.

#### 2.10.2. The Uncontrolled Geodesic Equation

If an object is left to move freely without any steering wheels, engines, or external forces, it will naturally follow a geodesic. This is described by the geometric equation:

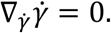

In simple terms, 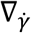 is the covariant derivative, which measures acceleration while factoring in the curvature of the space. Setting it to zero means the path has zero geometric acceleration—it is the straightest possible line on that curved surface.

To actually compute this on a computer, we convert it into local coordinates:

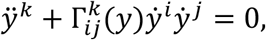

where 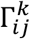 are Christoffel symbols,

*ÿ*^***k***^: The standard coordinate acceleration,

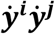: The current velocities in different directions interacting with one another,

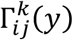: The Christoffel symbols. These act like a “geometric correction factor.” They calculate how much the coordinates themselves warp and bend at position *y*, twisting the path so it stays on the manifold.

#### 2.10.3. Controlled Reasoning (Steering the System)

Pure physics is rarely enough for an AI agent; the agent needs to make decisions to reach a target. For controlled reasoning, the formulation introduces an acting force. In other words, for controlled reasoning, the equation becomes

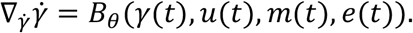

Instead of zero acceleration (= 0), the geometric acceleration is now driven by a controlled policy network *B*_*θ*_, parameterized by learnable weights *θ*. This force depends on:

***γ***(***t***): The current position on the manifold.

***u***(***t***): The control inputs (e.g., steering angle, motor voltage).

***m***(***t***): Internal memory, intent, or mental states of the agent.

***e***(***t***): External environmental factors (e.g., wind, terrain friction).

Therefore, the agent does not merely move in a straight Euclidean line. The agent does not blindly try to cut through walls or ignore physical boundaries by moving in a Euclidean straight line. Instead, it computes a trajectory where its natural momentum maps perfectly to the shape of the space, and it applies minimal control forces *B*_*θ*_ to gently steer that natural trajectory toward its destination.

Figure 6 displays geodesic planning.

**Figure 6.**
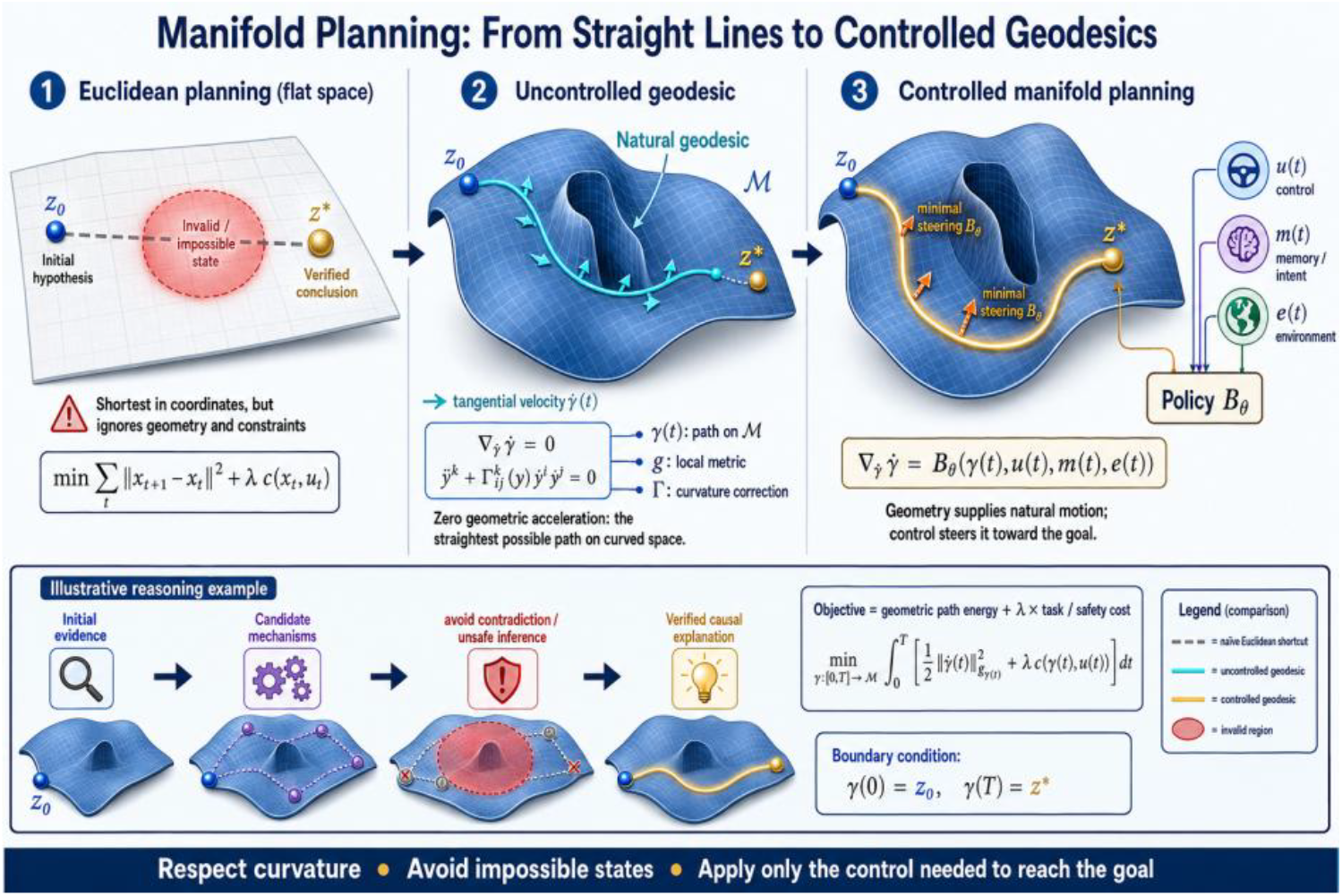
Geodesic planning.

### 2.11. HJB formulation on manifold space

This mathematical formulation describes the Hamilton-Jacobi-Bellman (HJB) framework extended to Riemannian manifolds. Instead of operating in a standard flat Euclidean space ℝ^*n*^, the system dynamics and optimization take place on a curved geometric space ℳ, such as a sphere, torus, or data manifold.

#### 2.11.1. The Cost Functional

For continuous-time optimal reasoning, define the cost functional:

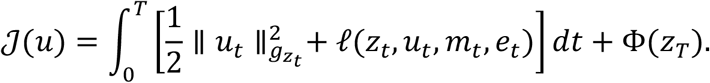

##### What it does

Measures the total “penalty” or cost of a chosen control strategy *u* from time 0 to *T*.

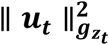: This is the control effort (kinetic energy). Because it is on a manifold, the norm is weighted by the Riemannian metric tensor *g* at the current state *z*_*t*_.

***ℓ***(***z***_***t***_, ***u***_***t***_, ***m***_***t***_, ***e***_***t***_): The running cost (e.g., state error, task-specific penalties). The variables *m*(*t*) and *e*_*t*_ represent auxiliary variables, context, or environmental factors.

**Φ**(***z***_***T***_): The terminal cost, punishing the system if it ends up in an undesirable final state *z*_*T*_ at time *T*.

#### 2.11.2. Controlled Manifold Dynamics

The controlled manifold dynamics are

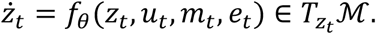

##### What it does

Describes how the state moves over time.

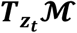**: (Tangent Space):** A manifold is curved, so you cannot just add a velocity vector directly to a state. Instead, the velocity *ż*_*t*_ must live in the tangent space at the current point *z*_*t*_ (think of a flat sheet of paper touching a globe at exactly one point). The system function *f*_*θ*_ maps the current state and control inputs to this flat tangent space.

#### 2.11.3. The Value Function

The value function is

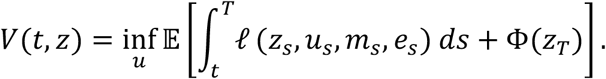

##### What it does

Represents the optimal remaining cost from time *t* up to the end horizon *T*, assuming you start at state *z* and act optimally (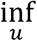 means taking the infimum, or absolute minimum).

(**Note:** The formulation provided omits the control effort term 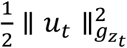 in the definition of *V*, but it is implicitly minimized as part of the total running cost package).

#### 2.11.4. The HJB Equation on the Manifold

The Hamilton-Jacobi-Bellman equation on the manifold is

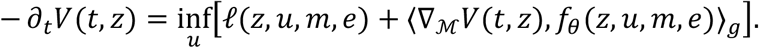

Here

∇_ℳ_*V* is the Riemannian gradient.

##### What it does

This is a partial differential equation (PDE) derived from the Principle of Optimality. It states that the time-decrease of your value function − ∂_*t*_*V*(*t, z*) must equal the minimized sum of your current running cost and your directional movement.

##### The Riemannian Inner Product <⋅,⋅>_*g*_

In flat space, you would use a standard dot product between the gradient of the value function and the system dynamics. On a manifold, you must use the metric inner product <⋅,⋅>_*g*_ to account for the curvature of the space.

##### ∇_ℳ_*V* (Riemannian Gradient)

This is the steepest ascent direction along the curved surface of the manifold.

#### 2.11.5. Local Coordinates Interpretation

In local coordinates,

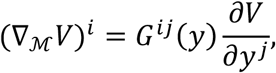

Here:

##### What it does

Explains how to actually compute the Riemannian gradient in practice using a local coordinate chart *y*.

***G***^***ij***^(***y***): This is the inverse of the Riemannian metric matrix *G* = *g*^−1^. It acts as a geometric correction factor. If the manifold stretches or compresses in certain directions, *G*^*ij*^ scales the standard partial derivatives 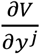 so that the gradient accurately reflects the true physical geometry.

#### 2.11.6. The Optimal Control

The optimal control is

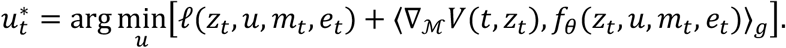

##### What it does

This provides the exact recipe for the best action to take at any split second.

**To find** 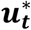, you look at your current state *z*_*t*_ and choose the control *u* that balances two things: minimizing your immediate cost (*ℓ*) and steering the system dynamics (*f*_*θ*_) downhill along the value function surface (∇ ℳ*V*).

### 2.12 Manifold memory

W#e introduce a framework that defines Manifold Memory as a geometry-aware memory system designed for artificial intelligence agents reasoning on non-Euclidean spaces ℳ. Instead of representing memory as a flat vector in Euclidean space (ℝ^*k*^), it structures memory as a dynamic set of “anchors” pinned directly to a geometric manifold, retrieving and updating information based on geodesic distances (the shortest paths along curved surfaces) rather than straight-line Euclidean distances. Here is the step-by-step mathematical formulation and breakdown of how this system operates.

#### 2.12.1. Memory Representation

Agent reasoning requires memory. In Euclidean models, memory may be a vector:

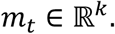

For manifold reasoning, memory should be geometry-aware. Let memory be a set of anchors on the manifold:

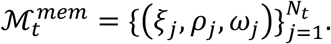

Each anchor *j* is a triplet containing three distinct mathematical components:

##### Geometric State *ξ*_*j*_ ∈ ℳ

A specific point (a stored manifold state) located on the manifold ℳ that serves as the spatial key or context for the memory.

##### Associated Knowledge (*ρ*_*j*_)

The actual content stored at that anchor (e.g., a rule, mechanism, semantic vector, or hypothesis).

##### Reliability (*ω*_*j*_)

A scalar weight indicating the trustworthiness, strength, or confidence level of this specific memory anchor.

#### 2.12.2. Geometry-Aware Memory Retrieval (Read)

When an agent encounters a current query or state *z*_*t*_ ∈ ℳ, it retrieves an aggregated memory vector *m*_*t*_ via a geodesic attention mechanism:

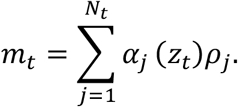

The attention weight *α*_*j*_(*z*_*t*_) determines how much the agent focuses on anchor *j*, formulated as a specialized softmax function:

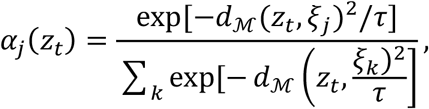

where:

***d***_**ℳ**_(***z***_***t***_, ***ξ***_***j***_): The geodesic distance function on the manifold ℳ measuring the true curvature-aware distance between the current state *z*_*t*_ and the memory anchor *ξ*_*j*_.

***τ***: A temperature hyperparameter controlling the sharpness of the retrieval distribution (lower values make retrieval exclusive to the closest anchors).

Thus memory retrieval is based on geodesic similarity, not Euclidean distance.

#### 2.12.3. Memory Overwrite & Update

When existing memories are relevant to the current situation, their internal knowledge payloads are updated using a localized gradient-style step:

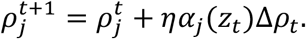

Where:

**Δ*ρ***_***t***_: The error correction, new observation, or gradient update generated at time *t*.

***η***: The learning rate.

***α***_***j***_(***z***_***t***_): Actively scales the update. Because *α*_*j*_(*z*_*t*_) decays exponentially with geodesic distance, only anchors geometrically close to the current state *z*_*t*_ are modified, preserving distant, unrelated memories.

#### 2.12.4. Grow-Don’t-Overwrite Allocation

Instead of forcing new information to overwrite old information when an agent encounters something completely foreign, the system grows dynamically.

New memory anchors are created when novelty is high:

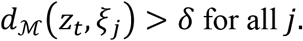

Then, a brand-new anchor is spawned at that exact coordinate:

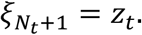

This gives a **grow-don’t-overwrite** memory mechanism. The total number of memory anchors increments (*N*_*t*+1_ = *N*_*t*_ + 1), allowing the network to permanently catalog new experiences without suffering from catastrophic forgetting.

### 2.13. Manifold hypothesis generation

To generate and formalize a manifold hypothesis, an agent shifts from flat Euclidean vectors to the curved geometric landscape of a smooth manifold ℳ. Instead of simply adding a noise vector, the agent uses the local geometry (tangent spaces and geodesic paths) to propose, move, and validate its beliefs. Here is the exact mathematical formulation broken down into a structured, four-step agentic process: Generate → Transport → Verify → Commit.

#### 2.13.1. Generate Local Perturbations

An agent should generate candidate explanations or hypotheses.

In Euclidean space, one may generate a set of *R* candidate hypotheses as simple vectors:

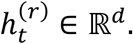

On a curved manifold ℳ, a hypothesis cannot just be a random point, or it risks falling off the space. Instead, it must be generated as a tangent vector inside the tangent space 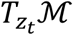 at the current state *z*_*t*_.

The agent uses a neural network generator 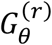 parameterized by *θ*. It takes the current state *z*_*t*_, internal memory *m*_*t*_, and environment context *e*_*t*_ to output a directional velocity vector:

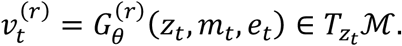

This yields a set of *R* candidate direction hypotheses:

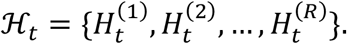

#### 2.13.2. Transport via Geodesics

To see where a directional hypothesis actually leads, the agent must project this straight tangent vector back down onto the curved surface of the manifold. This is done using the Exponential Map 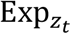, which generalizes the concept of “moving straight line in a flat space” to moving along a geodesic (the shortest path on a curved surface).

The predicted next state 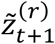 for each hypothesis *r* is formulated as:

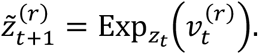

#### 2.13.3. Verify and Score Candidates

Each candidate hypothesis is evaluated using a scoring function *S*_*θ*_. This function takes into account the current state, the proposed tangent vector, the predicted state, memory, and environment cues:

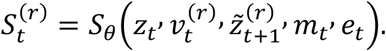

The comprehensive formulation breaks the total score down into four distinct structural constraints, prediction, geometry, verification/mechanism and causality:

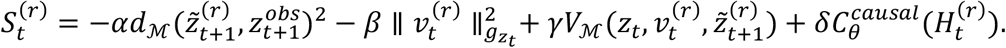

##### Prediction Error −*αd*_ℳ_

Measures the squared manifold distance *d*_ℳ_ between the prediction 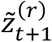 and the actual observation 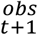. Higher error penalizes the score.

##### Geometric Least Action 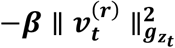

Penalizes excessive kinetic energy or long path lengths using the Riemannian metric tensor 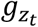. It enforces an economy of movement (Occam’s razor).

##### Verification/Mechanism *γV*_ℳ_

A positive reward mapping how well the transition aligns with known physical laws or structural mechanisms of the manifold.

##### Causal Rule Alignment 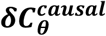

Evaluates whether the hypothesis 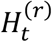 adheres to learned logical causal rules.

#### 2.13.4. Commit to the Optimal State

Finally, the agent uses an argmax selection policy to choose the best performing hypothesis index: *r*^∗^:

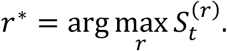

Once chosen, the agent commits to this hypothesis, updating its official trajectory belief state to the predicted destination of the winning candidate:

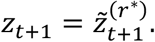

This converts agentic reasoning into **generate → transport → verify → commit** on a manifold. To help readers to understanding manifold hypothesis generation, we plot Fogure7 to visualize how the manifold hypothesis is generated.

**Figure 7.**
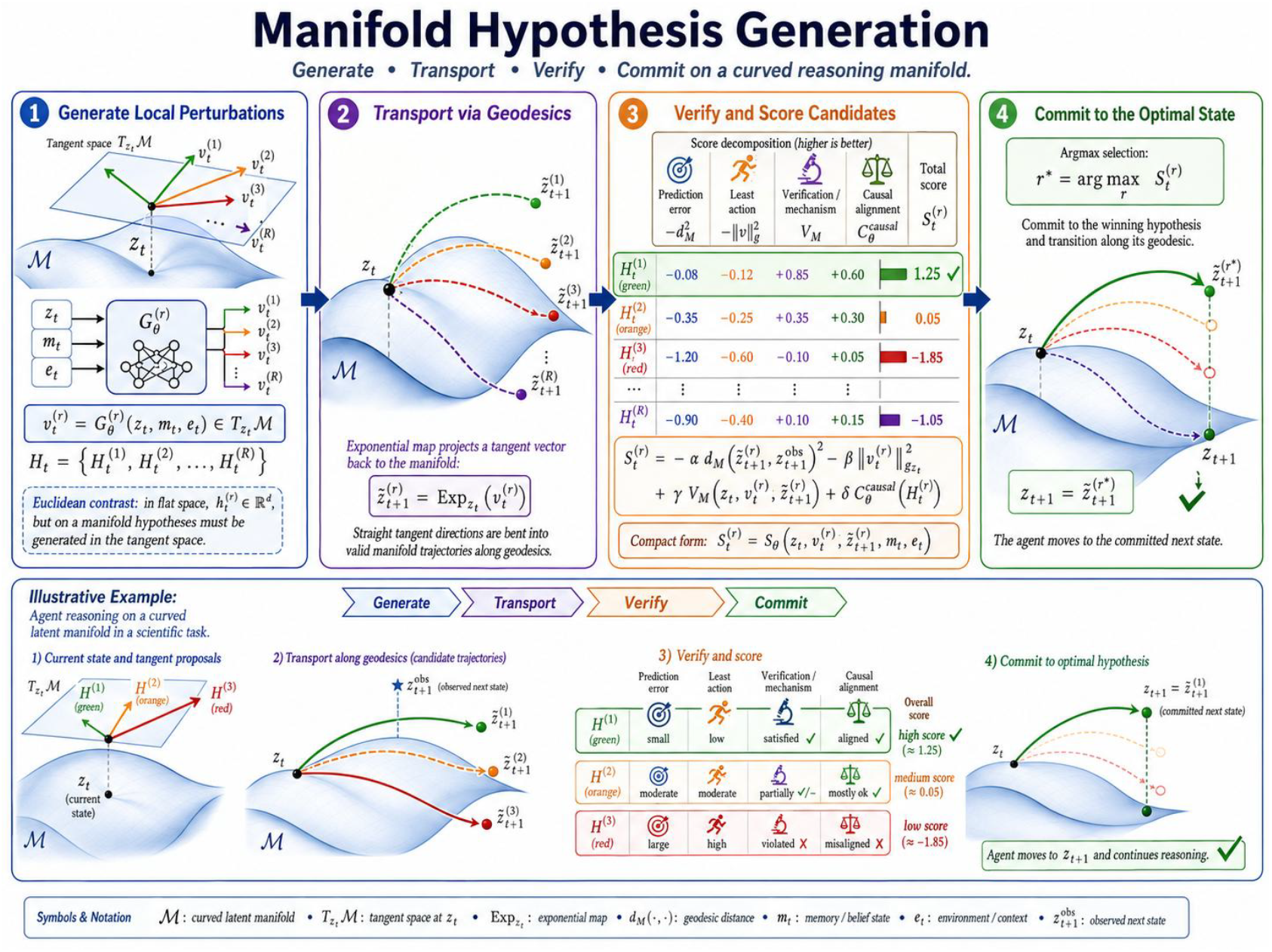
Manifold hypothesis generation.

### 2.14. Verification-gated manifold reasoning

A key problem in agent reasoning is invalid inference. On a manifold, an invalid inference may violate geometry, causality, biology, physics, or domain constraints. Verification-Gated Manifold Reasoning is a theoretical, advanced framework in AI alignment and geometric deep learning designed to prevent autonomous agents from making invalid logical leaps or “hallucinations” (Ruan et al. 2026).

Instead of treating an AI’s intermediate reasoning steps as pure text, this framework models reasoning as a continuous trajectory moving across a high-dimensional mathematical manifold ℳ. Valid reasoning traces conform to the physical, causal, or logical constraints embedded into the shape of that manifold. If a reasoning step takes a “wild shortcut” that violates reality (e.g., ignoring gravity or breaking a mathematical rule), the agent drifts off the manifold.

To prevent this drift, a verification gate checks every transition before letting the agent commit to its next thought or action. If the step fails the check, the agent is forced to backtrack, repair, or regenerate its hypothesis.

The core mathematical architecture of the framework can be broken down into three main phases: Mapping the Step Geometry, Learning to Verify, and Gating the Agent’s Commitment.

#### 2.14.1. Mapping the Step Geometry (The Verifier Domain)

Define a verifier:

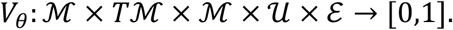

The system evaluates a single step transition from a current state *z*_*t*_ to a proposed future state *z*_*t*+1_ using five distinct geometric and environmental dimensions:

##### ℳ (The State Manifold)

The core underlying space where valid, real-world concepts exist.

##### *T*ℳ (The Tangent Bundle)

The space representing all possible valid directions and velocities (rates of logical change) the agent can move towards from its current spot.

##### *U* (Control/Action Inputs)

The specific tools, tokens, or intentional actions the agent chooses to apply.

##### *ℰ* (Environmental Context)

External constraints, such as physical laws, prompt parameters, or ground truth bounds.

When the agent proposes a transition *z*_*t*_ → *z*_*t*+1_, the verifier evaluates the precise trajectory using a logarithmic map:

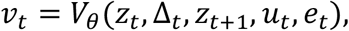

where

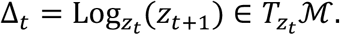

##### What Δ_*t*_ represents

In differential geometry, the logarithmic map 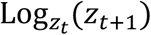 extracts the exact velocity vector in the tangent space 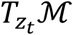 required to step from *z*_*t*_ to *z*_*t*+1_ along a shortest path (geodesic). This measures the “direction and distance” of the agent’s logic.

##### The Output *v*_*t*_

A continuous probability scalar between 0 and 1 representing how valid or “safe” the step is.

#### 2.14.2. Learning to Verify (The Loss Function)

To train the verifier function (*V*_*θ*_), the framework measures its error against a binary ground-truth target 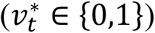—where 1 is valid and 0 is an invalid violation) using a Binary Cross-Entropy Loss function:

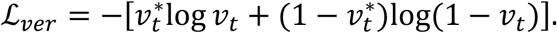

If a reasoning path is structurally valid 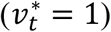, the loss penalizes low verifier confidence (*v*_*t*_ → 0).

If a reasoning path is invalid 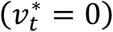, the loss heavily penalizes the verifier if it gets tricked (*v*_*t*_ → 1)

#### 2.14.3. Gating the Agent’s Commitment (The Admission Control)

Before the agent is allowed to execute the state transition, the step must pass through a strict Commitment Gate (*C*_*t*_):

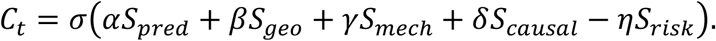

The gate utilizes a sigmoid function (*σ*) to squash a weighted combination of five distinct analytical sub-scores into a clean probability:

***S***_*pred*_: Likelihood score (how naturally this step follows according to the base AI model’s training).

***S***_***geo***_: Geometric alignment (how closely the step stays to the curvature of the valid manifold).

***S***_*mech*_: Mechanical/domain validity (conformity to explicit math formulas, code blocks, or physical properties).

***S***_*causal*_: Causal consistency (ensuring cause precedes effect, and that prerequisites are satisfied).

*S*_*risk*_: Threat/penalty multiplier (downweights the gate if the action carries catastrophic real-world risk).

#### 2.14.4. The Decision Rule (Admission Control)

The agent commits to a transition only if

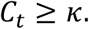

Otherwise, the agent repairs or regenerates hypotheses.

If ***C***_***t***_ ≥ ***κ***: The transition is admitted, and the agent permanently updates its state to *z*_*t*+1_.

If ***C***_***t***_ < ***κ***: The gate slams shut. The step is discarded, and the agent triggers an internal error-correction loop to repair or regenerate a completely new reasoning path.

To visualize the verification-gated manifold reasoning. we plot Figure 8 showinginformtion flow in the verification-gated manifold reasoning.

**Figure 8.**
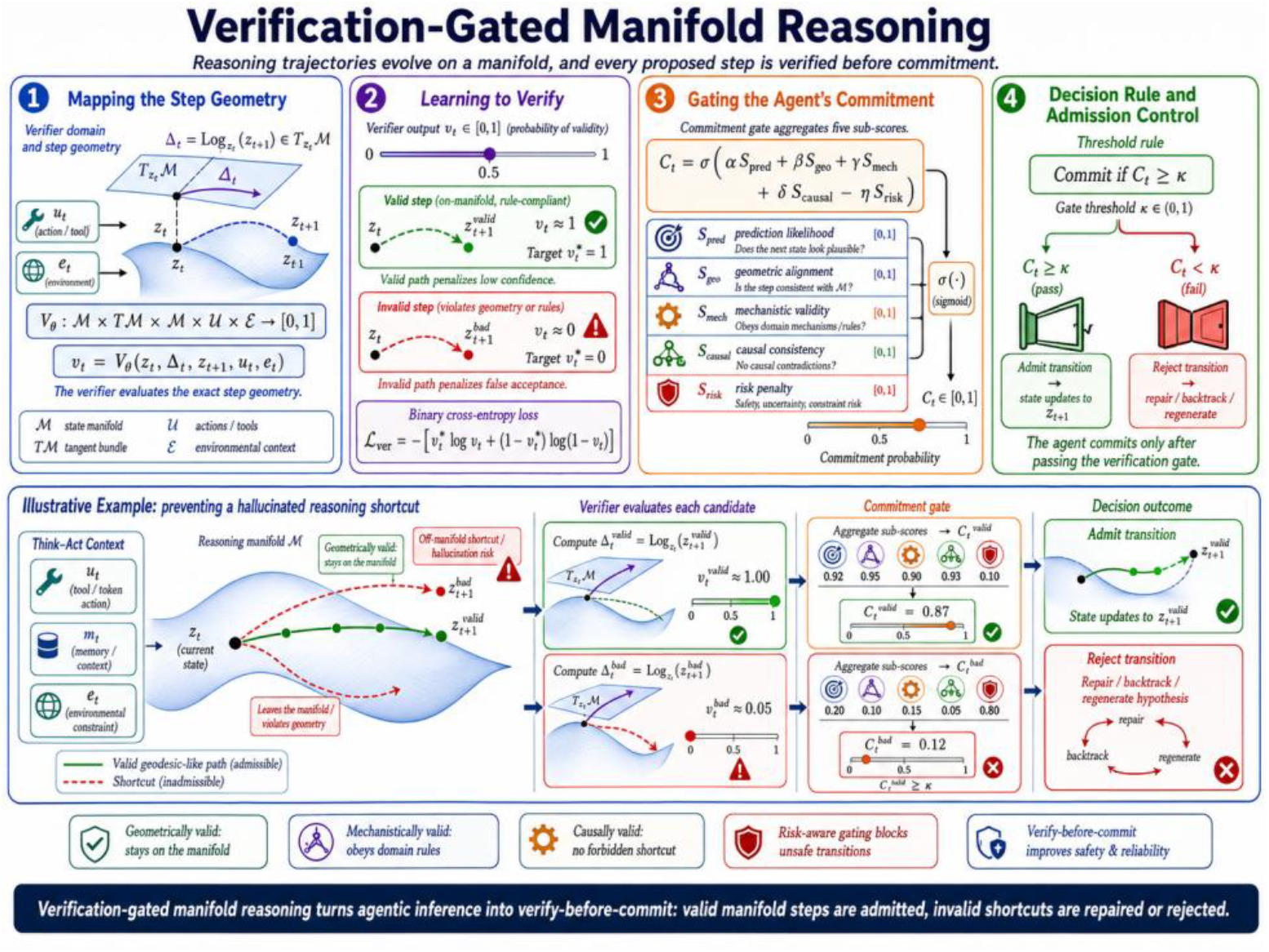
Verification-gated manifold reasoning.

### 2.15. Manifold self-repair

This formulation describes an optimization and geometric framework for manifold self-repair in artificial intelligence agents. It ensures that when an agent predicts or proposes an invalid next state (such as an physically impossible robot pose or a logically flawed plan), the agent should not simply accept the invalid transition. It should project that state back onto a valid “data manifold” ℳ before proceeding.

Here is the step-by-step breakdown of how manifold self-repair being formulated.

#### 2.15.1. Formulate the Optimization Problem

The first equation treats repair as a constrained optimization problem solved directly on the manifold ℳ. Suppose the proposed next state is 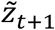. The repair module finds a nearby valid state:

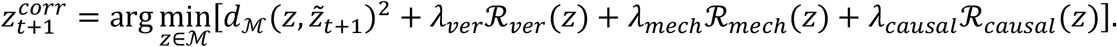

This objective function balances two competing goals: staying close to the agent’s original guess while enforcing physical and logical rules.

##### Proximity Constraint 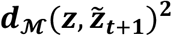

This computes the squared distance on the manifold between a valid state *z* and the corrupted proposal 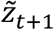. It ensures the corrected state remains as close to the original intent as possible.

##### Verification Regularizer (ℛ_*ver*_(*z*))

Penalizes states that violate basic environment verifications, scaled by the hyperparameter *λ*_*ver*_.

##### Mechanical/Physical Regularizer (ℛ_*mech*_(*z*))

Penalizes states that violate physical laws, kinematics, or conservation laws, scaled by *λ*_*mech*_.

##### Causal Regularizer (ℛ_*causal*_(*z*))

Penalizes states that break the logical or temporal causal chain of events, scaled by*λ*_*causal*_.

#### 2.15.2. Map to Tangent Space

Performing optimization directly on a curved manifold ℳ is computationally expensive. The second approach bypasses this by moving the correction step to a flat tangent space 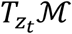 centered at the current state *z*_*t*_:

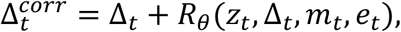

Instead of fixing the invalid state after it happens, this step fixes the velocity vector or update step (Δ_*t*_) that creates it.

**Δ**_***t***_: The original, uncorrected update step vector in the tangent space.

***R***_***θ***_(⋯): A learned neural network repair module parameterized by weights *θ*.

Context Inputs (*m*_*t*_, *e*_*t*_): The repair network takes the current state *z*_*t*_, the intended update Δ_*t*_, working memory *m*_*t*_, and environment feedback *e*_*t*_ to compute a vector correction.

#### 2.15.3. Project with Exponential Mapping

Once the corrected update vector 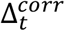 is calculated in the flat tangent space, it must be mapped back onto the curved surface of the manifold:

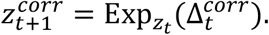

The Exponential Map 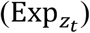 acts as a bridge. It takes the straight vector from the flat tangent space and rolls it cleanly onto the manifold’s surface along a geodesic (the shortest path on a curved surface). This guarantees that the final state 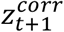 rests exactly within the bounds of valid reality (ℳ).

#### 2.15.4. Close the Regenerative Loop

The final expression summarizes the entire pipeline as a discrete runtime transformation:

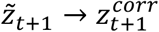

This represents a continuous, closed-loop regenerative correction mechanism. Rather than crashing, halting, or propagating errors forward when a plan fails verification, the agent intercepts the invalid 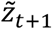 and dynamically warps it into 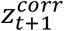.

The formulation establishes a dual-method system for self-repair: a global optimization method that pulls invalid states back to validity using penalization terms, and a local geometric method that corrects update trajectories within a tangent space before projecting them back to the manifold via an exponential map.

Figure 9 visualizes manifold self-repair process with concrete robotics as an illustrative example.

**Figure 9.**
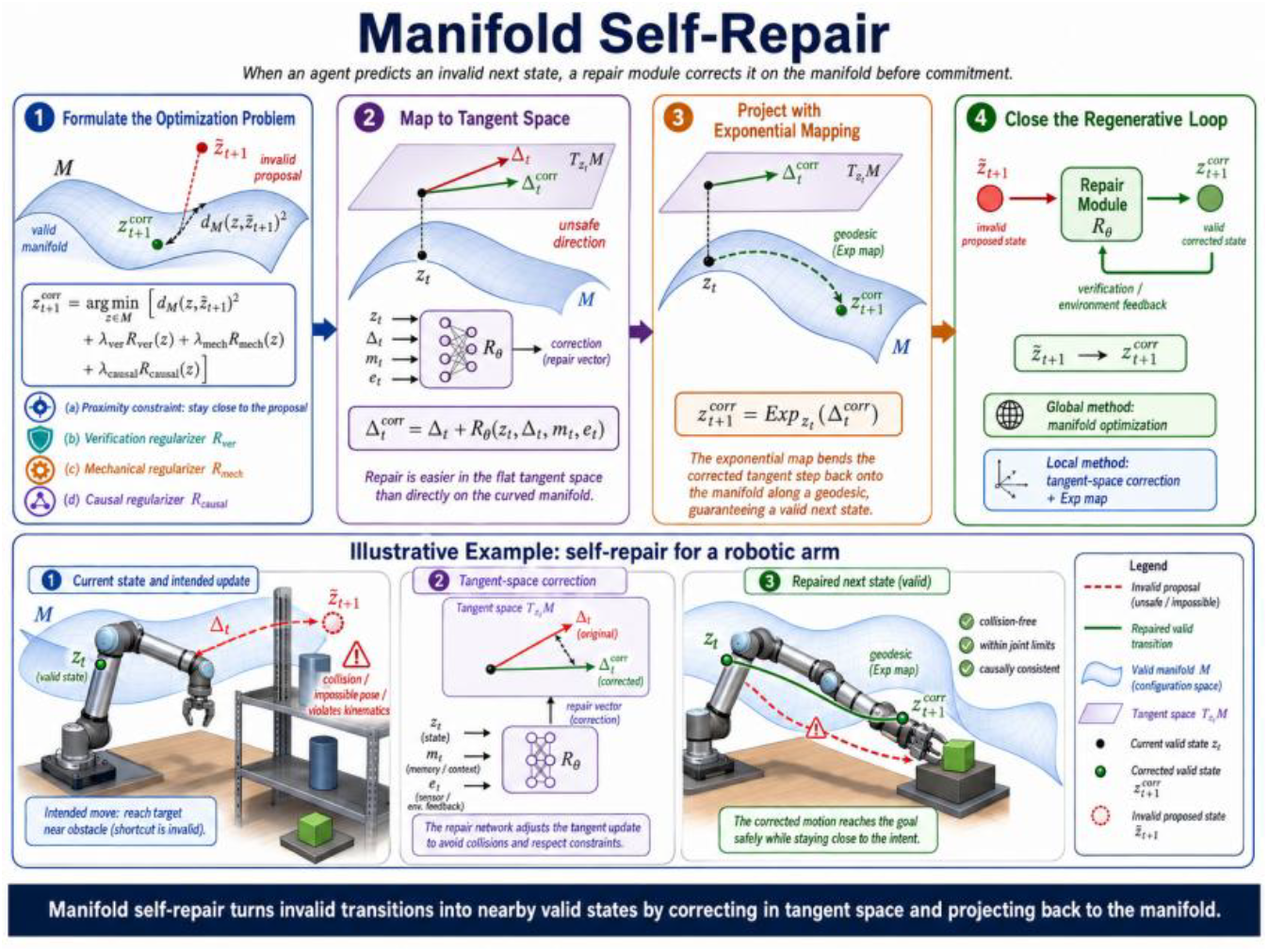
Manifold self-repair.

### 2.16. Graph-agentic reasoning on manifold space

The mathematical formulation of Graph-Agentic Reasoning on Manifold Space generalizes traditional Graph Neural Networks (GNNs) from flat Euclidean spaces to curved geometric structures (Riemannian manifolds). This framework allows autonomous agents to model complex biological and physical systems where data naturally lies on curved surfaces, such as hyperbolic spaces for hierarchies or hyperspheres for directional data.

Here is the proposed, step-by-step breakdown of the formulation.

#### 2.16.1. Define the Dynamic Graph Structure

For many biological and physical systems, the agent reasons over a graph:

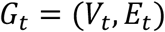

***V***_***t***_: The set of active nodes (e.g., biological entities, physical particles) at time *t*.

***E***_***t***_: The set of active edges representing relationships at time *t*.

#### 2.16.2. Formulate Manifold Node States

Unlike standard GNNs where node features belong to ℝ^*d*^, each node *i* possesses a state restricted to a Riemannian manifold ℳ:

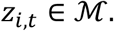

Because ℳ is curved, you cannot linearly add or subtract these state vectors directly without violating the geometric constraints of the space.

#### 2.16.3. Construct Edge Feature Vectors

Edges encode rich multi-modal relationships, acting as constraints or pathways for the reasoning agent. Each edge has features:

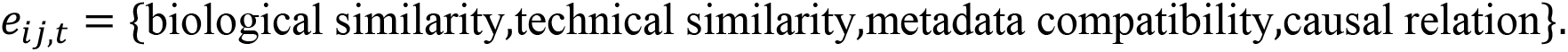

These features parameterize how information flows between nodes.

#### 2.16.4. Apply Tangent Space Transportation

To aggregate information from a neighbor*j* at node *i*, the neighbor’s state must be mapped into a local flat vector space where linear algebra is valid. This local space is the tangent 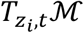. A manifold graph message from node *j* to node *i* should be transported into the tangent space of node *i*.

We compute the logarithmic map:

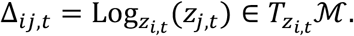

#### 2.16.5. Calculate Manifold Attention Weights

An attention mechanism determines the importance of neighbor *j* to node *i*. The scalar weight *a*_*ij*.*t*_ is computed using a scoring function *η*_*θ*_ passed through a Softmax normalization over the neighborhood *N*(*i*):

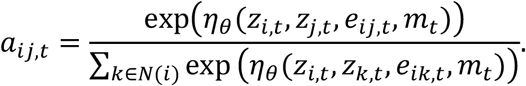

***η***_***θ***_: A neural network parameterized by *θ* that accounts for node states, edge properties, and global context *m*_*t*_.

#### 2.16.6. Aggregate Tangent Messages

Because all neighbor vectors Δ_*ij,t*_ have been mapped into the same tangent space 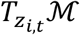, they can now be scaled and added linearly:

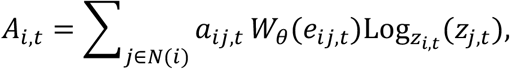

***W***_***θ***_(***e***_***ij***,***t***_): A learnable linear transformation tensor mapped to the edge features.

***A***_***i***,***t***_: The final aggregated message vector, safely residing in the tangent space 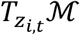.

#### 2.16.7. Execute Manifold Node Update

The agent updates the node’s state by translating the aggregated tangent message back onto the curved manifold surface. This uses the Exponential Map 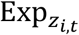, which projects a tangent vector back onto the manifold along a geodesic:

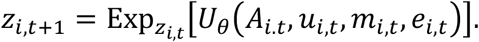

***U***_***θ***_: An update function (like a Riemannian GRU or MLP) that combines the message *A*_*i,t*_, local agent state *u*_*i,t*_, local memory *m*_*i,t*_, and self-edge features *e*_*i,t*_.

##### Euclidean Reduction (The Flat-Space Proof)

When the manifold ℳ is a flat Euclidean space (ℝ^*d*^), the tangent space at any point is identical to the space itself. The geometric operators simplify standard vector arithmetic:

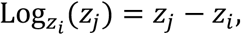

(The geodesic velocity vector is simply the displacement vector),

and

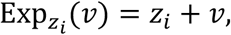

(Moving along a straight velocity line is simple vector addition).

This reduces to the following ordinary graph message passing.

Message Aggregation Reduction:

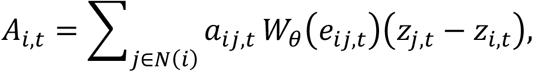

Node Update Reduction:

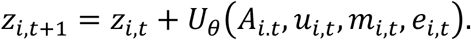

This mathematical reduction proves that standard Graph Neural Networks are merely a special, restricted case of this generalized manifold framework.

Figure 10 displays the architecture of graph-agentic reasoning on manifold space with tissue regeneration illustrative example.

**Figure 10.**
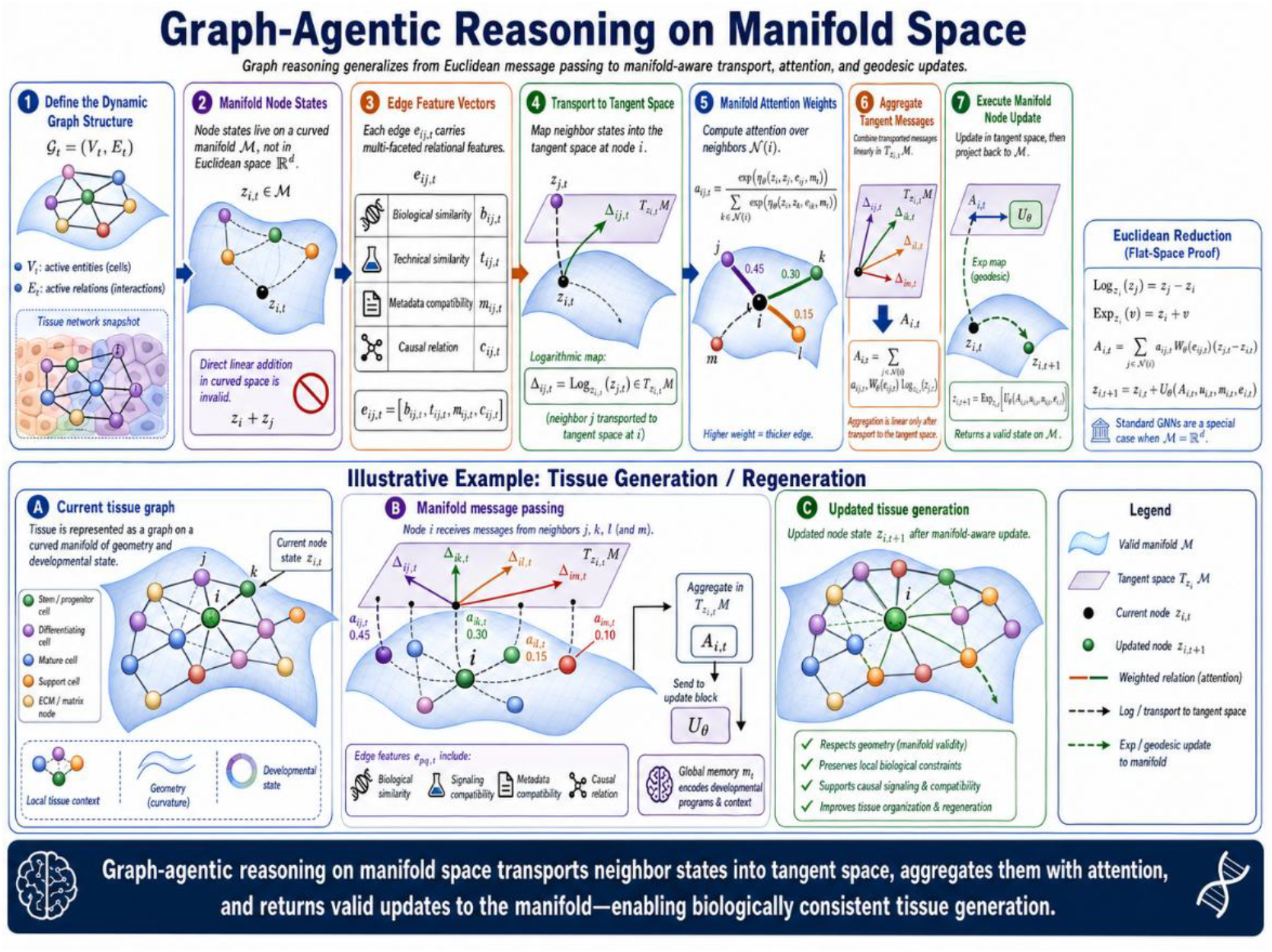
Graph-agentic reasoning on manifold space.

### 2.17. Causal manifold reasoning

Many scientific agents must reason causally, not only predictively. The core mathematical framework of causal manifold reasoning blends structural causal models (SCMs) with Riemannian geometry to model causal dynamics on non-Euclidean spaces.

Here is the step-by-step breakdown of its mathematical formulation.

#### 2.17.1. Define Causal Topology

The system structure is governed by a time-varying directed acyclic graph (DAG) denoted as

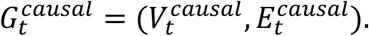

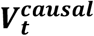: represents the set of vertices (variables) at time *t*.

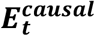: represents the set of directed edges defining causal dependencies.

**Every endogenous variable *z***_***j***_ is restricted to a smooth, non-Euclidean space called a Riemannian manifold, denoted as *Z*_*j*_ ∈ ℳ_*j*_.

#### 2.17.2. Formulate Structural dynamics

In a classic structural causal model, variables are updated via standard functions. Here, state transitions are mapped directly onto the manifold space using the exponential map, Exp_*p*_(*v*), which projects a tangent vector *n* from the tangent space *T*_*p*_ℳ back onto the manifold itself. Each variable corresponds to a manifold coordinate, mechanism, or observable:

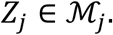

The state of variable *j* at time *t* + 1 is defined by:

Using exponential-map dynamics, the structural causal model for the state variable *j* at time *t* + 1 is defined by

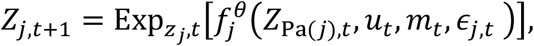

where

***Z***_***j***,***t***+**1**_ ∈ **ℳ**_***j***_: is the current point on the manifold acting as the base for the exponential map.

***Z***_**Pa**(***j***),***t***_ represents the states of the causal parent nodes of j at time t.

***u***_***t***_ is the action or control input.

***m***_***t***_ represents the context or environment state.

***ϵ***_***j***,***t***_ is the exogenous noise variable.

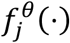 is a parameterized function (such as a neural network) that outputs a tangent vector in the tangent space 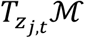.

#### 2.17.3. Factorize Interventional Transitions

An intervention represents an external modification of the system, written using Pearl’s do-calculus as do (*u*_*t*_ = *a*). This operation overrides the natural causal mechanisms of the intervened variables. Because the graph remains a DAG, the joint interventional transition probability factorizes according to the Markov property over the manifold states:

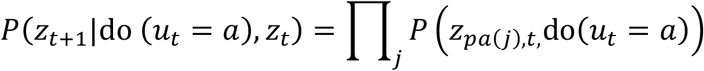

#### 2.17.4. Evaluate Counterfactual Queries

A counterfactual query determines what the state would have been under a different action *a*′, given that the actual observed state at time *t* was *z*_*t*_). It evaluates the structural system deterministically by fixing the shared background context *m*_*t*_ and the structural error terms *e*_*t*_:

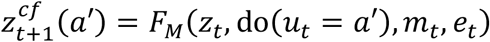

#### 2.17.5. Quantify Causal Effects

In Euclidean causal inference, the causal effect is usually measured by a simple subtraction of expected values (e.g.*E*[*Y*|do(*a*′)] − *E*[*Y*|do(*a*)]). Because straight subtraction is invalid on curved manifolds, the Causal Effect (CE) is measured using the geodesic distance *d*_ℳ_(⋅,⋅), which calculates the length of the shortest path between two points along the curved manifold surface:

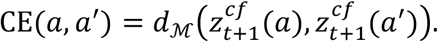

The framework successfully generalizes Pearl’s Causal Diagram framework into Riemannian geometry by replacing vector addition with the manifold exponential map and replacing Euclidean differences with geodesic distance.

### 2.18. Manifold hypergraph reasoning

Pairwise graphs are sometimes insufficient. Biological systems often depend on higher-order interactions:

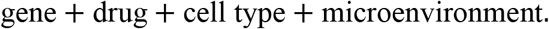

Below we introduce mathematical formulation of manifold hypergraph reasoning. The formulation describes a manifold hypergraph neural network designed for geometric *d* eep learning on non-Euclidean Riemannian spaces, capturing multi-entity biological interactions.

#### 2.18.1. Underlying Manifold Space

Use a causal hypergraph: ℋ_*t*_ = (*V*_*t*_, ℰ_*t*_),where each hyperedge *ε* ∈ ℰ_*t*_ connects multiple variables: *ε* = {*v*_1_, *v*_2_, …, *v*_*k*_}. The variables *z*_*v,t*_ and *z*_*ϵ,t*_ do not exist in flat vector space. They are points on a smooth curved surface called a Riemannian manifold ℳ (such as a Poincare hyperbolic disc used to model hierarchical biological trees). Traditional addition and subtraction do not work here; instead, the model uses geometry-aware operations.

#### 2.18.2. Hyperedge Message Construction

The hyperedge message formulation computes higher-order context:

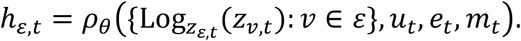

##### 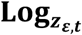 (*z*_*v,t*_) (Logarithmic Map)

This operator projects a point *z*_*v,t*_ from the curved manifold surface onto a flat tangent plane centered at the hyperedge reference point *z*_*ε,t*_. This converts curved geometric coordinates into flat Euclidean vectors so neural networks can process them safely.

{⋯}: The collection of flat vector representations for all nodes (e.g., “gene”, “drug”, “cell type”, “microenvironment”) belonging to that specific hyperedge *ϵ*.

***u***_***t***_, ***e***_***t***_, ***m***_***t***_: Global context vectors representing patient clinical status, environmental factors, or time-step variables.

***ρ***_***θ***_: A parameterised neural network layer that aggregates these tangent vectors into a unified multi-entity message vector *h*_*ε,t*_.

#### 2.18.3. Hyperedge Weighting Mechanism

The hyperedge weight scales message importance based on biological mechanism strength:

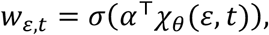

where

(**χ**_***θ***_(***ε, t***)): An encoding function evaluating the combined interaction state of all elements within hyperedge *ε* at time *t*,

***α***^⊤^: A learnable attention weight vector.

***σ*:** A sigmoid activation function squeezing the final attention score into a scalar probability weight between 0 and 1, representing interaction intensity.

#### 2.18.4. Node State Update

The state of individual nodes is advanced using a geometry-preserving manifold step:

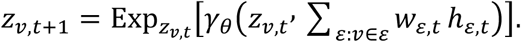

**∑**_***ε***:***v***∈***ε***_ ***w***_***ε***,***t***_ ***h***_***ε***,***t***_: A weighted sum combining messages from all hyperedges containing node *v*. This calculation happens safely inside the flat tangent space of node *v*.

***γ***_***θ***_: A neural network that updates the node’s tangent representation.

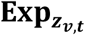 **(Exponential Map):** The inverse of the logarithmic map. It maps the updated flat tangent vector back down onto the curved Riemannian manifold surface, yielding the valid non-Euclidean node state *z*_*v,t*+1_.

### 2.19. Learning objective

This learning objective, denoted as ℒ_*MAR*_, represents a comprehensive, multi-objective loss function designed to train an advanced machine learning model. It goes beyond simple prediction accuracy by explicitly forcing the model to respect physics, geometry, causality, memory, and safety.

The full learning objective combines task accuracy, manifold geometry, dynamics, causal validity, memory retention, verification, and repair.

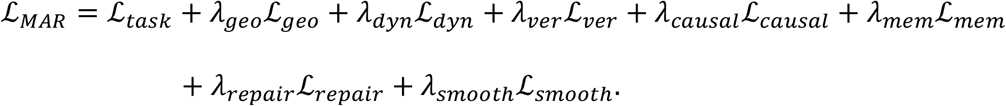

The task loss may be

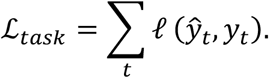

The dynamics loss is

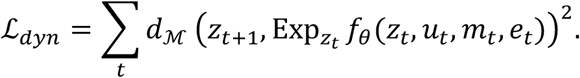

The causal loss is

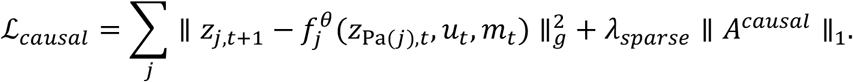

More geometrically:

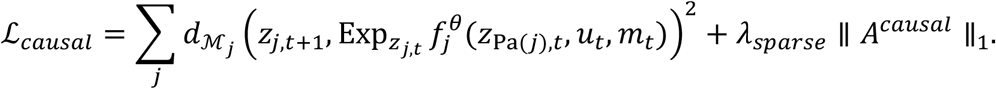

The memory retention loss is

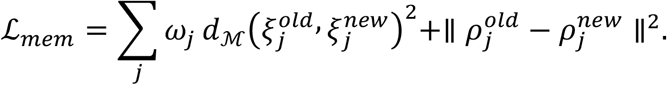

The repair loss is

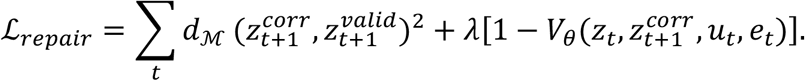

The smoothness loss is

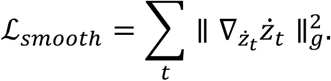

This penalizes unnatural acceleration on the manifold.

### 2.20. Full manifold-agent algorithm

A complete reasoning cycle is:

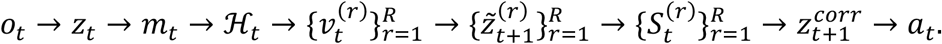

More explicitly:

**Step 1: Encode observation**

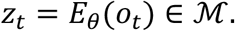

**Step 2: Retrieve manifold memory**

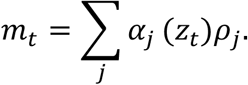

**Step 3: Generate candidate tangent hypotheses**

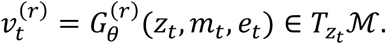

**Step 4: Project hypotheses onto manifold**

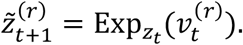

**Step 5: Score candidates**

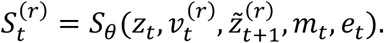

**Step 6: Verify**

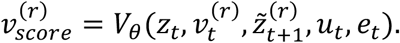

**Step 7: Commit or repair**

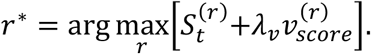

If

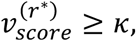

then

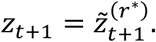

Otherwise,

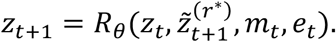

**Step 8: Choose action**

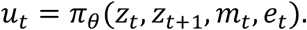

**Step 9: Update memory**

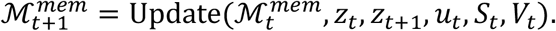

Full manifold-agent algorithm is complex. It is organized as a network with each node that may have possible multiple inputs and outputs. To help readers to understand architecture of the algorithm and how information flows in the algorithm, we plot Figure 11 to visualize full manifold-agent algorithm.

**Figure 11.**
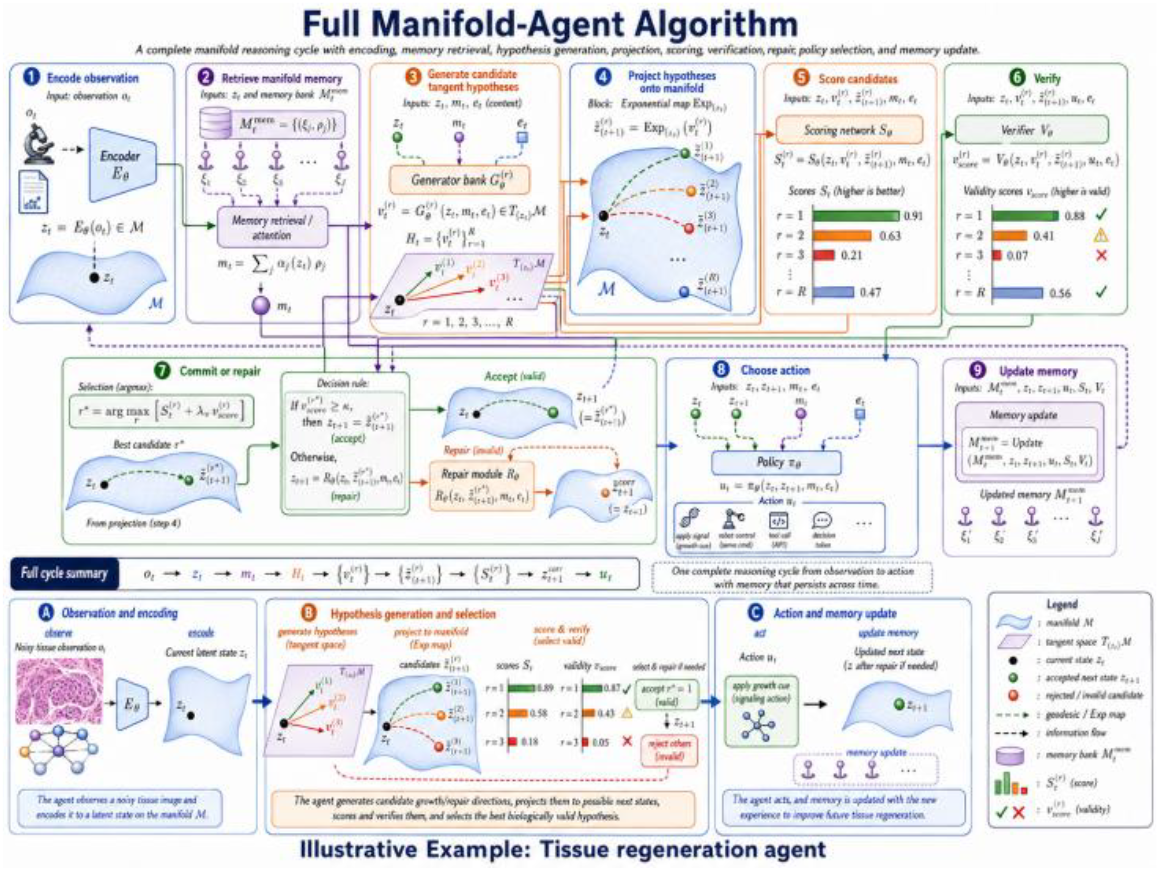
Full manifold-agent algorithm.

### 2.21. Relation to biological data analysis

For biological data, the manifold state may be

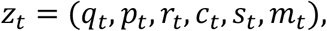

where *q*_*t*_ is tissue or cell-state configuration, *p*_*t*_ is developmental or dynamical momentum, *r*_*t*_ is regulatory mechanism state, *c*_*t*_ is cell-type composition, *s*_*t*_ is signaling or spatial state, *m*_*t*_ is nested biological memory.

The model may predict:

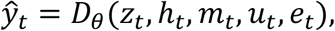

where *h*_*t*_ is a graph-connectome hidden state.

The biological reasoning transition becomes:

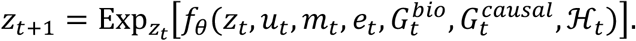

This allows the agent to reason over:

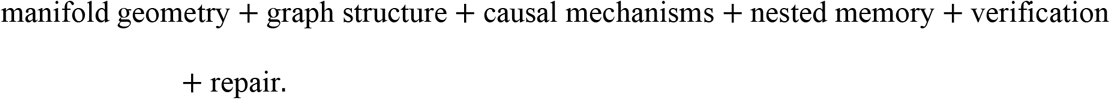

### 2.22. Relation to robotics and physical reasoning

For robotics, the manifold may include pose and configuration spaces:

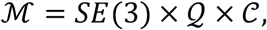

where *SE*(3) is rigid-body pose, *Q* is joint configuration space, *C* is contact mode space. The state is

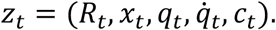

The transition is

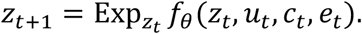

Contact-rich reasoning requires mode switching:

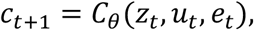

and different manifold dynamics for each contact mode:

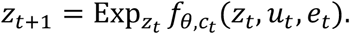

Thus manifold reasoning naturally handles non-Euclidean robot pose, geometric constraints, and contact transitions.

### 2.23. Euclidean reasoning as a special case

If

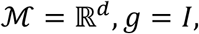

then

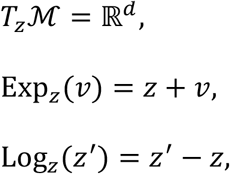

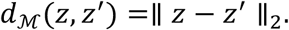

Therefore the manifold transition

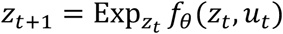

Becomes

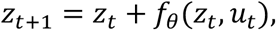

which is exactly the ordinary Euclidean residual update.

So the proposed manifold framework contains Euclidean agent reasoning as a special case.

### 2.24. Compact final formulation

The full manifold-agent reasoning model can be summarized as:

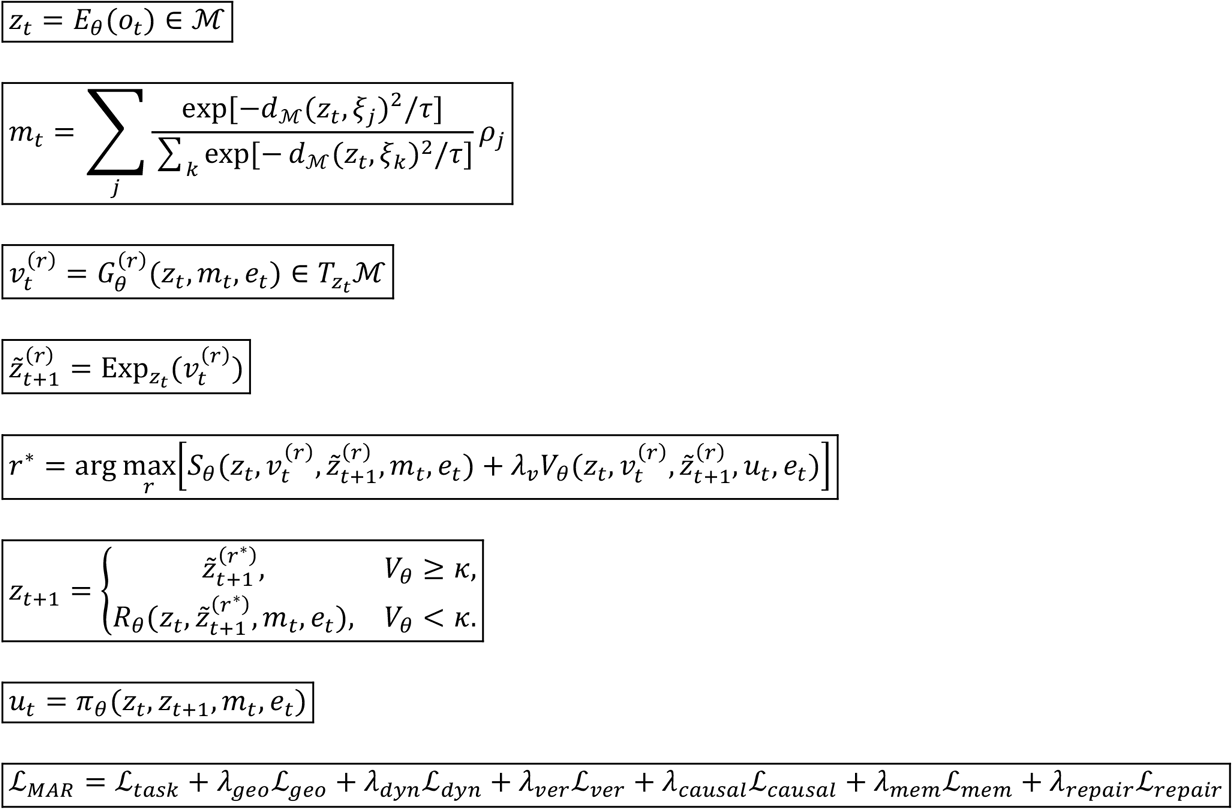

This is the mathematical core of extending agent reasoning from Euclidean space to manifold space.

### 2.25. Conceptual summary

The key replacement rules are:

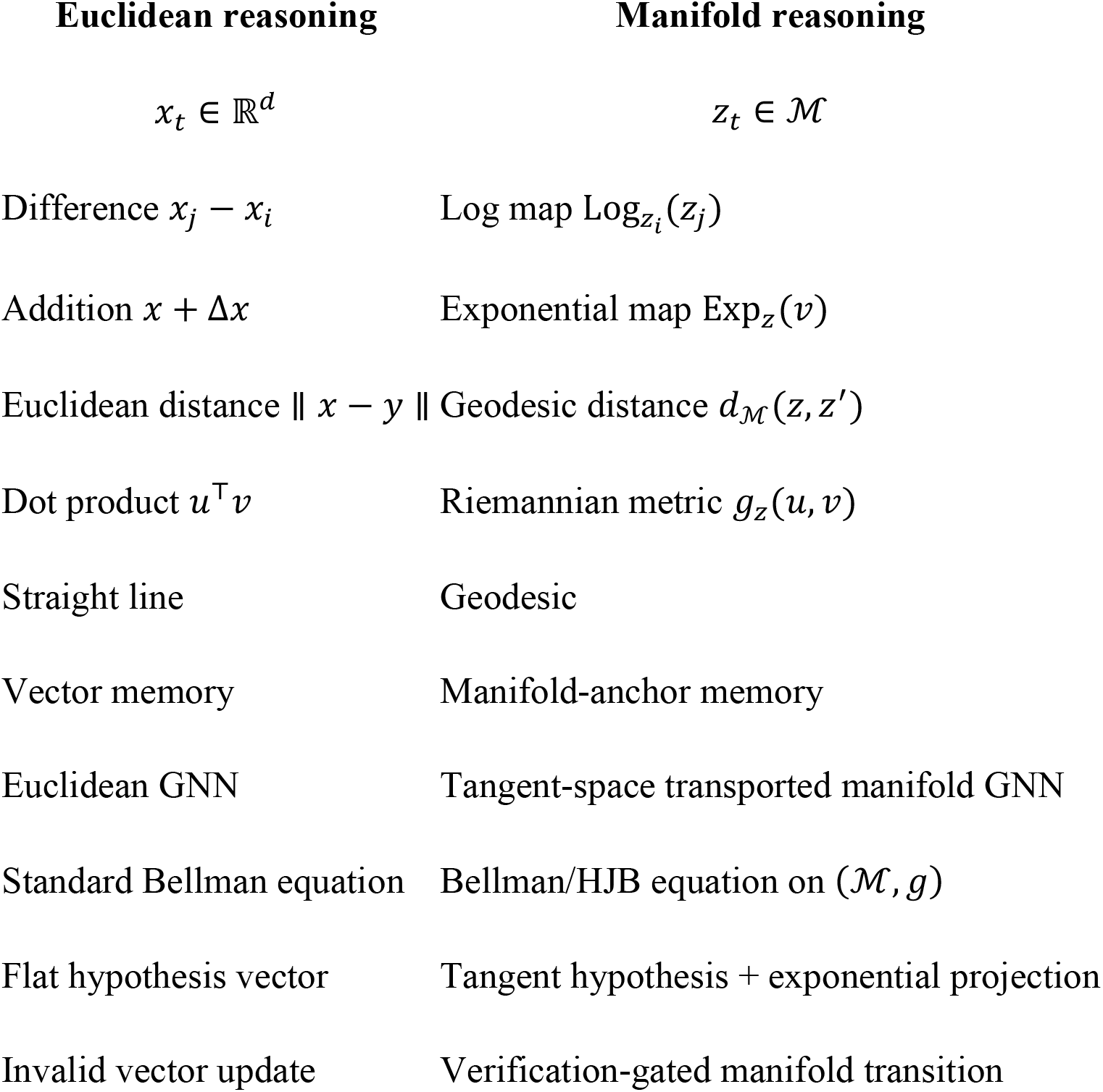

The essential shift is:

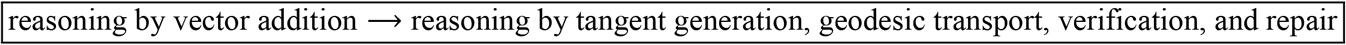

This gives a mathematically principled foundation for extending agentic reasoning to biological systems, robotics, manifold data analysis, and complex scientific discovery.

## 3. Extending Agentic POMDPs, Internal Reasoning Variables, and Post-Training Reinforcement Learning to Riemannian State Spaces

Extending Agentic POMDPs, Internal Reasoning Variables, and Post-Training Reinforcement Learning (RL) to Riemannian State Spaces is a highly viable, cutting-edge approach to manifold agentic reasoning. This mathematical framework allows autonomous agents to reason, plan, and learn on complex, non-Euclidean geometries (like spheres, tori, or positive definite matrices) which naturally encode physical constraints, rotational dynamics, and hierarchical data structures.

### 3.1. Geometric Agentic POMDPs

A standard Partially Observable Markov Decision Process (POMDP) is defined by the tuple:

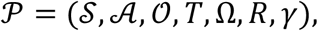

Where *s*_*t*_ ∈ *S* is the hidden environment state, *a*_*t*_ ∈ *A* is the external action, *o*_*t*_ ∈ *O* is the observation, *T*(*s*_*t*+1_ ∣ *s*_*t*_, *a*_*t*_) is the transition model, Ω(*o*_*t*_ ∣ *s*_*t*_) is the observation model, *R*(*s*_*t*_, *a*_*t*_) is the reward, and *γ* ∈ (0,1) is the discount factor.

#### Manifold-Valued Belief States

The agent does not directly observe *s*_*t*_. Instead, it maintains a belief state *b*, which becomes a probability distribution specialized for manifolds (such as a Riemannian Gaussian or a Von Mises-Fisher distribution):

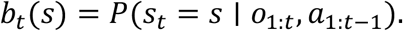

#### Riemannian Internal Reasoning Variables

Internal reasoning variables represent the agent’s hidden “mental” workspace (e.g., latent chain-of-thought, working memory, or implicit planning steps) before committing to an external action. The paper’s useful idea is to introduce an internal reasoning variable, say

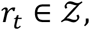

where *Z* is the thought or reasoning space.

Below mathematical formulation describes a Think-Then-Act cognitive loop for autonomous AI agents. Instead of reacting reflexively to a situation, the agent is forced to generate an internal “chain of thought” or planning step before committing to an external action.

#### The Contrast: Reflexive vs. Reasoned Behavior

##### Old Approach: Reflexive Policy

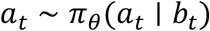

In standard reinforcement learning or basic LLM prompting, the agent looks at its belief state *b*_*t*_ and immediately outputs an external action *a*_*t*_. There is no dedicated time or space for the agent to weigh options, predict outcomes, or strategize.

#### New Approach: Think–Act Decomposition

Instead, it has a **think–act decomposition**. The policy is broken into two distinct sequential phases:

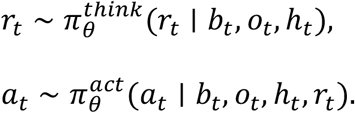

Here *h*_*t*_ is the interaction history or memory.

##### *z* ∈ *Z* (The Thought/Reasoning Space)

This is the agent’s internal workspace. If the agent is a Large Language Model (LLM), *Z* might be a sequence of hidden “thought tokens” (like OpenAI’s o1 or o3 models). If the agent is a control system, *Z* could be a trajectory plan or a simulated tree-search.

*r*_*t*_ (Internal Reasoning Variable): The specific strategy, deduction, or plan generated at time step *t*. Crucially, *r*_*t*_ is hidden from the environment; it only exists inside the agent’s mind.

***b***_***t***_ **(Belief State):** The agent’s current statistical understanding of where it is in the world, filtering out noise from past observations.

***o***_***t***_**(Current Observation):** The immediate sensory input or data packet received from the environment at the current moment.

***h***_***t***_**(Interaction History / Memory):** The long-term log of what happened in previous steps. This prevents the agent from repeating past mistakes.

So the agent first generates an internal reasoning state *r*_*t*_, and then chooses an external action *a*_*t*_.

#### Step-by-Step Execution Flow

This gives the core structure:

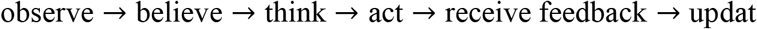

##### Information Gathering

The agent receives the latest observation *o*_*t*_ and combines it with its memory *h*_*t*_ and belief *b*_*t*_.

##### Step 1 **Think** 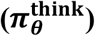

The agent processes this data to sample a reasoning state *r*_*t*_ from its thought space ***Z*** (Dai et al. 2023). Example: “If I move left, I might hit that wall. I should calculate a path to move around it.”

##### Step 2 **Act** 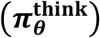

The agent looks at the exact same data, plus the reasoning *r*_*t*_ it just generated. It uses that reasoning to pick the final external action *a*_*t*_. Example: Executes the safe detour command.

To help readers to intuitively understand the geometric agentic POMDPs, we plot Figure 12 show the dynamic process of agentic POMDPs.

**Figure 12.**
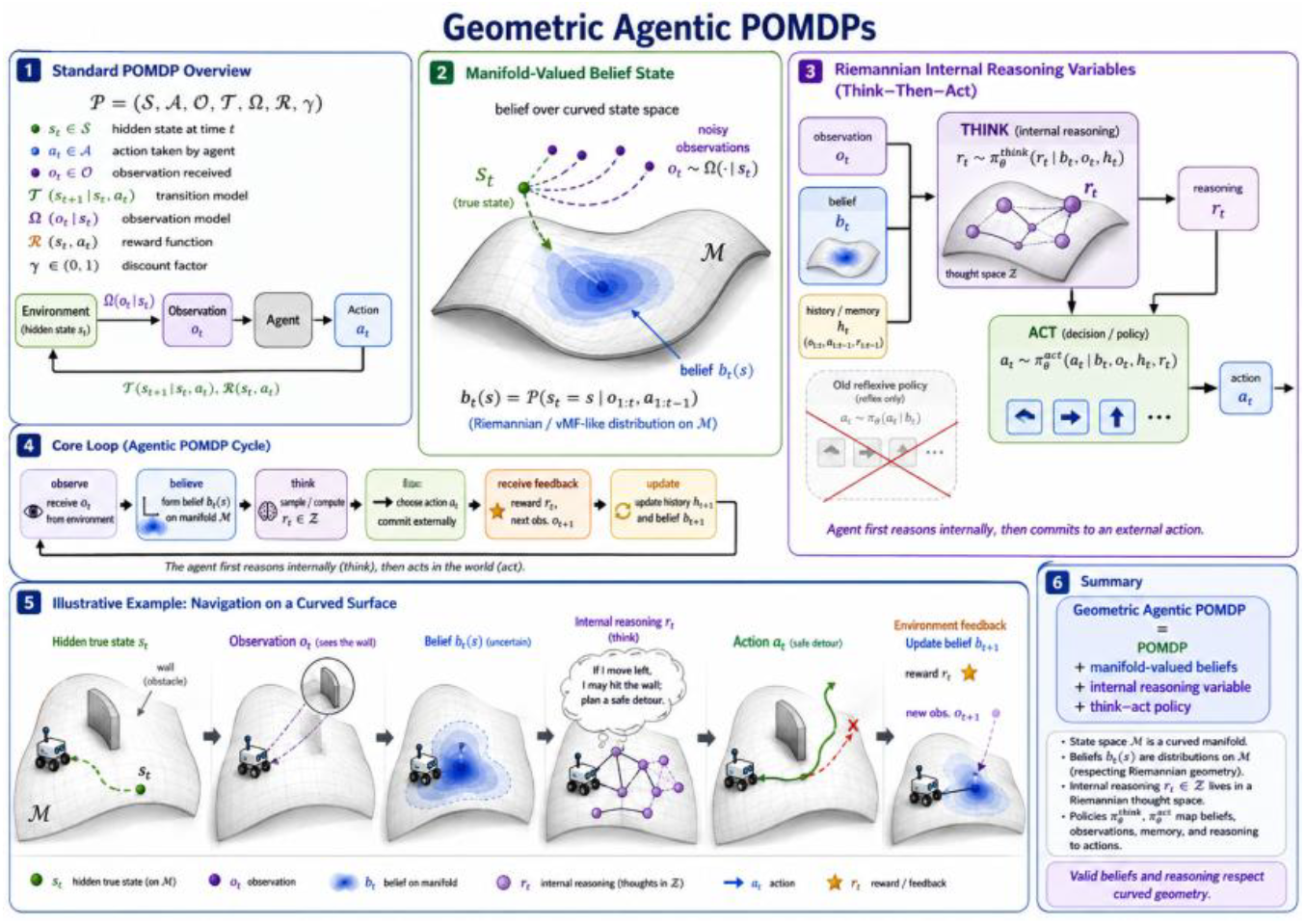
Geometric agentic POMDPs.

### 3.2. Extension to manifold space

This extension updates standard artificial intelligence frameworks by replacing flat, linear vector spaces with curved, structured surfaces called manifolds. Instead of assuming an AI agent reasons and moves in an infinite, unconstrained flat grid (ℝ^*d*^), this framework restricts the agent to non-Euclidean geometries (like spheres, tori, or complex continuous constraint surfaces) to better model real-world physics, rotation, and complex cognitive structures.

#### 3.2.1. From Flat Space to Manifolds

In standard reinforcement learning or robotic control, variables exist in flat Euclidean space:

##### Classic State Space (*s*_*t*_ ∈ ℝ^*d*^)

The environment state is a simple list of *d* numbers.

##### Classic Internal Reasoning (*r*_*t*_ ∈ ℝ^*k*^)

The agent’s internal thoughts or memory vectors are a simple list of *k* numbers.

In Euclidean agentic reasoning, one usually assumes

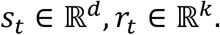

The manifold extension generalizes these to:

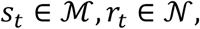

where (ℳ, *g*_ℳ_) is the environment-state manifold, and(*N, g*_*N*_) is the internal reasoning manifold. In other words, we have

##### Environment Manifold (*s*_*t*_ ∈ ℳ)

The environment state lives on a smooth manifold ℳ equipped with a Riemannian metric *g*_ℳ_. The metric *g*_ℳ_ calculates the true “shortest path” (geodesics) and distances on this curved space (e.g., navigating the curved surface of the Earth Reasoning Manifold (*r*_*t*_ ∈ *N*): The agent’s internal thoughts live on an independent internal reasoning manifold *N* with its own metric *g*_*N*_. This allows the agent’s internal logic, confidence metrics, or latent abstractions to possess hierarchical or hyperbolic geometry (which is ideal for tree-like decision structures), instead of tunneling through it).

#### 3.2.2. The Manifold POMDP Model

A Partially Observable Markov Decision Process (POMDP) models decision-making where the agent cannot directly see the true state of the world. The formulation defines a Manifold POMDP tuple:

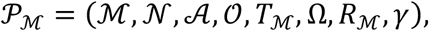

where

**ℳ:** The underlying curved state space of the environment,

***N*:** The structured internal reasoning space of the AI,

***A*:** The set of actions the agent can take,

***O*:** The set of observations the agent can receive,

***T***_**ℳ**_: The transition probability function, mapping how actions move the hidden state across the manifold ℳ,

**Ω:** The observation probability function,

***R***_**ℳ**_: The reward function based on the manifold state,

***γ*:** The discount factor for future rewards.

##### Core State Variables

*s*_*t*_ = *z*_*t*_ ∈ ℳ: The true, hidden physical or environmental state at time *t*.

*r*_*t*_ = *ρ*_*t*_ ∈ *N*: The agent’s current internal thought process or cognitive state.

*b*_*t*_ ∈ *P*(ℳ): The “belief distribution.” Because the agent cannot see *z*_*t*_ directly, it maintains a probability distribution over the manifold ℳ representing where it thinks it is.

#### 3.2.3. The Agentic Reasoning Loop

The agentic reasoning loop (He and Yu 2026) becomes

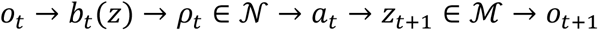

*o*_*t*_: The agent receives an imperfect sensor reading or observation from the world.

*b*_*t*_(*z*): The agent updates its belief distribution over the environmental manifold ℳ using this new observation.

*ρ*_*t*_ ∈ *N*: The agent maps this belief state into its internal reasoning manifold to deliberate, weigh options, or update its memory.

*a*_*t*_: Based on its internal thought state, the agent executes an action.

*z*_*t*+1_ ∈ ℳ: The action forces the environment’s true hidden state to change, moving to a new point on the environment manifold.

*o*_*t*+1_: The cycle repeats as the new state triggers a new observation.

This is the manifold version of the POMDP-based think–act framework. By utilizing manifolds, this formulation ensures that the AI’s planning algorithms respects structural constraints automatically. For example, if a robotic arm can only rotate up to 180 degrees, embedding its state space into a manifold natively prevents the AI from calculating mathematically valid but physically impossible “flat linear shortcuts” that would break the robot.

### 3.3. Manifold belief update

The following formula adapts Bayesian belief filtering to curved, non-Euclidean spaces (Perich et al. 2025). It replaces flat Euclidean state spaces with a Riemannian manifold (ℳ), which is crucial for tracking states with inherent geometric constraints, like rotations or sphere surfaces. Et al. 2026)

The belief is no longer a density over ℝ^*d*^. It is a density over a manifold:

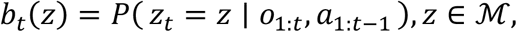

where belief state *b*_*t*_(*z*) is the probability density. It tracks the hidden state *z* on the manifold ℳ. It conditions on past actions (*a*) and observations (*o*).

The belief update is

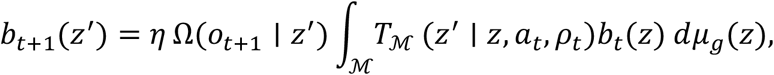

where

**Observation Likelihood Ω**: The measurement model. It evaluates how likely observation *o*_*t*+1_ is from state *z*′.

**Transition Dynamics *T*_ℳ_**: The motion model. It calculates the probability of moving from *z* to *z*′ via action *a*_*t*_.

**Volume Measure *dμ*_*g*_(*z*)**: The Riemannian volume element. It adjusts for space curvature using the metric tensor *g*.

**Normalization *η***: A scaling constant. It ensures total probability sums to one.

#### What Changes from Standard Filtering?

The key change is that integration is now over the manifold, not over a flat Euclidean space.

##### Integration Space

The integral operates strictly over ℳ. Standard filters integrate over flatℝ^*d*^.

##### Volume Distortion

Straight-line distances do not apply here. The measure *dμ*_*g*_(*z*) scales probabilities based on local curvature.

##### Motion Constraints

Dynamics follow valid paths along the surface. Particles cannot “fly off” the manifold during updates.

### 3.4. Manifold transition model

The manifold transition model extends standard Euclidean state transitions to curved surfaces (manifolds) by generating a change vector in the local flat tangent space and projecting it back onto the manifold using the exponential map.

Here is the step-by-step breakdown of its mathematical formulation.

#### 3.4.1. From Euclidean to Curved Space

In a standard Euclidean Partially Observable Markov Decision Process (POMDP), the next state *s*_*t*+1_ is calculated using simple vector addition:

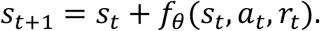

On a curved manifold ℳ, you cannot simply add a vector to a point because straight lines do not stay on the surface (e.g., adding a straight vector to a point on a sphere pushes it into empty space). Instead, the model splits the transition into two distinct steps:

1. Predict a velocity vector in the flat tangent space 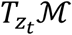 at the current point *z*_*t*_.
2. Wrap that vector onto the manifold along a locally straight path (a geodesic).

#### 3.4.2. The Tangent Vector Prediction

The function *f*_*θ*_ acts as the transition dynamics neural network:

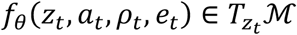

***z***_***t***_: The current state on the manifold ℳ.

***a***_***t***_: The action taken by the agent.

***ρ***_***t***_, ***e***_***t***_: Reward or environmental noise variables.

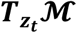: The tangent space, a localized flat vector space touching the manifold exactly at *z*_*t*_. The output of *f*_*θ*_ is not the next state itself, but a directional vector (velocity) rooted at *z*_*t*_.

#### 3.4.3. The Exponential Map Projection

To find the actual next state *z*_*t*+1_ on the manifold, the model applies the exponential map 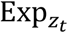:

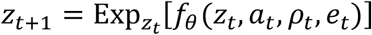

The operator 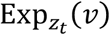 takes the tangent vector *v*, moves along the manifold in that direction along a geodesic (the shortest path on a curved surface) for a distance equal to the length of *v*, and outputs the landing point on ℳ.

#### 3.4.4. The Manifold Gaussian Transition Density

To account for uncertainty, the model defines a probability distribution over the next state *z*′ given the current state *z*, action *a*, and parameters *ρ*. The transition density can be written as

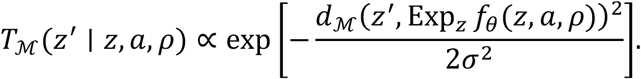

∝: Represents proportionality, meaning a normalization constant (the Riemannian volume element) is omitted.

**Exp**_***z***_***f***_***θ***_(***z, a, ρ***): The predicted mean next state on the manifold.

***d***_**ℳ**_(⋅,⋅): The geodesic distance function, measuring the shortest path along the curved surface between the true next state *z*′ and the predicted mean state.

***σ***^**2**^: The variance parameter controlling transition noise.

This equation is the Riemannian generalization of a standard Gaussian distribution (often called a Riemannian Gaussian or Von Mises-Fisher-like generalization), where Euclidean distance ‖*x* − *μ*‖^2^ is swapped for squared geodesic distance *d*_ℳ_(*z*′, *μ*)^2^.

In summary, the manifold transition model replaces flat coordinate shifting with a two-step geometric pipeline: it evaluates neural network dynamics in a local flat vector space, then maps those vectors back to the true curved geometry via geodesics to calculate valid state probabilities.

### 3.5. Internal reasoning variable on a manifold

In this section, we introduce internal reasoning variable on a manifold which models the process of logical thinking not as a series of discrete, flat vector operations, but as a continuous geometric trajectory on a smooth Riemannian manifold *N*, where thoughts evolve along curved paths via tangent vectors and exponential maps.

#### 3.5.1. The Reasoning Space (*ρ*_***t***_ ∈ *N*)

Instead of representing a “thought state” as a standard vector in flat Euclidean space ℝ^*d*^, this formulation places it on a reasoning manifold *N*.

##### Geometric constraints

Spaces like a hypothesis manifold or causal-graph manifold have strict topological rules (e.g., probabilities must sum to 1, graphs must be directed and acyclic).

##### Curvature

A flat space assumes moving from point A to point B is always a straight line. On a manifold, the “shortest path” between two logical ideas is a curved line called a geodesic, which respects the underlying structure of the knowledge domain.

#### 3.5.2. The Thought Vector Space 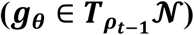

The function *g*_*θ*_(*ρ*_*t*−1_, *b*_*t*_, *o*_*t*_, *m*_*t*_) represents the mental update or “force” generated by a neural network with parameters *θ*. It takes the previous thought state *ρ*_*t*−1_, current beliefs *b*_*t*_, observations *o*_*t*_, and memory *m*_*t*_ to compute the next reasoning direction.

##### Tangent space

This update vector does not live on the manifold itself; it lives in the tangent space 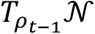.

##### Local flat boundary

The tangent space is a localized, completely flat vector space tangent to the manifold at the exact point of the current thought *ρ*_*t*−1_. This allows the neural network to output standard flat vectors (*g*_*θ*_) before they are projected onto the curved manifold.

This is important because reasoning itself may not be flat. For example, reasoning trajectories may lie on structured spaces such as:

hypothesis manifolds, proof-state manifolds, causal-graph manifolds, tool-use policy manifolds, biological mechanism.

#### 3.5.3. Walking the Curve via the Exponential Map 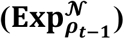

Because you cannot simply add a flat vector to a point on a curved surface (*ρ*_*t*−1_ + *g*_*θ*_) would shoot off the manifold into empty space), the formulation uses the Exponential Map 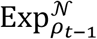:

A reasoning update is

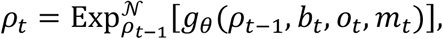

where

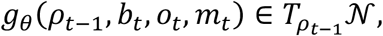

##### Manifold projection

The exponential map takes the flat update vector *g*_*θ*_ from the tangent space and “rolls” it safely onto the curved surface of the manifold,

##### Geodesic stepping

It traces a geodesic curve starting at *ρ*_*t*−1_ in the direction of *g*_*θ*_(*ρ*_*t*−1_, *b*_*t*_, *o*_*t*_, *m*_*t*_) for a distance equal to the length of*g*_*θ*_, landing exactly on the new valid thought state *ρ*_*t*_.

So thinking itself becomes a trajectory on a reasoning manifold

Figure 13 shows geometric reasoning on a manifold.

**Figure 13.**
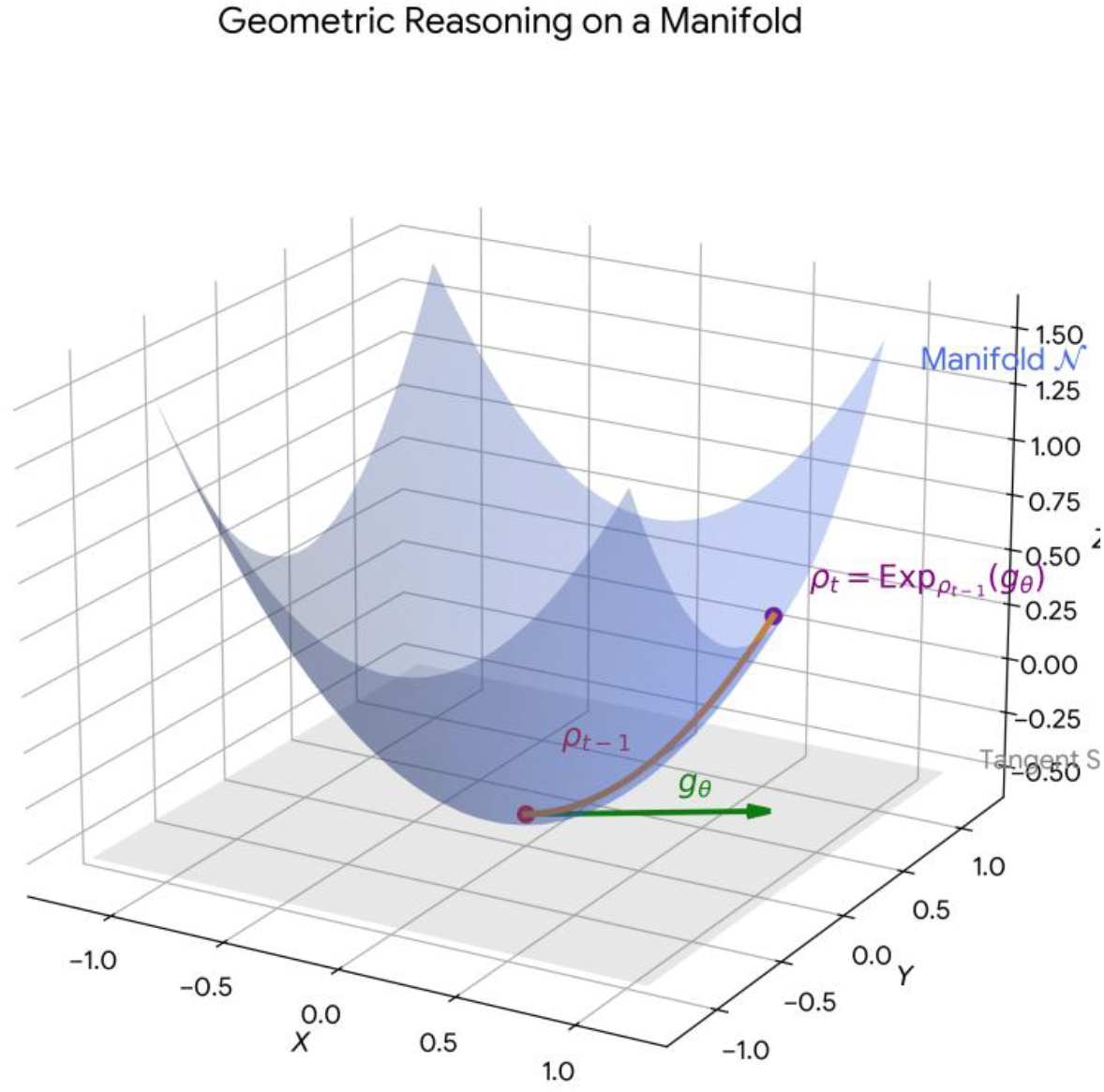
Geometric reasoning on a manifold.

#### 3.5.4. Thinking as a Continuous Trajectory

By chaining these steps together, the model transforms the act of “thinking through a complex problem” into differential geometry. A long sequence of reasoning steps forms a continuous piecewise geodesic trajectory across the manifold. For instance, solving a math problem is represented as a path traveling through a proof-state manifold, where every twist and turn of the trajectory corresponds to a geometric correction dictated by the logical structure of mathematics. To help readers to easily understand that thinking can be taken as a continuous trajectory, we use *x*^2^ − 5*x* + 6 = 0 as an example to plot Figure 14 to visualize thinking as a continuous trajectory.

**Figure 14.**
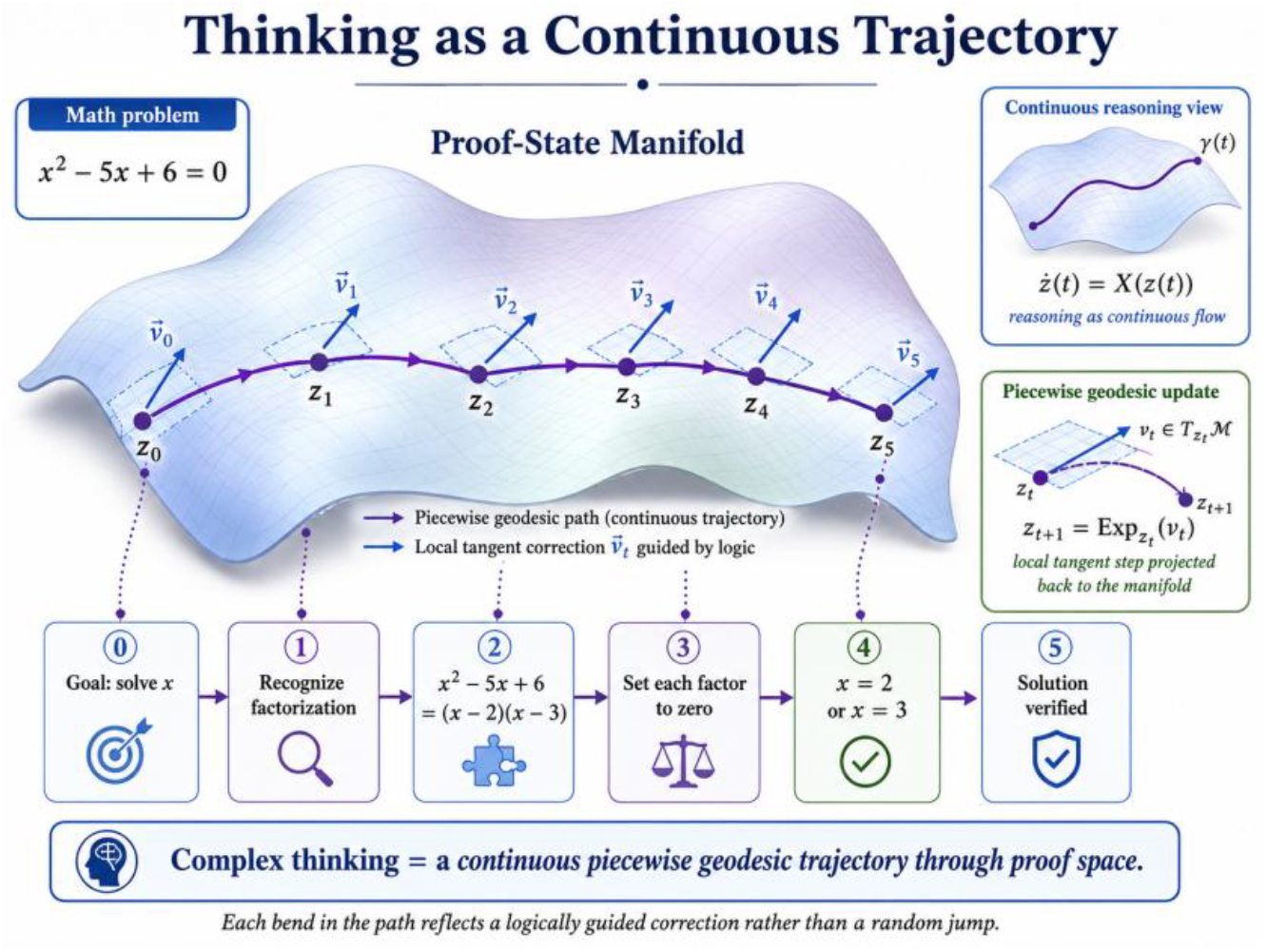
Thinking as a continuous trajectory.

### 3.6. Think–act decomposition on manifolds

We will introduce “think-act decomposition on manifold”. This framework represents a Differential Geometric formulation of a Partially Observable Markov Decision Process (POMDP) containing internal reasoning. It models an agent whose thinking states, physical actions, and environmental states update along smooth mathematical surfaces (manifolds) rather than flat Euclidean spaces.

#### 3.6.1. Variables and Components

The policy becomes

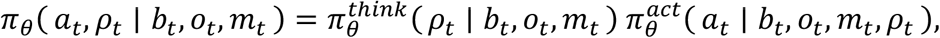

where

***b***_***t***_: Belief state at time *t* (the agent’s internal probability distribution over hidden environmental states).

***o***_***t***_: Observation received from the environment at time *t*.

***m***_***t***_: Memory state at time *t* tracking long-term contextual historical information.

***ρ***_***t***_: Internal thinking state at time *t* that lives on a smooth Riemannian thinking manifold *N*(*ρ*_*t*_ ∈ *N*).

***a***_***t***_: Action taken at time *t*. It exists in the tangent space of the environmental state manifold ℳ at the current state 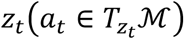.

***z***_***t***_: Geometric environmental or agent state at time *t* residing on a state manifold ℳ(*z*_*t*_ ∈ ℳ).

#### 3.6.2. Step-by-Step Execution Flow

##### Step 1: Internal Thinking Update

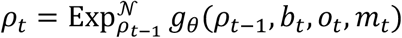

###### Tangent Vector Generation

The neural network function *g*_*θ*_ processes the previous thought state *ρ*_*t*−1_, current belief *b*_*t*_, observation *o*_*t*_, and memory *m*_*t*_. It outputs a flat vector belonging to the tangent space of the manifold *N* at position *ρ*_*t*−1_.

###### Exponential Mapping

The operator 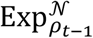 takes this flat tangent vector and projects it onto the curved geometry of the thinking manifold *N*. This ensures the updated thought state *ρ*_*t*_ naturally obeys the geometric constraints of the manifold space (e.g., maintaining unit length on a sphere or remaining positive-definite).

**The manifold think policy** is

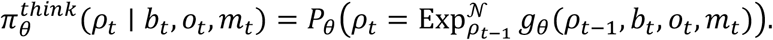

##### Step 2: Action Selection

The action policy is

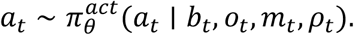

###### Conditioned Generation

The physical action policy 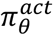 calculates the next move. Critically, its output distribution is explicitly conditioned on the fresh thinking state *ρ*_*t*_ generated in Step 1. This formalizes a “think before you act” pipeline.

##### Step 3: Geometric State Transition

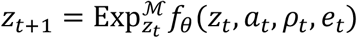

###### Manifold Physics Dynamics

If the agent operates a system bound by complex non-Euclidean constraints (e.g., camera rotations, robotic joints, or aerodynamic configurations), the next physical state *z*_*t*+1_ cannot be derived via standard flat vector addition (*z*_*t*_ + *a*_*t*_).

###### Geometric Projection

The system transition function *f*_*θ*_ processes the current state *z*_*t*_, the action *a*_*t*_, the thought *ρ*_*t*_, and any environmental noise *e*_*t*_ to compute a geometric displacement vector. The exponential map 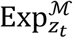 maps this vector along a curved path (geodesic) on the state manifold ℳ to safely land at the legitimate physical state configuration *z*_*t*+1_.

#### 3.6.3. Core Policy Decomposition

Standard flat policy networks map directly from a history tuple to an action: *π*(*a*_*t*_|history). This framework splits that monolithic policy into two sequentially dependent processes:

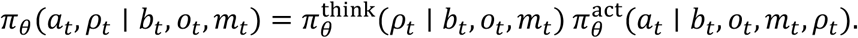

By enforcing this factorization, the agent is mathematically barred from acting reflexively; it must pass its observations through an explicit latent reasoning layer *π*^think^ before deciding how to manipulate its environment *π*^*act*^.

**In summary**, the “Think–Act Decomposition on Manifolds” provides a rigorous mathematical bridge between internal latent reasoning (Chain-of-Thought style processes) and non-Euclidean physical dynamics (robotics, orientation physics, and non-linear tracking structures). It wraps a standard internal-reasoning POMDP in a differential geometry framework, utilizing exponential mapping functions to guarantee that both the agent’s thoughts and physical states always evolve within their respective valid geometric boundaries.

So the complete manifold think–act policy is

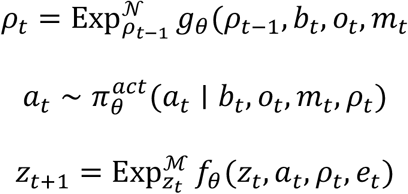

This is the direct manifold extension of the paper’s internal-reasoning POMDP.

To intuitively understand why we need think-act decomposition on manifolds for improving reasoning, we plot Figure 15 that vitalizes the structure of think-act decomposition on manifolds and their decomposition.

**Figure 15.**
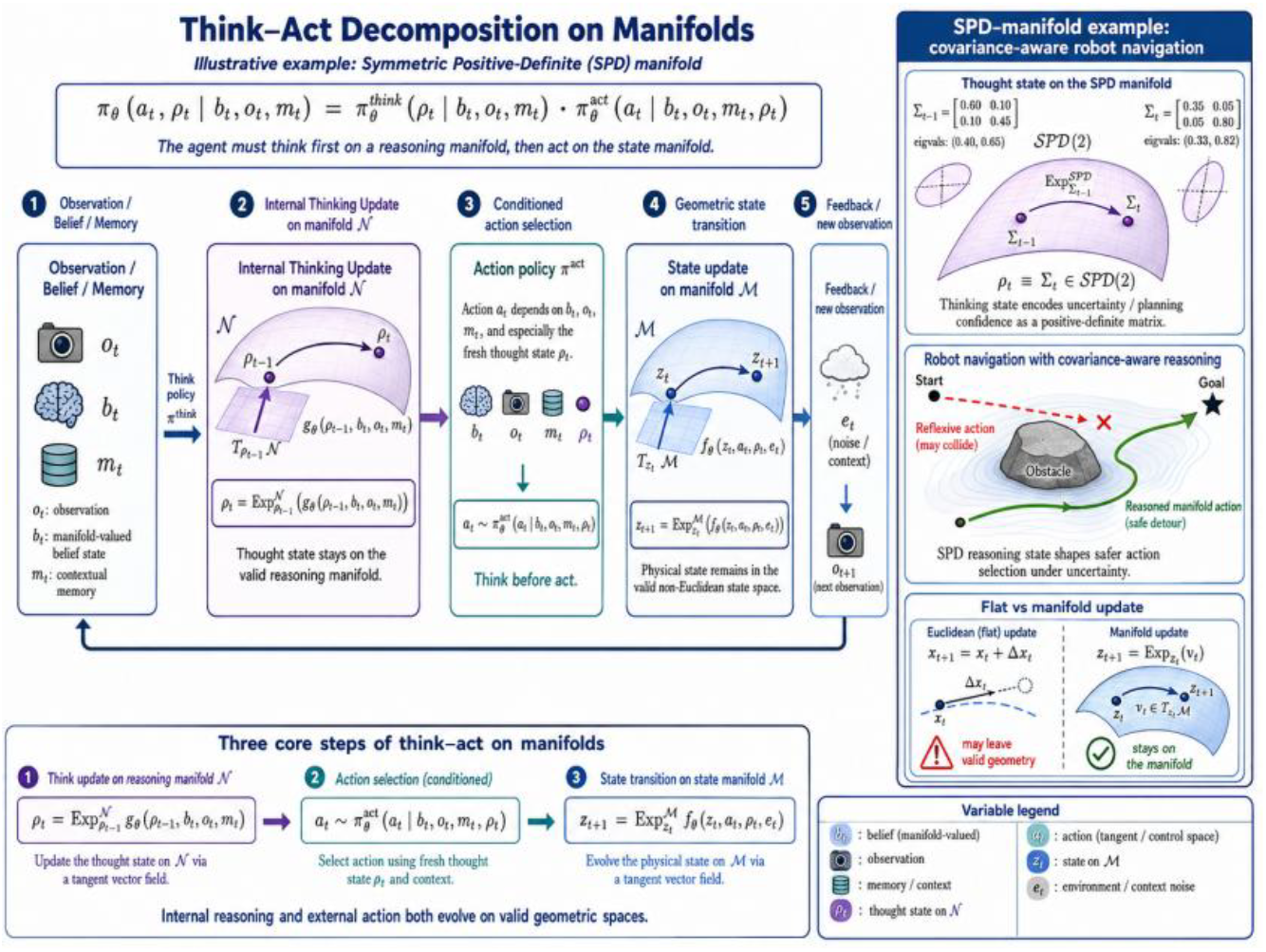
Think-act decomposition on manifolds.

### 3.7. Post-training reasoning with reinforcement learning on manifolds

We distinguish in-context reasoning from post-training reasoning, where post-training uses RL or SFT to optimize agent behavior. The following formulation extends Post-Training Reinforcement Learning (RL) into the domain of differential geometry (manifolds) to optimize how an AI agent thinks and acts. Instead of treating reasoning steps as arbitrary text tokens, this framework models the agent’s hidden thought trajectories (*ρ*_*t*_) and environmental states (*z*_*t*_) as moving along continuous Riemannian manifolds.

Here is the detailed breakdown of the mathematical framework.

#### 3.7.1. The Core Objective Function

In the manifold version, the goal is to optimize the expected cumulative discounted reward over a manifold trajectory:

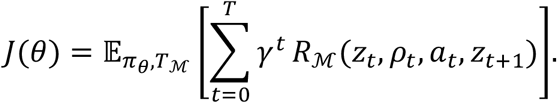

***θ***: The learnable parameters (weights) of the neural network policy.

***π***_***θ***_: The agent’s policy, which dictates both internal thought generation and external action selection.

***T***_**ℳ**_: The transition dynamics governing how states evolve on the manifold.

***γ*** ∈ [**0, 1**]: The temporal discount factor that balances immediate rewards against long-term success.

***R***_**ℳ**_: The manifold-constrained reward function evaluating the quality of a state transition given an internal reasoning state.

#### 3.7.2. The Multi-Objective Reward Decomposition

The global reward *R*_ℳ_ linearly balances performance, logical coherence, and geometric constraints. The reward can include task success, geometric validity, causal validity, verification score, and reasoning efficiency:

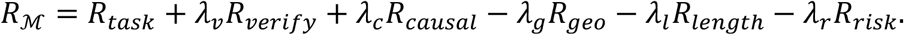

The scalar coefficients *λ* act as hyperparameters to weight each constraint.

##### Geometric Invariance Penalty (*R*_*geo*_)

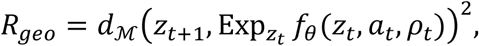

###### Concept

Measures how much the actual next state *z*_*t*+1_ deviates from the agent’s forward dynamics model *f*_*θ*_ projected onto the manifold.

###### Geometry

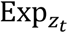 is the Exponential Map. It takes a tangent vector generated by the neural network *f*_*θ*_ and projects it back down onto the curved manifold surface along a geodesic (the shortest curved path).

###### Purpose

***d***_**ℳ**_(⋅,⋅)^**2**^ is the squared Riemannian distance. Minimizing this forces the agent’s actions and thoughts to respect the physical or structural laws of the manifold.

##### Automated Verification Score (*R*_*verify*_)

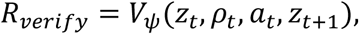

###### Concept

Evaluates the mathematical or logical correctness of a transition.

###### Implementation

A critique or verifier model *V*_*ψ*_ evaluates whether the thought process *ρ*_*t*_ yields a valid action *a*_*t*_ changing the state to *z*_*t*+1_.

##### Reasoning Efficiency Penalty (*R*_*length*_)

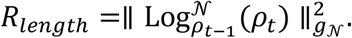

###### Concept

Penalizes long, winding, or repetitive thinking loops.

###### Geometry

Log^*N*^ is the Logarithmic Map (the inverse of the exponential map) on the internal reasoning manifold *N*. It maps the step from the last thought *ρ*_*t*−1_ to the current thought*ρ*_*t*_ into a flat tangent space vector.

###### Purpose

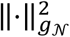 calculates the squared length of this vector using the local Riemannian metric tensor *g*_*N*_. This serves as a geometric regularizer: instead of counting raw words or tokens, it penalizes the true “informational distance” traversed in the latent thought space.

The last term penalizes unnecessarily long internal reasoning trajectories.

#### 3.7.3. Policy Decomposition: Co-Distinguishing Thought and Action

The fundamental trick of this framework is splitting a single token-generation pipeline into two distinct components: a internal latent deliberation channel and an external execution channel. Given the agent state (belief *b*_*t*_), observation *o*_*t*_, memory *m*_*t*_, the complete policy factorizes cleanly:

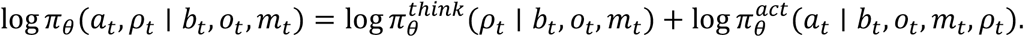

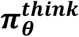: Controls the latent mental state *ρ*_*t*_ (the “Chain of Thought” or internal planning trajectory) conditioned on what the agent currently knows.

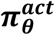: Selects the environmental execution *a*_*t*_, conditioned on both the baseline environment state and the specific path chosen by the internal thinking step *ρ*_*t*_.

#### 3.7.4. Policy Gradient Optimization

The RL objective is

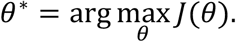

To maximize *J*(*θ*), the framework takes the derivative of the objective function with respect to the network weights. Applying the classic log-derivative trick yields:

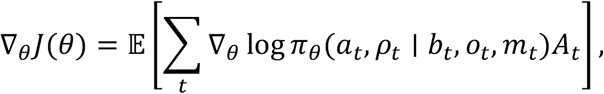

Where

Where the Advantage Function *A*_*t*_ calculates how much better the chosen thought-action pair performs compared to the baseline average:

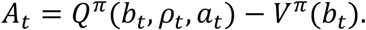

Because the policy decomposes into thinking and acting,

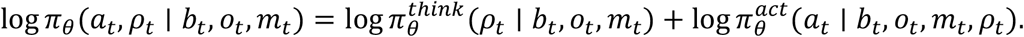

Substituting the policy factorization into the gradient formula distributes the update directly into both sub-policies. Therefore,

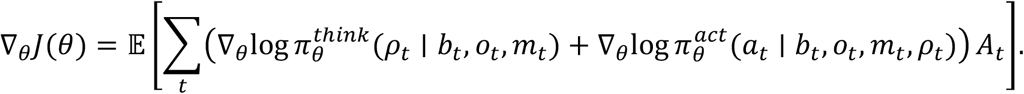

This gives RL for both:

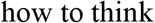

and

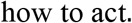

### 3.8. Manifold PPO-style post-training

The mathematical formulation of Manifold PPO-style post-training generalizes Proximal Policy Optimization (PPO) to large language models (LLMs) that execute complex internal reasoning (“thinking”) before generating an answer (“acting”), while constraining their latent transitions to smooth geometric paths (manifolds).

Here is the step-by-step breakdown of its components, structured from its foundational probability decomposition to the final composite loss function.

#### 3.8.1. Think–Act Decomposition

Standard reinforcement learning models a direct policy *π*_*θ*_(*a*|*s*). For advanced reasoning models, the policy is split into a “think” phase (internal chain-of-thought, reasoning steps, or latent path selection) and an “act” phase (generating the final visible token or action).

Given:

*b*_*t*_: Background context / history

*o*_*t*_: Current observation

*m*_*t*_: Memory stateType equation here.

*ρ*_*t*_: The “think” or reasoning variable at step t

*a*_*t*_: The final “act” or output action at step *t*

The total policy probability is factored as:

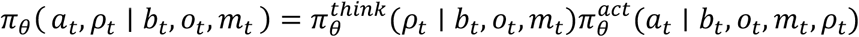

##### Probability Ratio Split

In PPO, we track the ratio of the new policy *π*_*θ*_ to the old policy 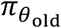. Because of the product rule above, the total ratio *r*_*t*_(*θ*) splits cleanly into a product of the thinking ratio and the acting ratio:

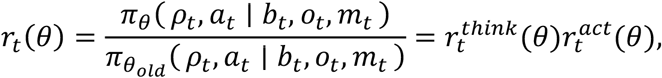

where

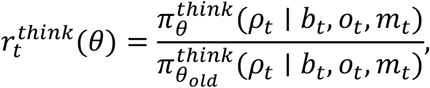

and

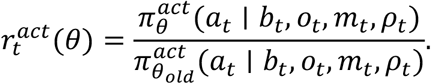

#### 3.8.2. The PPO Objective

The core reinforcement learning objective uses the combined ratio *r*_*t*_(*θ*) multiplied by the generalized advantage estimator *A*_*t*_ (which measures how much better this specific trajectory is compared to the average expected reward).To prevent the updated policy from changing too drastically from the old policy, the objective clips the ratio within a narrow window [1 − *ϵ*, 1 + *ϵ*]:

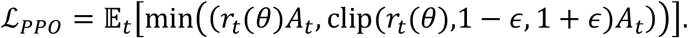

Because 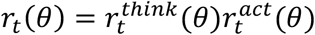, a massive deviation in either the internal reasoning sequence or the final text output will trigger the clipping mechanism, stabilizing joint training.

#### 3.8.3. Geometric Regularization (ℒ_*geoRL*_)

The unique aspect of Manifold PPO is the geometric penalty ℒ_*geoRL*_. Instead of assuming hidden states or reasoning paths exist in flat Euclidean space, it assumes they live on Riemannian manifolds (*N* and ℳ) with specific metrics (*g*_*N*_ and *g*_ℳ_).

For manifold reasoning, we add geometric regularization:

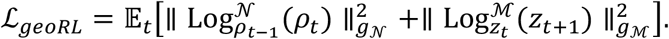

##### KEY Concepts

###### Logarithmic Map (log_*p*_(*q*))

In Riemannian geometry, the log map takes two points on a curved manifold (*p* and *q*) and maps them to a vector in the flat tangent space at point *p*. The length of this vector is exactly equal to the geodesic (shortest path) distance between the two points on the curved surface.

###### Geodesic Distance Minimization

Minimizing the squared norm 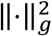 penalizes the model if its successive reasoning steps (*ρ*_*t*−1_ → *ρ*_*t*_) or latent states (*z*_*t*_ → *z*_*t*+1_) make violent, unnatural jumps.

###### Purpose

It forces the LLM’s internal thinking process to progress along a smooth, logically continuous topological path, preventing erratic “hallucinatory” leaps in logic.

#### 3.8.4. Total Post-Training Objective

The final comprehensive loss function ℒ_*post*_ is a multi-objective optimization problem. It balances maximum reward, value function tracking, geometric smoothing, answer verification, and distribution stability.

The total post-training objective is defined as

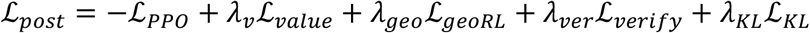

where 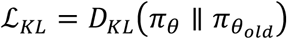 prevents excessive policy drift.

Each component in ℒ_*post*_ serves a distinct role (weighted by their respective hyperparameter *λ*:

−***ℒ***_***PPO***_ **(Policy Gradient):** We minimize the negative PPO loss, which is mathematically equivalent to maximizing the expected advantage/rewards.

***ℒ***_***value***_ **(Critic Loss):** Minimizes the mean squared error of the value network, ensuring the model accurately predicts the expected long-term rewards of current states.

***ℒ***_***geoRL***_ **(Manifold Regularization):** Smooths the latent trajectories of internal thought processes as detailed above.

***ℒ***_***verify***_ **(Verification Loss):** An objective (often a classification or rule-based loss) that evaluates final outcome accuracy (e.g., verifying if a math answer or block of code compiles and runs correctly).

***ℒ***_***KL***_ **(Kullback-Leibler Divergence):** Computed as 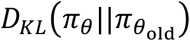, it prevents the model’s overall token probability distribution from collapsing or drifting too far away from the reference base model, preserving conversational fluency.

A more geometric KL can be decomposed as

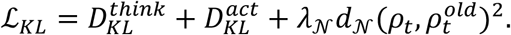

**In summary**, by decomposing policies into explicit thinking and acting ratios and calculating distances via Riemannian Log maps, Manifold PPO guides models to not only find correct answers but also construct structurally stable internal reasoning pathways.

### 3.9. Value function on belief and reasoning manifold

This mathematical framework represents an advanced extension of Partially Observable Markov Decision Processes (POMDPs). It explicitly models not just the agent’s uncertainty about the external world (the belief state), but also the computational and structural state of its own thinking process (the reasoning state).

Here is the detailed breakdown of the components, mechanics, and geometric implications of this formulation.

#### 3.9.1. The State Space: Belief and Reasoning Manifolds

In standard reinforcement learning, a value function depends only on a physical state *s*. In POMDPs, it depends on a belief state *b*, which is a probability distribution over hidden world states (*b*(*s*) = *P*(*s*)).

This formulation introduces a dual-state representation:

Belief State (*b*_*t*_): A point on a belief manifold. It tracks environmental uncertainty based on historical observations.

##### Internal Reasoning State (*ρ*_*t*_)

A point on a reasoning manifold. This represents the agent’s latent cognitive state, such as current active hypotheses, deep-thinking tokens, memory states, or computational traces (e.g., in a reasoning LLM or Monte Carlo Tree Search).

Thus, the value function *V*^*π*^(*b*_*t*_, *ρ*_*t*_) evaluates the expected long-term reward given what the agent knows about the world (*b*_*t*_) paired with how the agent is currently processing that information (*ρ*_*t*_).

#### 3.9.2. The Discrete-Time Bellman Equation

The discrete-time Bellman equation formalizes how the value propagates step-by-step:

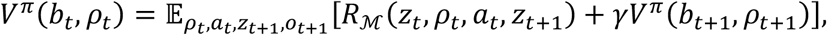

where the value function *V*^*π*^(*b*_*t*_, *ρ*_*t*_) should depend on both the belief state and the internal reasoning state.

##### Breaking down the Expectation Operator (*E*)

The expectation is taken over multiple sources of randomness and transitions:

*a*_*t*_~*π*(⋅ |*b*_*t*_, *ρ*_*t*_): The action chosen by policy *π*.

*z*_*t*_, *z*_*t*+1_: The true hidden world states at times *t* and *t* + 1.

*o*_*t*+1_: The new observation received from the environment after taking action *a*_*t*_.

##### The Components

***R***_**ℳ**_(***z***_***t***_, ***ρ***_***t***_, ***a***_***t***_, ***z***_***t***+**1**_): The meta-reward function. Crucially, it depends on the hidden world transition (*z*_*t*_ → *z*_*t*+1_) and the internal reasoning state *ρ*_*t*_. This allows the system to reward or penalize the agent for the quality or cost of its reasoning (e.g., penalizing long inference times or ***γ***: The discount factor (0 ≤ *γ* < 1), weighting future rewards against immediate ones.

***b***_***t***+**1**_: The updated belief state, calculated via a filtering update (like a Bayes filter) incorporating (*b*_*t*_, *a*_*t*_) and *o*_*t*+1_.

*ρ*_*t*+1_: The updated reasoning state, tracking how the internal thought process evolves after making a decision and observing the outcome.

The Optimal Value Function (*V*^∗^) simply solves this by taking the supremum/maximum over all valid policies *π*:

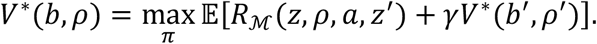

This is a **manifold belief-reasoning Bellman equation**.

#### 3.9.3. The Continuous-Time Realization: The HJB Equation

When reasoning and environment transitions occur continuously rather than in distinct steps, the system transitions from a Bellman equation to a Hamilton-Jacobi-Bellman (HJB) partial differential equation:

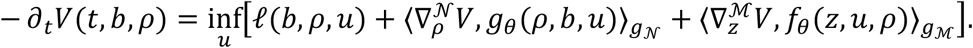

This couples the geometry of the world state and the geometry of the reasoning state.

##### Deconstructing the Continuous Operators

−**∂**_***t***_***V***(***t, b, ρ***): The time-derivative of the value function, representing how the total remaining value changes over time.

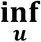: The optimal control input *u* (analogous to action *a*)) chosen to optimize the trajectory.

***ℓ***(***b, ρ, u***): The instantaneous cost/loss rate (the continuous version of the negative reward).

##### The Geometric Coupling (Inner Products)

The core of this equation lies in the two inner product terms, which utilize Riemannian geometry metrics (*g*_*N*_) and *g*_ℳ_) to project value gradients along the tangent spaces of the manifolds:

###### 1. Reasoning Dynamics 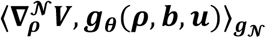

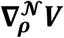 is the gradient of the value function with respect to the reasoning manifold *N*.

***g***_***θ***_(***ρ, b, u***) is a vector field defining how the internal thought state changes continuously over time.

**The inner product** 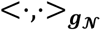 measures how much the value function changes along the trajectory of the reasoning state, scaled by the metric tensor *g*_*N*_ of the reasoning manifold.

###### 2. World Dynamics 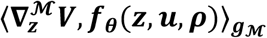

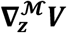 is the gradient of the value function with respect to the latent world state manifold ℳ.

***f***_***θ***_(***z, u, ρ***) represents the physical system dynamics of the hidden state.

**The inner product** 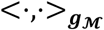 measures how much value changes as the external world evolves, scaled by the world manifold’s metric tensor *g*_ℳ_.

##### Summary of the Manifold Coupling

What makes this formulation unique is that *g*_*θ*_ depends on *b*, and *f*_*θ*_ depends on *ρ*.

###### World-to-Reasoning Influence (*b* → *g*_*θ*_)

Your current belief about the world changes how your internal thoughts evolve. For example, high uncertainty in *b* might trigger a sharp shift in *ρ* toward active exploration or deeper information gathering.

###### Reasoning-to-World Influence (*ρ* → *f*_*θ*_)

Your internal cognitive state influences how actions manipulate the external state or how you perceive it.

By binding these together via Riemannian metrics ((*g*_*N*_, *g*_ℳ_)), the framework proves that optimal decision-making requires navigating a unified geometry where thinking about the world and interacting with the world are mathematically inseparable.

### 3.10. Verification-gated think–act reasoning

A key improvement is to place a verifier between thinking and acting. The mathematical framework below describes a Verification-Gated Think–Act Loop used in advanced AI agents.

It ensures that an agent does not execute an action in the real world unless its internal reasoning and predicted outcomes pass a strict safety and utility gate.

Here is the step-by-step breakdown of the mathematical formulation and universal concepts.

#### 3.10.1. Thinking Phase (Reasoning Generation)

The agent generates a reasoning state:

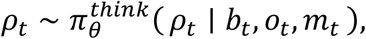

where:

***ρ***_***t***_ **(Reasoning State):** The internal chain-of-thought, hidden rationale, or mental simulation generated by the agent at time step *t*.

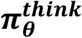: The policy parameterized by weights *θ* that generates thoughts.

##### Conditioning Variables

The thought is based on the agent’s belief state (*b*_*t*_), current observation (*o*_*t*_), and the main mission or task goal (*m*_*t*_).

#### 3.10.2. Acting Phase (Action Proposal)

Then it proposes an action:

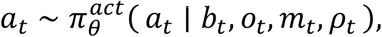

where:

***a***_***t***_ **(Proposed Action):** The concrete command or tool call the agent wants to execute.

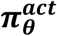: The acting policy.

##### Crucial Difference

Unlike standard agents, this policy is strictly conditioned on *ρ*_*t*_ (the thought generated in Step 1), ensuring actions are backed by explicit reasoning.

#### 3.10.3. Prediction Phase (World Modeling)

The verifier checks the tuple:

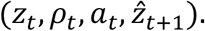

The predicted next state is

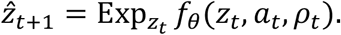

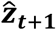 **(Predicted Next State):** The agent’s internal prediction of how the environment will change after taking action *a*_*t*_.

***f***_***θ***_: A dynamics model (or world model) mapping the current latent state *z*_*t*_, proposed action *a*_*t*_, and reasoning *ρ*_*t*_ to a tangent vector.

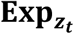 **(Exponential Map):** A term from differential geometry. It means the state space is treated as a curved manifold rather than flat space. The function *f*_*θ*_ calculates a “direction of change,” and the exponential map projects it back onto the valid state space starting from *z*_*t*_.

#### 3.10.4. Verification Phase (Evaluation)

The verification score is

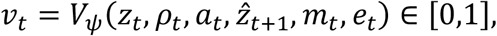

where:

***v***_***t***_ **(Verification Score):** A scalar score between 0 (completely invalid/dangerous) and 1 (perfectly sound).

***V***_***ψ***_: A separate verification neural network parameterized by *ψ*.

##### Inputs

It cross-checks everything: current state (*z*_*t*_), the logic (*ρ*_*t*_), the action (*a*_*t*_), the predicted outcome 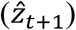, the mission (*m*_*t*_), and environmental constraints or explicit safety rules (*e*_*t*_).

#### 3.10.5. Gating Phase (The Commitment Gate)

The agent decides whether to commit to the action using a multi-objective utility function squeezed through a sigmoid function *σ*:

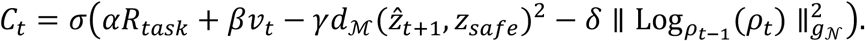

The gate balances four distinct factors using tuning weights (*α, β, γ, δ*):

***αR***_***task***_ **(Reward):** Expected progress toward finishing the task.

+***βv***_***t***_ **(Logical Validity):** The verification score from Step 4.

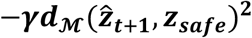 **(Safety Distance):** Penalizes the action if the predicted next state 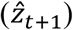 is far from the designated safe state zone (*z*_*safe*_), measured using a manifold metric distance ***d***_**ℳ**_.

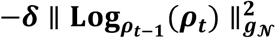 **(Cognitive Inertia/Consistency):** Penalizes wild, erratic shifts in thinking. It uses a Logarithmic map (Log) and a Riemannian metric (*g*_*N*_) to measure the “distance” or velocity between the previous thought (*ρ*_*t*−1_) and the current thought (*ρ*_*t*_). It prevents the agent from completely contradicting its own ongoing logic.

#### 3.10.6. Execution or Rethinking (The Gating Mechanism)

The action *a*_*t*_ is accepted if

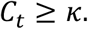

##### Otherwise (*C*_*t*_ < *κ*, threshold failed)

The action is blocked. The agent is forced to rethink and update its reasoning state using a correction model *h*_*ψ*_:

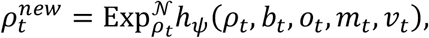

the function *h*_*ψ*_ takes the failed thought and the low verification score (*v*_*t*_) as criticism, calculating a correction step to update the reasoning manifold 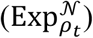. The loop restarts at Step 2 (3.10.2) with this refined 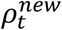 until an action passes the gate.

This gives:

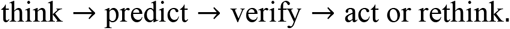

We plot Figure 16 to intuitively visualize Verification-gated think–act reasoning.

**Figure 16.**
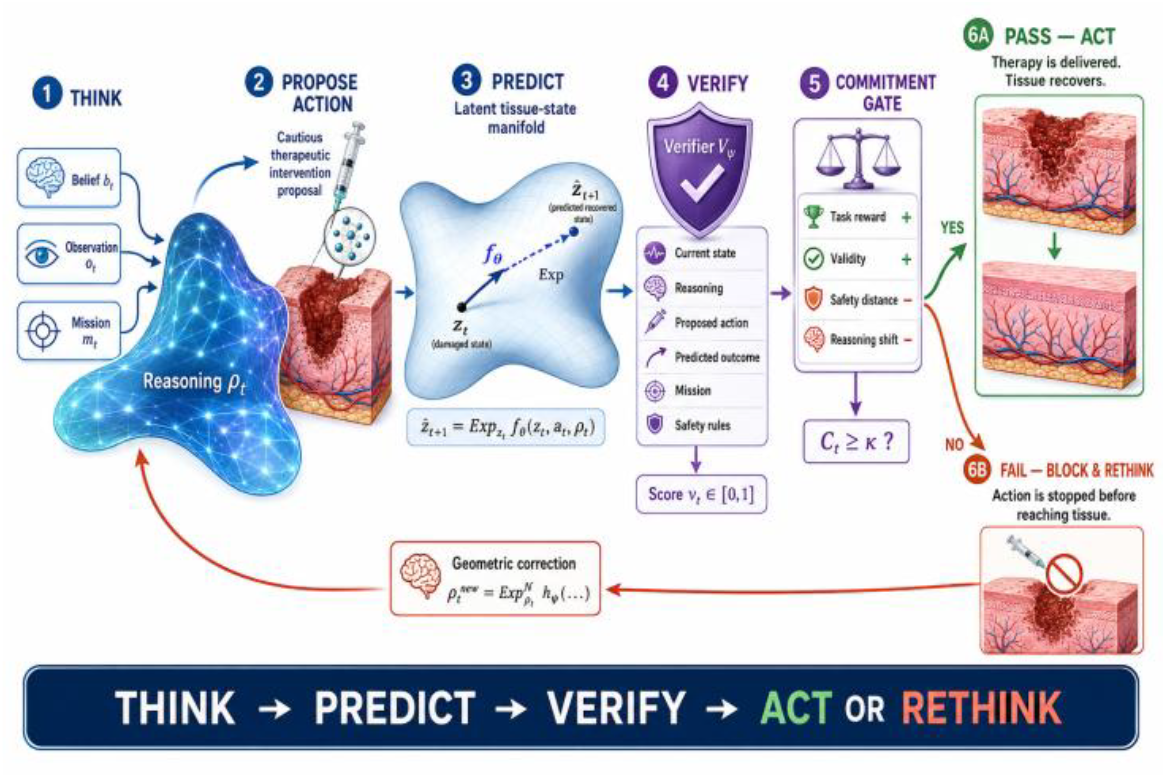
Verification-gated think–act reasoning.

### 3.11. Manifold post-training with process reward

For LLM agents, post-training should not only reward final answers. It should also reward valid intermediate reasoning. The total post-training reward *R*_total_(*τ*) optimizes LLM agents on a geometric reasoning manifold by balancing final correctness, step-by-step progress, logical consistency, and risk reduction.

Here is the step-by-step mathematical breakdown of the framework.

#### 3.11.1. Formulate Trajectory Representation

The trajectory *τ* models the agent’s step-by-step interactions:

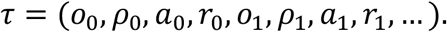

***o***_***t***_ **(Observation):** The environment state or prompt context at step *t*.

***z***_***t***_ **(Latent Hidden State):** The internalized abstract concept or representation of the problem context.

***ρ***_***t***_ **(Position on Reason Manifold):** The current coordinates or embedding vector mapping the agent’s logical state on the smooth mathematical manifold *N*.

***a***_***t***_**(Action):** The generated token, text segment, or reasoning step.\

***r***_***t***_ **(Environment Reward):** Intermediate feedback provided by the environment (if any).

#### 3.11.2. Quantify Outcome Reward

The outcome reward focuses strictly on the destination defined as outcome reward:

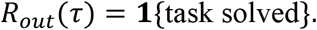

**1**{⋅} **(Indicator Function):** Outputs 1 if the terminal agent response is completely correct, and 0 otherwise.

##### Purpose

Traditional Reinforcement Learning from Human Feedback (RLHF) relies heavily on this binary signal, which lacks granular guidance for complex, multi-step agent reasoning.

#### 3.11.3. Compute Geometric Process Reward

The process reward evaluates the quality of intermediate steps by summing three distinct terms over time:

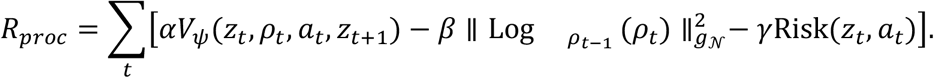

##### Term A: Value-Based Progress (*αV*_*ψ*_)

###### *V*_*ψ*_ (Value Function): An auxiliary model (critic) parameterized by *ψ* that estimates the expected future outcome success from the transition

###### Intuition

Measures the directional forward progress made from state *z*_*t*_ to *z*_*t*+1_ given action *a*_*t*_ at position *ρ*_*t*_. Higher values mean the step moved closer to a correct solution.

##### Term B: Manifold Geodesic Penalty 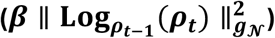

###### Log 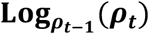 (Riemannian Logarithm Map)

Projects the manifold point *ρ*_*t*_ into the tangent space of the previous point *ρ*_*t*−1_. It computes the velocity vector required to travel between them.

###### 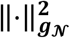 (Riemannian Metric Norm): Calculates the squared length of the shortest path (geodesic) between the two logical points on manifold *N*

###### Intuition

Penalizes massive, erratic jumps or logical disconnects in thought. It acts as a geometric regularization constraint that forces the agent to follow a smooth, continuous, and logically sound chain of thought.

##### Term C: Step Risk Mitigation (*γ* Risk(*z*_*t*_, *a*_*t*_)

###### Risk(*z*_*t*_, *a*_*t*_)

Evaluates the immediate harm, error propagation probability, hallucination score, or safety violations introduced by taking action *a*_*t*_.

###### Intuition

Explicitly penalizes choices that might introduce unrecoverable flaws or unsafe trajectories into the reasoning loop.

#### 3.11.4 Aggregate Total Optimization Reward

The objective combines both components using a scaling hyperparameter *λ*_*proc*_:

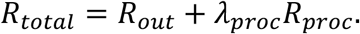

***λ***_***proc***_: Controls the trade-off intensity between achieving final correctness versus adhering to smooth, low-risk, and structured intermediate reasoning pathways.

##### Execute Policy Optimization

The optimal model weights *θ*^∗^ are trained via Policy Gradient or Actor-Critic Reinforcement Learning methods to maximize expected total rewards:

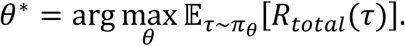

This encourages the agent to learn not merely correct answers, but **valid reasoning trajectories** on the reasoning manifold.

***π***_***θ***_ **(Agent Policy):** The LLM model generating the trajectory actions based on parameters *θ*. **Final Impact:** Rather than just guessing the right final answer through shortcuts, the LLM agent is explicitly driven to navigate along valid structural trajectories inside the geometric reasoning manifold.

To illustrate how to compute reward valid intermediate reasoning and incorporate it into the total reward, we plot Figure 17 to visualize manifold post-training with process reward.

**Figure 17.**
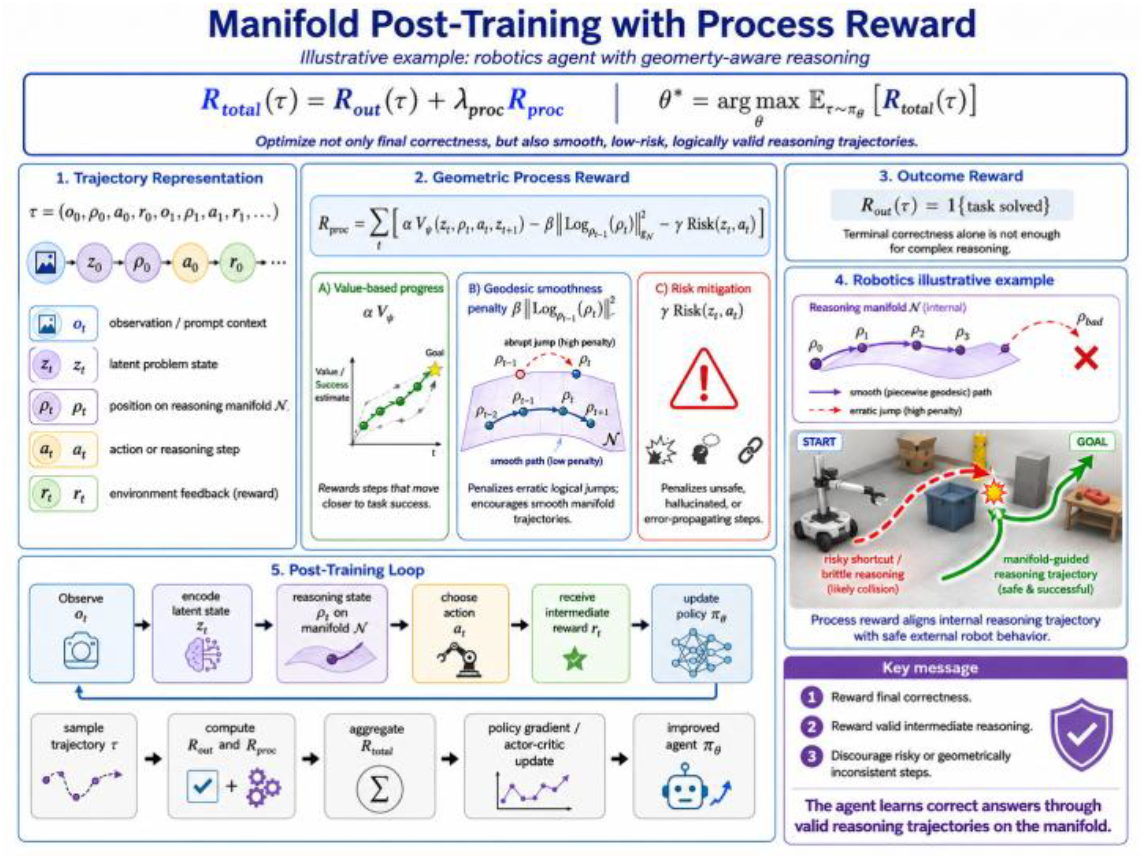
Manifold post-training with process reward.

### 3.12. Geometry-aware memory for POMDP reasoning

Below mathematical formulation describes a Geometry-aware Memory Bank designed for POMDPs. It enables an agent to retrieve relevant knowledge by evaluating how closely the current situation matches past experiences, measured across two distinct geometric manifolds: environment states and internal reasoning tracks.

Here is a breakdown of the mathematical formulation, explaining the components, the retrieval mechanism, and why this geometric approach is highly effective.

#### 3.12.1. The Manifold Memory Bank Structure

The memory bank at time *t* is structured as a set of *N*_*t*_ slots:

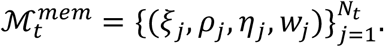

Each slot *j* contains a distinct four-tuple representing a multi-dimensional snapshot of a past experience:

***ξ***_***j***_ ∈ **ℳ (Environment State):** A point lying on a specialized manifold **ℳ** representing a past environment state. Because POMDPs suffer from partial observability, this acts as a geometric anchor for what the environment looked or felt like.

***ρ***_***j***_ ∈ ***N* (Reasoning State):** A point lying on a separate reasoning manifold *N*. This tracks the agent’s internal cognitive state, belief state, or multi-step logic process at that moment.

***η***_***j***_ **(Semantic/Mechanistic Knowledge):** The core payload or value stored in the memory slot. This could be a vector embedding of a policy, environmental transition rules, causal relationships, or abstract concepts learned during that specific episode.

***w***_***j***_ **(Reliability):** A scalar weighting factor indicating how trustworthy, stable, or frequently reinforced this specific memory slot is.

#### 3.12.2. Geometry-Aware Attention Weights (*α*_*j*_)

When an agent encounters a new observation *z*_*t*_ and possesses an internal reasoning state *ρ*_*t*_, it determines how much attention to pay to each past memory slot *j* using a specialized Softmax distribution:

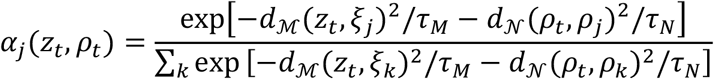

This attention mechanism is governed by two intrinsic geometric properties:

##### Geodesic Distances (*d*_ℳ_, *d*_*N*_)

Instead of standard Euclidean distance (straight lines), *d*_ℳ_ and *d*_*N*_ compute distances along the curves of the manifolds ℳ and *N*. This respects the true underlying geometry of the state spaces (e.g., hyperbolic spaces for hierarchical data or spherical spaces for directional data).

##### Temperature Scaling (*τ*_*M*_, *τ*_*N*_)

These hyper-parameters control the sharpness of the retrieval. A lower temperature forces the agent to focus strictly on the single closest memory, while a higher temperature blends multiple related memories together.

#### 3.12.3. Dual-Track Memory Retrieval Mechanism

The final retrieved memory vector *m*_*t*_ is a weighted linear combination of all stored knowledge payloads, scaled by the attention weights:

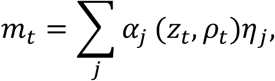

where

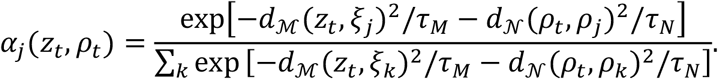

The core strength of this system is that it utilizes a dual-track alignment system to calculate *α*_*j*_. Thus memory is retrieved by both: state similarity and reasoning similarity.

#### 3.12.4. Why This Architecture Matters for POMDPs

In a standard MDP, looking at the environment state is usually enough to choose an action. In a POMDP, the current observation *z*_*t*_ is ambiguous and incomplete.

By utilizing geometry-aware memory:

1. **Disambiguation:** If two different locations look identical (*d*_ℳ_ ≈ 0), the agent can differentiate them based on its internal reasoning track (*d*_*N*_), recalling the memory that aligns with its current plan.
2. **Context-Aware Recall:** The agent avoids retrieving memories that match the physical scenery but are completely irrelevant to its current phase of multi-step reasoning.
3. **Geometric Efficiency:** Operating on manifolds allows the memory bank to capture non-linear relationships and structural hierarchies far more efficiently than standard flat vector databases.

### 3.13. Complete model: Manifold Agentic POMDP

The mathematical formulation of the Manifold Agentic Partially Observable Markov Decision Process (MAP-POMDP) extends traditional POMDPs by separating the environment’s physics from the agent’s internal cognitive reasoning. It maps both components onto continuous, curved geometric surfaces called Riemannian manifolds.

Here is the structured breakdown of the components, transitions, and belief update equations.

#### 3.13.1 Define the Tuple Components

The full model can be defined by the 10-tuple:

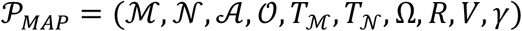

where:

**ℳ: World-state manifold** containing physical environment states *z*_*t*_.

*N***: Reasoning manifold** representing internal cognitive, memory, or thought states *ρ*_*t*_.

***A*: Action space** containing the physical actions *a*_*t*_ available to the agent.

***O*: Observation space** containing the sensory inputs or environment tokens *o*_*t*_.

***T***_**ℳ**_**: World transition function** governing physical state changes *z*_*t*+1_ given actions and reasoning states.

***T***_***N***_**: Internal reasoning transition** updating the thought state *ρ*_*t*_ based on memory and observations.

**Ω: Observation model** defining the probability distribution of observations *o*_*t*_ given physical states.

***R*: Reward function** mapping states, reasoning, and actions to scalar feedback.

***V*: Verifier model** evaluating the validity or logical truth of transitions.

***γ*: Discount factor** weighting immediate versus future rewards (*γ* ∈ [0.1}).

#### 3.13.2. Formulate the Internal Reasoning State

The agent updates its internal thought process *ρ*_*t*_ ∈ *N* on the reasoning manifold:

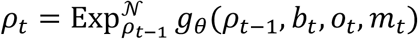

##### Tangent Vector Field

The neural network function *g*_*θ*_ maps the previous reasoning state *ρ*_*t*−1_, current belief state *b*_*t*_, observation *o*_*t*_, and working memory token *m*_*t*_ to a vector in the tangent space 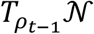.

##### Geometric Projection: The exponential 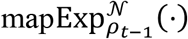 projects this flat tangent vector back onto the curved geometry of the reasoning manifold *N* to yield the updated cognitive state *ρ*_*t*_

#### 3.13.3. Sample the Action Selection

The physical action *a*_*t*_ ∈ *A* executed in the environment is sampled probabilistically:

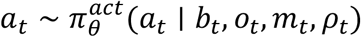

##### Policy Network

The policy function 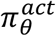 conditions action selection on the environmental variables (*b*_*t*_, *o*_*t*_, *m*_*t*_) wrapped together with the geometric reasoning state *ρ*_*t*_.

#### 3.13.4. Advance the Physical World State

The environment state *z*_*t*+1_ ∈ ℳ updates via a geometric transition mapping:

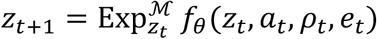

##### Environmental Perturbation: The function *f*_*θ*_ calculates the trajectory direction based on the current state *z*_*t*_, action *a*_*t*_, agent reasoning state *ρ*_*t*_, and external stochastic environment noise *e*_*t*_

##### Manifold Geodesic: The exponential map 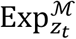 steps along the geodesic path on the world manifold ℳ to determine the exact destination state *z*_*t*+1_

#### 3.13.5. Generate the Next Observation

The environment produces a new observation token \(o_{t+1}\) based on the updated physical state:

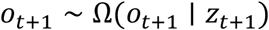

##### Partial Observability: The agent cannot directly read *z*_*t*+1_. It samples *o*_*t*+1_ through the observation probability distribution model Ω

#### 3.13.6. Update the Continuous Belief State

Because the environment is partially observable, the agent maintains a probability distribution over all possible states, updated via manifold integration:

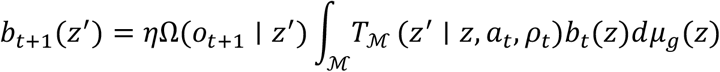

##### Manifold Integration: The integral aggregates the previous belief *b*_*t*_(*z*) across the entire surface of ℳ weighted by the world transition density *T*_ℳ_

##### Volume Element: The term 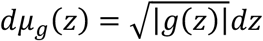 represents the Riemannian volume form dictated by the manifold’s metric tensor *g*

##### Bayesian Correction: The output is multiplied by the observation likelihood Ω(*o*_*t*+1_ ∣ *z*′) and a normalizing constant *η* to ensure the new belief distribution integrates perfectly to 1

In summary, the model coordinates physical space (ℳ) and internal reasoning space (*N*) using Riemannian geometry, swapping standard Euclidean addition for exponential maps (**Exp**) to preserve the structural integrity of curved manifolds during continuous agent updates.

Figure 18 visualize the Complete model for Manifold Agentic POMDP.

**Figure 18.**
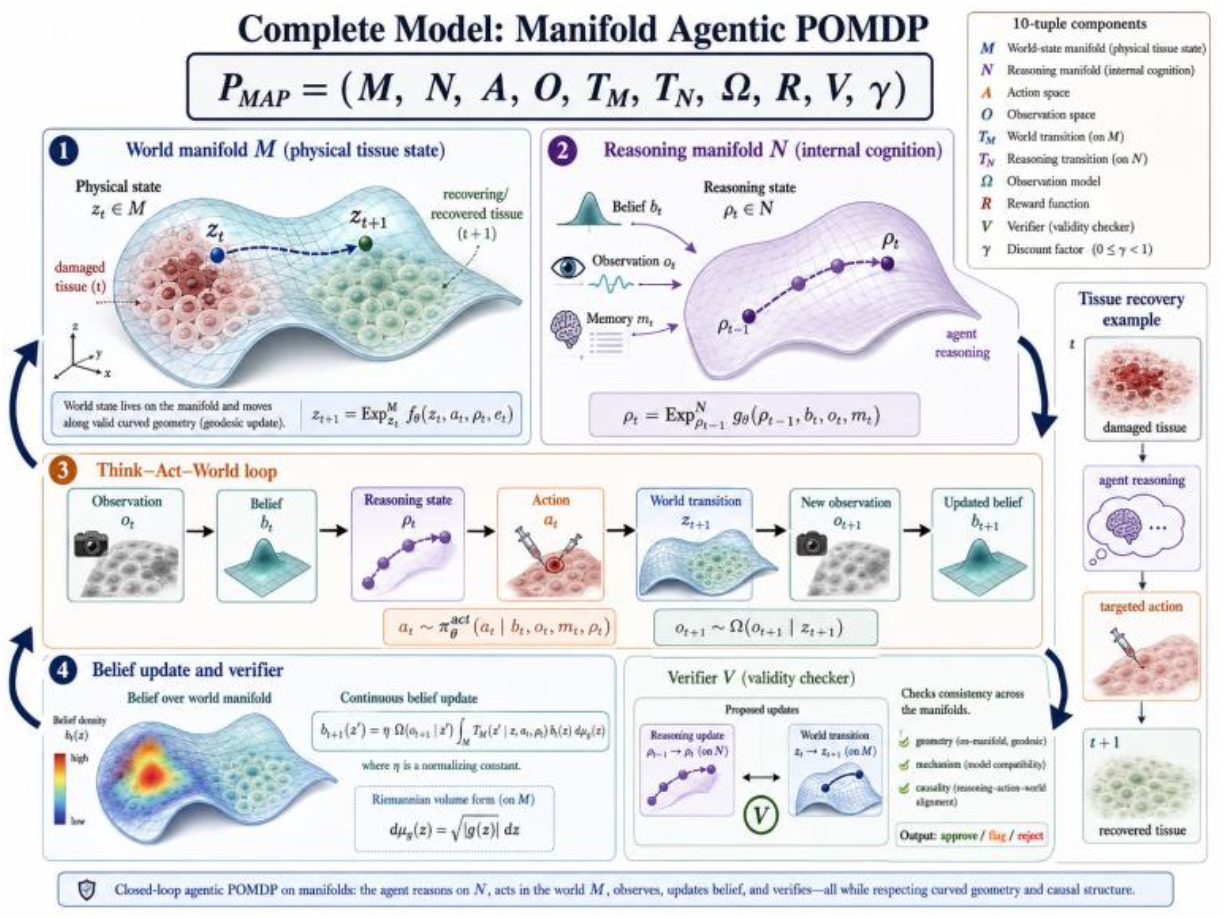
Complete model: Manifold Agentic POMDP.

### 3.14. Learning objective

The primary learning objective of Manifold Agentic POMDP Reasoning (MAP) is to optimize an autonomous agent’s decision-making by embedding its latent reasoning states and policy space onto Riemannian manifolds, ensuring geometric, causal, and mathematical consistency during post-training.

By framing the Partially Observable Markov Decision Process (POMDP) geometrically, this objective ensures that the agent’s thoughts, beliefs, actions, and memories follow continuous trajectories on smooth mathematical surfaces rather than unconstrained Euclidean spaces.

Here is the breakdown of what each component in the total training loss objective balances to achieve this.

The complete training loss is:

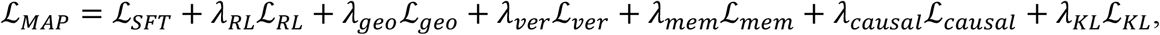

where:

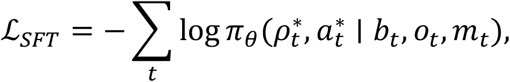

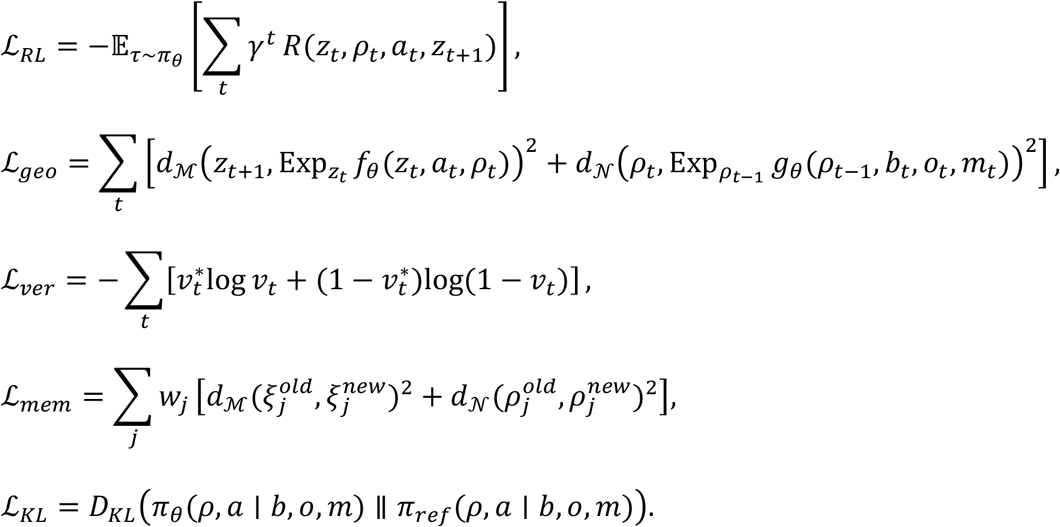

#### 3.14.1. Grounding in Expert Trajectories (*ℒ*_*SFT*_)

##### Supervised Fine-Tuning

This term acts as the behavioral cloning anchor, forcing the agent to maximize the log-likelihood of selecting expert geometric paths 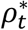 and expert actions 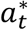.

##### Contextual Inputs

It conditions the policy on the current belief state *b*_*t*_, raw environment observation *o*_*t*_, and memory context *m*_*t*_ to mirror top-tier human or oracle reasoning.

#### 3.14.2. Maximizing Long-Term Environmental Rewards (*ℒ*_*RL*_)

##### Policy Optimization

Reinforcement Learning guides the agent to seek out policies *π*_*θ*_ that maximize expected cumulative discounted rewards.

##### Geometric Transitions

The reward function *R*(*z*_*t*_, *ρ*_*t*_, *a*_*t*_, *z*_*t*+1_) explicitly evaluates how well the agent’s manifold hidden state *z*_*t*_, reasoning trajectory *ρ*_*t*_, and concrete action *a*_*t*_ interact to drive the agent into a favorable next state *z*_*t*+1_.

#### 3.14.3. Preserving Smooth Geodesic Trajectories (*ℒ*_*geo*_)

##### Manifold Exp Map

This enforces physical and mathematical consistency using the Exponential Map (Exp_*p*_*v*), which projects vector steps smoothly along a curved manifold surface.

##### State & Reason Mechanics

The first term minimizes the manifold distance *d*_ℳ_ between the true next latent state *z*_*t*+1_ and the state predicted by the geometric transition function *f*_*θ*_. The second term minimizes the manifold distance *d*_*N*_ to ensure the agent’s internal chain-of-thought progression *ρ*_*t*_ transitions smoothly from its prior thought *ρ*_*t*−1_ based on environmental inputs.

#### 3.14.4. Continuous Self-Verification (*ℒ*_*ver*_)

##### Veridical Calibrations

This binary cross-entropy loss trains an internal critic to output a correctness or verification score *v*_*t*_ ∈ [0,1].

##### Self-Correction

It ensures the agent evaluates its own intermediate logic steps against a ground-truth verifier 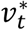, preventing the model from hallucinating or pursuing flawed reasoning paths.

#### 3.14.5. Consolidating Coherent Long-Term Memories (*ℒ*_*mem*_)

##### Memory Preservation

When updating the model, this term prevents catastrophic forgetting of previously learned manifold representations.

##### Geometric Alignment

It penalizes the weighted distance (*w*_*j*_) between old and new memory coordinates (*ξ*_*j*_) and old and new reasoning traces (*ρ*_*j*_), ensuring old knowledge is smoothly warped rather than shattered during optimization.

#### 3.14.6. Ensuring Causal Action Directionality (*ℒ*_*causal*_)

##### Directed Dependencies

Though the explicit definition isn’t expanded in your formula list, this term enforces directed graphical constraints.

##### Anti-Hallucination

It mathematically penalizes spurious correlations, forcing the agent’s actions *a*_*t*_ to be strictly caused by past beliefs and observations, rather than looking ahead or hallucinating non-existent patterns.

#### 3.14.7. Anchoring via Distributional Safety (*ℒ*_*KL*_)

##### Policy Regularization

This calculates the Kullback-Leibler divergence between the active learning policy *π*_*θ*_ and an uncorrupted pre-trained baseline reference policy *π*_*ref*_.

Exploration Control: It keeps the agent’s behavior bounded, preventing the language/reasoning model from degrading into gibberish or collapsing into unsafe execution states while exploring the manifold.

This gives a complete post-training objective for manifold-valued agentic reasoning.

**In summary**, the ultimate goal of the Manifold Agentic POMDP Reasoning objective is to train a resilient, self-verifying, and geometrically constrained agent that navigates partial information by treating the act of “thinking” and “acting” as continuous, mathematically smooth paths along structured geometric shapes (manifolds).

### 3.15. Why this extension is powerful

The original POMDP plus internal reasoning variable is useful because it makes reasoning explicit:

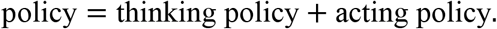

The manifold extension makes it geometrically correct for complex scientific domains:

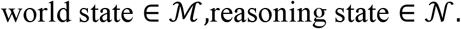

This is important for biological and physical systems because their states often lie on curved, constrained, or multi-chart spaces rather than in flat Euclidean coordinates.

For example:

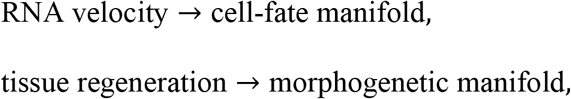

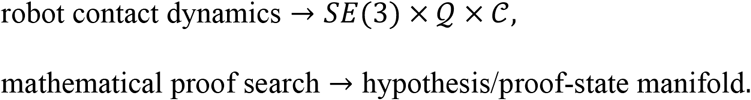

Thus, the agent does not merely generate tokens or actions. It learns a controlled trajectory over:

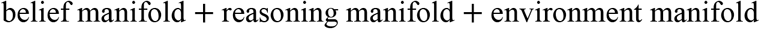

### 3.16 Final compact formulation

The core model is:

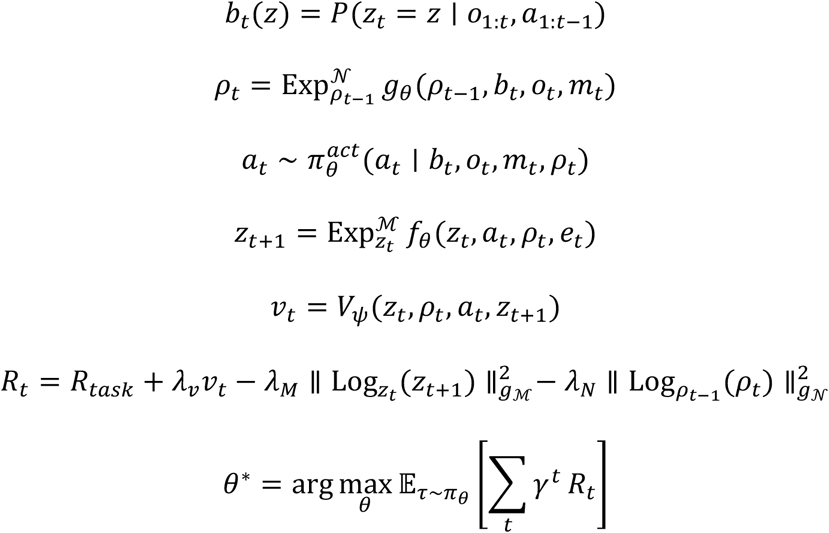

This gives a principled way to extend the paper’s **POMDP + internal reasoning + post-training RL** framework from Euclidean space to manifold space.

## 4. Simulation Result

### 4.1. Curved Tissue Manipulation and Recovery (CTMR)

#### 4.1.1. Core idea

We simulate a tissue sheet as a graph of cells living on a curved manifold. The agent must:

observe the damaged tissue, infer its latent state, propose repair hypotheses, verify them, choose a repair action, and recover the tissue.

This is very similar in spirit to the tissue manipulation / regeneration workflow shown in your full manifold-agent figure.

Figure 19 shows the work flow of Curved Tissue Manipulation and Recovery (CTMR).

**Figure 19.**
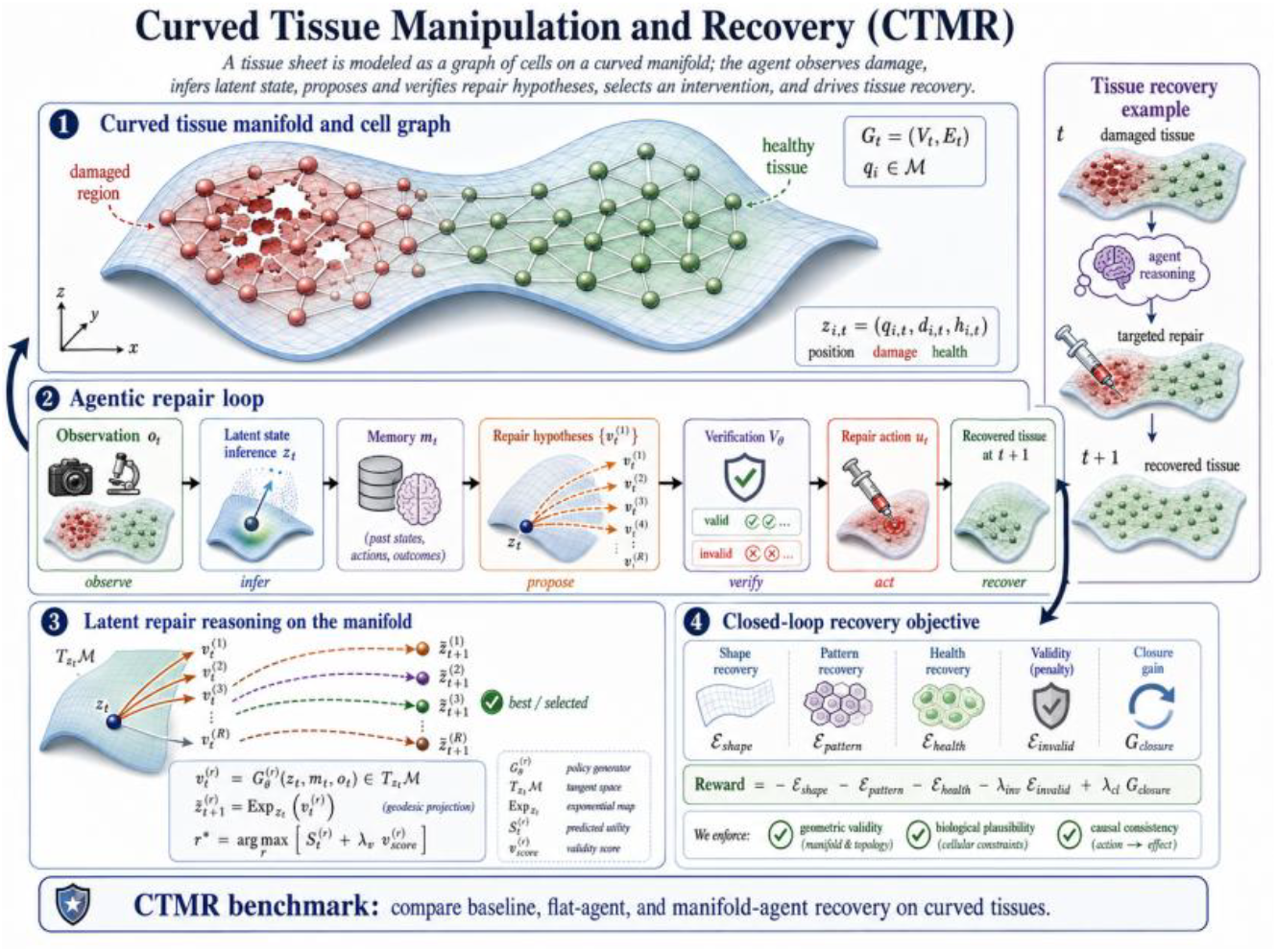
The work flow of Curved Tissue Manipulation and Recovery (CTMR).

### 4.2. Environment definition

#### 4.2.1. Tissue manifold

Let the tissue lie on a smooth 2D manifold embedded in ℝ^3^:

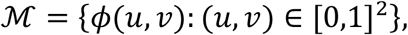

with

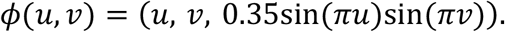

This gives a curved tissue sheet.

You can also make a harder version with stronger folding:

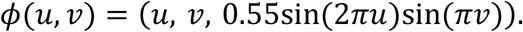

So the tissue is intrinsically 2D, but extrinsically curved in 3D.

#### 4.2.2. Tissue graph

The tissue is represented as a graph:

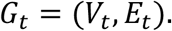

- *V*_*t*_: cells
- *E*_*t*_: local adjacency relations

Use a 10 × 10grid of 100 cells in chart coordinates (*u*_*i*_, *v*_*i*_), then map them to ℳ. Each node *i* has state:

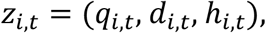

where

- *q*_*i,t*_ ∈ ℳ: cell position on the tissue manifold
- *d*_*i,t*_ ∈ [0,1]: differentiation / gene-expression state

*h*_*i,t*_ ∈ [0,1]: health / viability.

#### 4.2.3. Target healthy tissue

Define a target healthy tissue state:

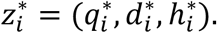

Take:

- 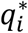: original undamaged manifold position,
- 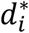: desired developmental gradient, for example

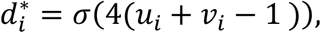

where *σ* is the logistic sigmoid,

- 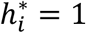: healthy viability.

### 4.3. Damage and manipulation task

#### 4.3.1. Initial damage

At time *t* = 0, damage a local region centered at (*u*_*c*_, *v*_*c*_).

For nodes in the wound region:

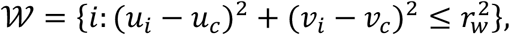

set:

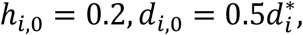

and displace them along the manifold tangent direction:

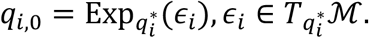

Thus the tissue is damaged both:

- **geometrically**,
- **biologically**.

#### 4.3.2. Agent actions

At each step, the agent chooses a localized intervention:

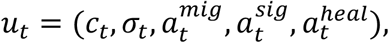

where

- 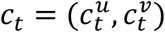: center of intervention in chart coordinates
- *σ*_*t*_: intervention radius
- 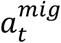: migration cue
- 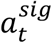: signaling / differentiation cue
- 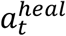: healing / viability cue

These induce a Gaussian spatial kernel:

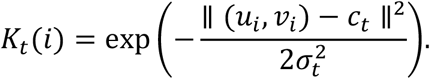

So the agent applies a local repair field.

### 4.4. Tissue dynamics

We update geometry, differentiation, and health.

#### 4.4.1. Geometric interaction

For node *i*, define manifold neighbor transport:

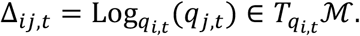

Aggregate neighborhood geometry:

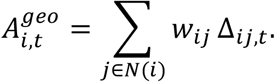

#### 4.4.2. Position update on the manifold

Let 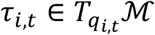 be a local migration direction field induced by the intervention center. Then:

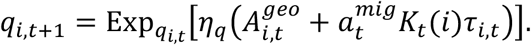

So cells move along the manifold, not through ambient Euclidean shortcuts.

#### 4.4.3. Differentiation / signaling update

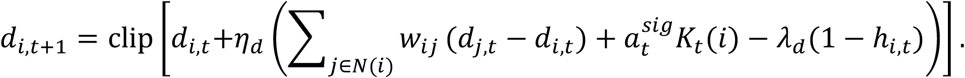

This captures:

- graph diffusion of developmental state,
- intervention-induced signaling,
- damage-related penalty.

#### 4.4.4. Health / viability update

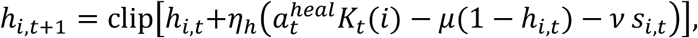

where *s*_*i,t*_ is local stress, e.g.

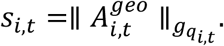

### 4.5. Observation model

The agent does not observe the true state perfectly.

Let observation be:

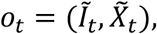

where

- 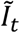: noisy tissue image,
- 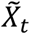: partial node measurements.

For example:

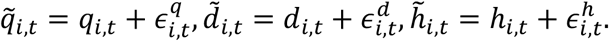

Thus the task is naturally a partially observed reasoning problem.

### 4.6. Algorithms to compare

#### 4.6.1. A simple reactive policy

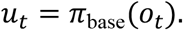

Features:

- no explicit memory *m*_*t*_,
- no hypothesis set 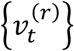,
- no verifier,
- no repair module,
- no manifold geometry.

This could be a shallow MLP or a standard graph policy.

#### 4.6.2. Full flat-agent reasoning algorithm

Use the full reasoning loop, but flatten everything into Euclidean latent space.

**State encoding**

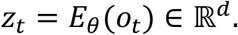

**Memory**

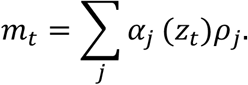

**Hypotheses**

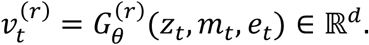

**Projection**

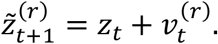

**Scoring, verification, repair**

same pipeline as the full model, but with Euclidean distances and ordinary vector arithmetic.

**Action**

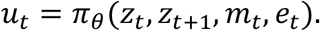

So this model is **reasoning-rich**, but **geometry-wrong**.

#### 4.6.3. Full manifold-aent reasoning algorithm

Use the full model we defined in method section.

**Encode**

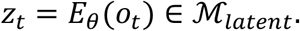

**Memory**

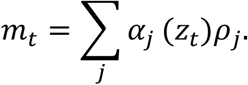

**Candidate hypotheses**

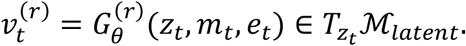

**Project**

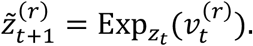

**Score**

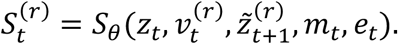

**Verify**

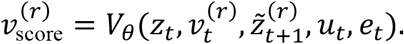

**Commit or repair**

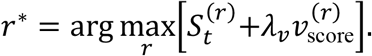

If

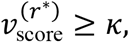

then

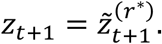

Otherwise,

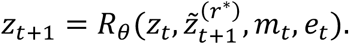

**Action**

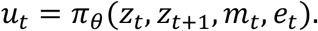

This is the geometry-aware full reasoning algorithm.

### 4.7. Reward and evaluation objective

Define the per-step reward:

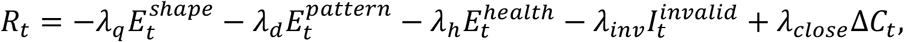

where:

**Shape error**

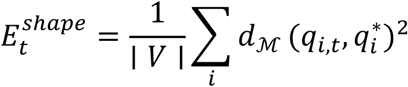

**Pattern error**

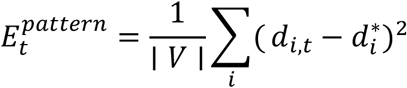

**Health loss**

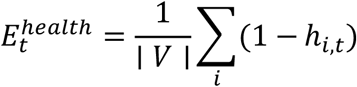

**Invalid transition indicator**

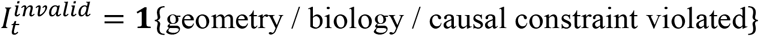

**Wound closure improvement**

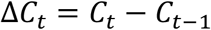

where *C*_*t*_ is fraction of wound region recovered.

### 4.8. Evaluation metrics

Use the following metrics on held-out test episodes:

1. **Recovery success rate**

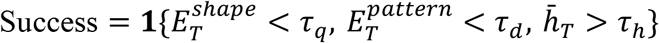
2. **Final geodesic shape error**

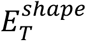
3. **Final pattern MSE**

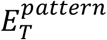
4. **Final mean health**

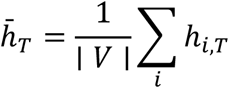
5. **Invalid intervention rate** percentage of proposed transitions failing validity constraints
6. **Repair success rate** among rejected transitions, how many are successfully corrected
7. **Cumulative reward**
8. **Steps to recovery** first time step when recovery criteria are met

### 4.9. Illustrative numerical results

Table 1 summarizes aggregate test performance with mean ±sd across 20 seeds, comparing the

**Table 1.** Aggregate test performance of four models.

| Method | Success (%) | Final geodesic shape error | Final pattern MSE | Final mean health | Invalid transition rate (%) | Repair success (%) | Cumulative reward |
| --- | --- | --- | --- | --- | --- | --- | --- |
| Baseline reasoning | 41.8 ± 4.9 | 0.162 ± 0.014 | 0.081 ± 0.008 | 0.781 ± 0.021 | 18.7 ± 2.8 | N/A | 42.6 ± 5.3 |
| Full flat-agent reasoning | 68.9 ± 4.1 | 0.096 ± 0.010 | 0.047 ± 0.005 | 0.872 ± 0.017 | 10.9 ± 1.9 | 57.4 ± 5.8 | 66.8 ± 4.7 |
| Full manifold-agent reasoning | 89.7 ± 3.2 | 0.044 ± 0.006 | 0.021 ± 0.003 | 0.946 ± 0.011 | 3.2 ± 0.9 | 81.6 ± 4.4 | 87.9 ± 3.9 |
| MAP-R agent | **92.4 ± 2.8** | **0.038 ± 0.005** | **0.018 ± 0.003** | **0.957 ± 0.010** | **2.6 ± 0.8** | **84.9 ± 4.1** | **91.8 ± 3.6** |

Full manifold-agent reasoning against the full flat-agent reasoning and Baseline reasoning.

As Table 1 shown, the MAP-R and Full manifold-agent reasoning consistently improves over against the full flat-agent reasoning and Baseline reasoning under all 7 indies. MAP-R achieves the largest success rate (92.4 ± 2.8, on average), on average, 2.7, 23.5 and 50.6 points are larger than the full manifold-agent reasoning, the full flat-agent reasoning and the Baseline reasoning, respectively. The MAP-R improves, 13.6%, 60.4% and 76.5% over the full flat-agent reasoning and Baseline reasoning under the final geodesic shape error reduction, respectively.

Under the final pattern MSE, the MAP-R improves 14.3%, 61.7% and 77.8% over the full manifold-agent reasoning, the full flat-agent reasoning and Baseline reasoning, respectively. Under invalid transition rate, the MAP-R improves 18.8%, 76.1% and 86.1% over the full manifold-agent reasoning, the full flat-agent reasoning and Baseline reasoning, respectively. Under the final mean health, the MAP-R increase 1.1%, 8.5 % and 17.6% over the full flat-agent reasoning, the full flat-agent reasoning and Baseline reasoning, respectively. The MAP-R achieves the largest repair success rate (on average, 84.9 ± 4.1), on average 4.4 and 27.5 points larger than the full manifold-agent reasoning, and the full flat-agent reasoning, respectively. the MAP-R obtains the largest cumulative reward (on average, 91.8 ± 3.6), on average 3.9, 25 points and 49.2 points larger than the full manifold-agent reasoning, the full flat-agent reasoning and Baseline reasoning, respectively.

## 5. Discussion

We presented two types of manifold-agent reasoning: the MAP-R agent reasoning and the full manifold-agent reasoning as a geometric extension of agentic reasoning from flat Euclidean latent spaces to curved Riemannian state and reasoning spaces. We showed that the performance rank of our types of agents are MAP-R > full manifold-agent reasoning > full flat-agent reasoning > Baseline reasoning. We demonstrated that when the world is curved, the agent should reason on the curve.

The central motivation is that many biological, physical, and embodied systems do not naturally evolve in unconstrained Euclidean coordinates. Tissue morphology, cell-state transitions, developmental trajectories, robotic configurations, and mechanistic reasoning processes often lie on curved, constrained, or locally charted spaces. In such settings, reasoning by ordinary vector addition can produce invalid shortcuts, whereas reasoning by tangent-space generation, exponential-map projection, geodesic distance, verification, and self-repair preserves the intrinsic geometry of the system and are rich in the reasoning.

A key observation from the Curved Tissue Manipulation and Recovery simulations is that flat reasoning can work reasonably well when the underlying state space is nearly flat. This is expected from differential geometry: locally, a smooth manifold can be approximated by its tangent space. When curvature is weak and interventions are small, a Euclidean update may approximate the corresponding manifold update. This explains why flat-agent reasoning can improve substantially over a simple baseline. The full flat-agent still benefits from memory retrieval, candidate hypothesis generation, scoring, verification, repair, and policy selection. Thus, even without explicit manifold geometry, structured agentic reasoning provides a major advantage over a reactive baseline.

However, Euclidean reasoning fails as curvature, folding, and biological constraints become more important. In a curved tissue space, the shortest path in ambient Euclidean coordinates may not be a valid biological or geometric path on the tissue manifold. A flat update of the form *z*_*t*+1_ = *z*_*t*_ + Δ*z*_*t*_ can move through off-manifold regions, cut across folds, ignore local tissue topology, or create physically implausible transitions. This failure becomes especially visible in high-curvature CTMR conditions, where the flat-agent may select interventions that appear efficient in Euclidean coordinates but are invalid with respect to the intrinsic tissue geometry. The result is higher geodesic shape error, increased invalid transition rate, and reduced recovery success.

The full manifold-agent reasoning algorithm addresses this limitation by replacing Euclidean differences and additions with logarithmic and exponential maps. Candidate hypotheses are generated as tangent vectors 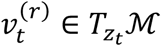, projected back to the manifold through 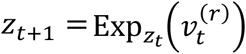, and evaluated using geodesic distances and manifold-aware validity criteria. This construction makes the agent’s reasoning trajectory geometrically admissible by design. The agent does not merely predict a next vector state; it proposes a valid movement direction on the tangent space and then follows the curved structure of the underlying manifold.

The MAP-R formulation further extends the full manifold-agent algorithm by placing it inside a manifold-valued POMDP with internal reasoning variables and reinforcement learning. This distinction is important. The full manifold-agent provides the correct geometric architecture, but MAP-R learns how to use that architecture through interaction, delayed feedback, and policy optimization. In MAP-R, the hidden world state lies on a manifold, the belief state is a probability distribution over manifold states, and the internal reasoning variable evolves on a reasoning manifold. The think-act policy first generates an internal reasoning state and then conditions action selection on that reasoning state. Reinforcement learning then optimizes not only immediate prediction or local validity, but also long-term recovery, cumulative reward, and robustness under partial observability.

This explains why manifold reasoning with reinforcement learning can outperform full manifold reasoning without reinforcement learning. The non-RL manifold-agent may generate geometrically valid transitions, but it is not necessarily optimized for long-horizon intervention strategy. MAP-R can learn which reasoning paths, repair hypotheses, and actions produce the best downstream tissue recovery. It can adapt to noisy observations, hidden damage severity, delayed health improvement, and uncertainty in the tissue state. Thus, MAP-R combines geometric validity with policy improvement, making it especially appropriate for sequential decision problems such as tissue manipulation and recovery.

Graph-Agentic Manifold Reasoning provides an additional advantage for biological systems because tissues are inherently relational. A tissue is not simply a collection of independent points on a manifold. It is a graph of interacting cells, extracellular matrix components, local neighborhoods, signaling relationships, mechanical constraints, and causal dependencies. Standard graph neural networks assume node states lie in flat vector spaces and aggregate messages through Euclidean operations. In contrast, graph-agentic manifold reasoning assigns each node a manifold-valued state (*z*_*it*_ ∈ ℳ), transports neighbor information into the tangent space by 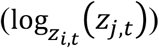, computes attention using biological and contextual edge features, aggregates tangent messages, and updates node states through exponential maps.

This architecture is particularly suitable for tissue recovery because it respects both local cell-cell interactions and global tissue geometry. Edge features can represent biological similarity, technical similarity, metadata compatibility, ligand-receptor relationships, mechanical coupling, or causal relations. The manifold component ensures that cell-state updates remain geometrically valid, while the graph component ensures that recovery decisions depend on local tissue context. This combination is difficult to achieve with purely Euclidean agents or ordinary GNNs.

The stability of manifold reasoning arises from its tangent-space and geodesic structure. Rather than allowing arbitrary jumps in latent space, the agent first computes a local tangent direction and then maps it back to the manifold. This prevents the reasoning trajectory from drifting into invalid regions of the state space. In CTMR, this means that repair trajectories follow the curved tissue surface rather than cutting through the embedding space. More generally, it means that agent reasoning is constrained by the geometry of the domain, which can reduce instability, overshooting, and invalid intermediate states.

Verification and self-repair are also essential. Agentic systems can generate plausible but invalid intermediate steps. In language models, this is often described as hallucination. In manifold reasoning, a related failure occurs when a proposed transition violates geometry, mechanism, causality, or safety constraints. The verifier (*V*_*θ*_) evaluates whether a candidate transition is admissible before the agent commits to it. If the transition fails verification, the self-repair module maps the invalid proposal to a nearby valid state or regenerates a better hypothesis. This creates a verify-before-commit architecture that reduces propagation of invalid reasoning steps. The ablation results support this interpretation. Removing memory weakens long-horizon recovery because the agent loses access to past interventions and outcomes. Removing the verifier increases invalid transitions because candidate hypotheses are no longer filtered.

Removing the repair module prevents the agent from correcting failed proposals. Removing manifold geometry reduces the method to a full flat-agent, which retains reasoning depth but loses geometric validity. These results suggest that the performance of the full system is not due to a single component, but to the interaction between geometry, memory, verification, repair, and policy selection.

The curvature sweep further clarifies the role of geometry. When curvature is low, the difference between flat and manifold reasoning is smaller because Euclidean approximations are locally adequate. As curvature increases, however, the flat-agent degrades while the manifold-agent remains stable. This supports the central claim of the paper: manifold reasoning is most valuable precisely when the underlying data-generating process is curved, folded, or constrained.

There are several limitations. The CTMR benchmark is a controlled toy simulation and does not yet capture the full complexity of real tissue biology, spatial transcriptomics, or regenerative medicine. The exponential and logarithmic maps are simplified in the simulation, whereas real biological manifolds may require learned charts, approximate geodesics, or data-driven metric tensors. The verifier and repair modules are also idealized and would need careful validation in real applications. Finally, reinforcement learning introduces additional design choices, including reward shaping, exploration strategy, and safety constraints.

Despite these limitations, the results suggest that Manifold Agentic Reasoning provides a principled framework for reliable agentic decision-making in curved scientific domains. The framework unifies manifold geometry, POMDP belief updating, internal reasoning variables, graph-agentic message passing, verification-gated commitment, and self-repair. In biological tissue recovery, this allows an agent to reason over damaged tissue as a curved graph-structured system, generate manifold-valid repair hypotheses, reject invalid transitions, and choose interventions that improve recovery. More broadly, the approach may be useful for spatial biology, regenerative medicine, robotics, physical reasoning, and other domains where valid reasoning must respect geometry, mechanism, and causality.

In summary, flat reasoning can succeed when the world is nearly flat, but it becomes unreliable when the true state space is curved. Full manifold-agent reasoning improves stability by using tangent-space hypotheses, geodesic transitions, and verification-gated repair. MAP-R further improves performance by optimizing manifold reasoning policies through reinforcement learning under partial observability. Graph-Agentic Manifold Reasoning extends these benefits to relational biological systems such as tissues. Together, these results support the view that future agentic AI systems for scientific and embodied domains should reason not only over tokens or vectors, but over geometrically valid trajectories on manifolds.

The purpose of this paper is to stimulate discussions about how to develop efficient and powerful AI agents, unify Euclidean and manifold approach and facilitate the rapid development of strong and safe AI systems.

## Acknowledgements

The authors wish to acknowledge the use of AI-powered language models (ChatGPT) for assistance in creating images, improving the grammar, spelling, and readability of this manuscript.

## Author contributions

TX and ZH: Derive formulas and perform data analysis, XS: Problem formulation and formula derivation, LJ: Design project, MX: Design project, perform data analysis and write paper.

